# The impact of biological invasion and genomic local adaptation on the geographical distribution of *Aedes aegypti* in Panama

**DOI:** 10.1101/744607

**Authors:** Kelly L. Bennett, W. Owen McMillan, Jose R. Loaiza.

## Abstract

Local adaptation is an important consideration when predicting arthropod-borne disease risk because it can impact on vector population fitness and persistence. However, the extent that vector populations are adapted to local environmental conditions and whether this can impact on species distributions generally remains unknown. Here we find that the geographic distribution of *Ae. aegypti* across Panama is rapidly changing as a consequence of the recent invasion by its ecological competitor, *Aedes albopictus*. Although *Ae. albopictus* has displaced *Ae. aegypti* in some areas, species coexist across many areas, raising the question: What biological and environmental factors permit population persistence?. Despite low population structure and high gene flow in *Ae. aegypti* across Panama, excepting the province of Bocas del Toro, we identify 128 candidate SNPs, clustered within 17 genes, which show a strong genetic signal of local adaptation. This putatively adaptive variation occurs across relatively fine geographic scales with the composition and frequency of candidate adaptive loci differing between populations in wet tropical environments along the Caribbean coast and the dry tropical conditions typical of the Pacific coast of Panama. Temperature and vegetation were important predictors of adaptive genomic variation in *Ae. aegypti* with potential areas of local adaptation occurring within the Caribbean region of Bocas del Toro, the Pacific coastal areas of Herrera and Panama City and the eastern Azuero Peninsula. Interestingly, several of these locations coincide with areas where *Ae. aegypti* and *Ae. albopictus* co-exist, suggesting that *Ae. aegypti* could have an adaptive edge under local environmental conditions that impacts on inter-specific competition with *Ae. albopictus*. Our results guide future experimental work by suggesting that locally adapted *Ae. aegypti* are able to persist on invasion by *Ae. albopictus* and, as a consequence, may fundamentally alter future arborviral disease risk and efforts to control mosquito populations.

**Author Summary:** Local environmental adaptation of mosquito vectors can alter the landscape of arthropod-borne disease by impacting on life history traits that increase their relative fitness thus promoting population persistence. We have identified a number of genomic loci in *Ae. aegypti* from Panama that exhibit a signal of natural selection associated with variation in the environment. Loci with a signal of local adaptation are predominately partitioned between wet and dry tropical environments with variation largely impacted by temperature and vegetation indices. Local adaptation in tandem with changes in the geographic distribution of *Ae. aegypti* due to the recent invasion of its ecological competitor, *Ae. albopictus,* has the potential to alter the landscape of arborviral disease.

## Introduction

The establishment and persistence of vectors within new geographic locations poses a serious threat from emerging and endemic arboviral diseases [1,2]. For example, shifts in the distribution of ticks and *Culex* mosquitoes are linked to the rise of West Nile Virus and tick-borne encephalitis viruses within North America [3–5]. In addition, the introduction of invasive *Aedes* mosquitoes has facilitated the recent spread of Zika and Chikungunya viruses throughout the Americas [6,7]. Although introduced vector populations are unlikely to be at their fitness optimum when first confronted with a new environment, local adaptation may play a large role in disease dynamics as vectors adapt to their environment, increase their relative fitness and acquire new traits, thus potentially increasing the threat of human arboviruses. However, local environmental adaptation has not yet been characterised for any *Aedes* mosquito.

The importance of adaptation for human disease is exemplified in *Aedes aegypti*’s evolution to human commensalism and the establishment of a number of arboviruses worldwide [8]. This mosquito has undergone behavioural and genetic changes in comparison to its ancestral African form, including the evolution of house-entering behaviour and a preference for human odour and blood-feeding [9–11]. The adaptation of *Ae. aegypti* to exploit human environments has allowed for the spread of zoonotic arboviral diseases from forest animals to humans and promoted invasiveness through human-assisted dispersal [8]. Another human commensal, *Aedes albopictus*, is similar to *Ae. aegypti* across many ecological axes. The tiger mosquito has expanded from Asia within the last ~40 years and is now also globally distributed [12]. In many locations, *Ae. albopictus* has displaced resident *Ae. aegypti* [13,14], but the factors that facilitate co-occurrence are still unclear [15]. Identifying the abiotic and biotic factors important in *Aedes* species interactions, particularly whether the two *Aedes* mosquitoes coexist is critical. These interactions are likely to fundamentally reshape the arboviral disease landscape worldwide.

Here we characterize genome-wide variation in *Ae. aegypti* across Panama and use this data to explore the interplay between invasion history, the potential for local adaptation, and ecological change. Panama provides an ideal opportunity to begin to understand how these factors interact and, ultimately, affect the disease landscape by impacting on *Aedes* species distributions. Panama is a small country, measuring just 772 kilometres East to West and 185 km North to South, but provides a wealth of contrasting climatic conditions and discrete environments. This is largely owing to its situation as a narrow isthmus flanked by the Caribbean Sea and Pacific Ocean as well as the Cordillera Central mountain range, which acts as a North-South divide. Panama is also a hub of international shipping trade, providing an important route of *Aedes* mosquito invasion into the Americas. Panama’s worldwide connections have potentially facilitated multiple introductions of the invasive *Ae. aegypti* mosquito dating back to the 18^th^ century in association with the global shipping trade [8,16,17]. In addition, the Pan-American highway bisects the country, stretches almost 48,000km throughout mainland America and provides important conduit for human-assisted dispersal of *Aedes* mosquitoes [13,18].

We first investigate how genomic variation in *Ae. aegypti* is distributed across Panama. Secondly, we evaluate the historical and current geographic distributions of both this mosquito and *Ae. albopictus*. *Aedes albopictus* was first documented in Panama in 2002, providing the opportunity to study how the interactions between the two species play out across a heterogeneous landscape. Finally, we investigate whether local environmental adaptation could play a role in *Aedes* population dynamics by identifying loci with a genomic signal of local adaptation that are associated with discrete environmental conditions. These genomic regions might allow *Ae. aegypti* populations to persist in competition with invading *Ae. albopictus*. How this scenario plays out in Panama will provide insight into global species interactions and the spatial heterogeneity of viral transmission.

## Results

### Characterisation of sequence variation in *Ae. aegypti*

We processed 70 *Ae. aegypti* individuals with hybridisation capture-based enrichment from 14 localities widespread across Panama. An average number of 27,351,514 reads were mapped to the genome for each individual with 62 % of these targeted to the designed capture regions. The mean coverage depth per individual was approximately 74X. After applying stringent quality filters, 371,307 SNP’s were identified throughout all captured regions for downstream analyses.

#### Global and local population structure of *Ae. aegypti*

Our large SNP dataset allowed us to examine population structure across both global and local scales. Comparison of global population structure was achieved by comparing a subset of 2,630 of our SNP’s from Panama that were shared with a previously acquired *Ae. aegypti* SNP dataset from 26 other countries worldwide [19–23].

FastStructure analysis revealed that the number of model components and model maximum likelihood were maximised by assigning each individual to between K=4-6 populations (S1 Fig). Similar to that reported previously, we found that the new world variation is composed of an admixture of populations distinct from African and Asian sources at higher values of K [19,20] (S1 Fig). Individuals from Panama, Costa Rica, Colombia, the Caribbean islands and populations from Arizona and Texas in South western USA were consistently composed of a similar composition throughout each possible value of K (S1 Fig). Thus, *Ae. aegypti* from Panama were genetically similar to those found throughout the Americas, consistent with a strong geographic component to the distribution of genetic variation across the world [24].

Within Panama, the much larger dataset including all 371,307 SNP’s, highlighted significant population structure. There were two major genomic clusters (Fig 1B & 1C) that distinguished individuals from Bocas del Toro province in the western Caribbean region compared to individuals from all other regions across Panama, revealed on both FastStructure and PCA analysis of all SNP’s. In addition, *Ae. aegypti* from the eastern Azuero Peninsula also appeared somewhat genetically discrete (Fig 1C). All areas of Panama, including sampling locations on the Azuero Peninsula had similar levels of heterozygosity and therefore the population differences we observed are not expected to result from a recent population bottleneck or from insecticide spraying treatment, which is irregularly applied during epidemics to target adults only within the urban areas of Panama (Fig 1B).

**Fig 1.**
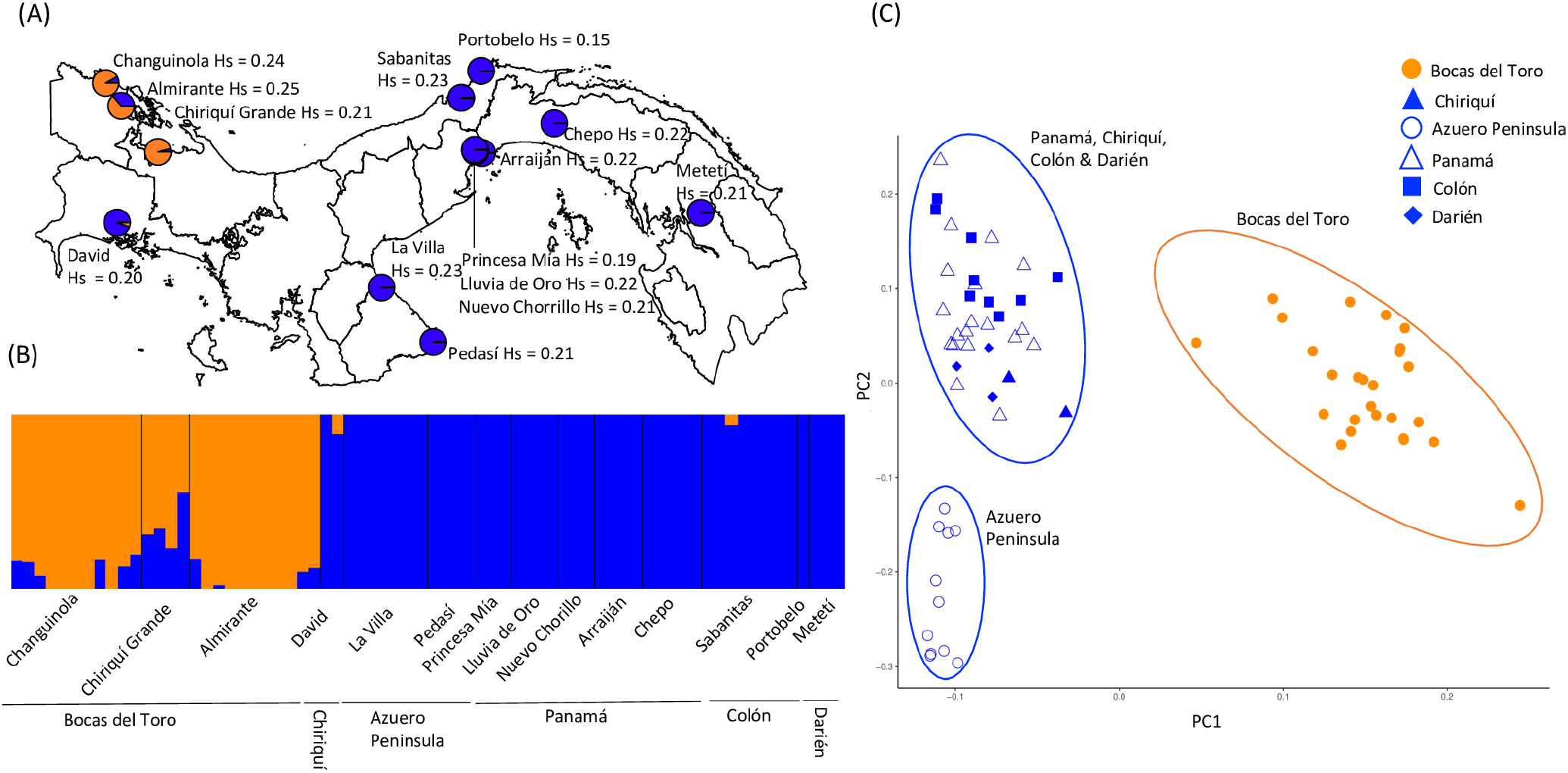
Strong local population structure within the context of regional homogeneity and global population structure: (A) FastStructure plot of K=6 populations comparing 2,630 SNP’s in individuals of *Ae. aegypti* from Bocas del Toro and the rest of Panama to genetically similar populations originating from South-western USA, Caribbean islands, Costa Rica and Columbia. FastStructure assigns each individual to one or more K populations, as indicated by its colour. Genetically similar populations share the same colour or similar admixture composition on comparison. (B) Admixture proportions of K=2 populations in relation to sampling locations and population heterozygosity Hs of *Ae. aegypti* across Panama as determined by FastStructure for 371,307 SNP’s. (C) PCA of all *Ae. aegypti* SNP’s grouped by region.

### The geographical distribution of *Ae. aegypti* in response to invasion by *Ae. albopictus*

To understand how the recent introduction of *Ae. albopictus* has shaped populations of *Ae. aegypti* across Panama over the last decade, we coupled historical surveys of mosquito populations with intensive sampling of focal populations over the last three years. Over the sampling period, there has been significant changes in the geographic distribution of *Ae. aegypti* (Fig 2). Analysis of all occurrence data throughout all years revealed that the presence of *Ae. aegypti* is positively and significantly associated with the presence of *Ae. albopictus* (GLM, Z = 18.93, *d.f* = 7390, P = 0.000), reflecting the ecological similarity of the two species and the continued expansion of *Ae. albopictus* throughout much of *Ae. aegypti*’s historical range. Although both species now co-exist in many areas throughout Panama, areas in the wet and humid western Azuero Peninsula, rural Chiriquí, Veraguas and the province of Panamá outside of Panama City (Gamboa and Chilibre), were solely inhabited by *Ae. albopictus*. This includes regions, from which *Ae. aegypti* was previously documented by the health authorities, confirming that *Ae. albopictus* has indeed replaced *Ae. aegypti* in these areas. The replacement of *Ae. aegypti* by *Ae. albopictus* was further supported by a general decrease in the proportion of positive sampling sites. This proportion has decreased for *Ae. aegypti* since 2005 from ~50 % to ~20 %, while the presence of *Ae. albopictus* has increased from 0 to ~65 % (S2 Fig). *Ae. aegypti* continued to be found in high abundance in Bocas del Toro and Darién, where *Ae. albopictus* has only recently arrived (Darién) or has not yet been documented (Bocas del Toro).

**Fig 2.**
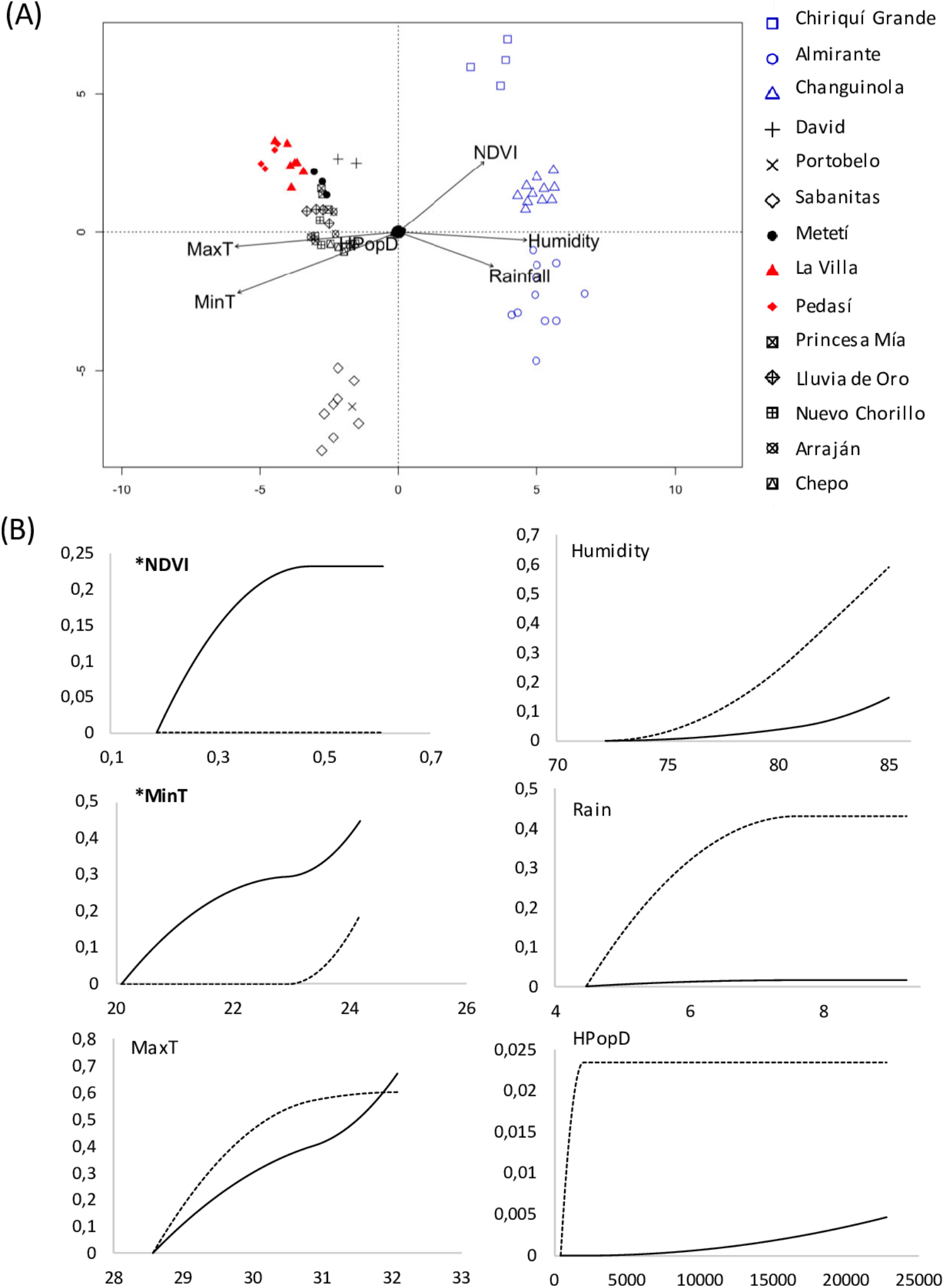
The species replacement and co-occurrence of *Aedes* species is condition dependent: (A) The presence of *Ae*. *aegypti* (orange), *Ae. albopictus* (blue) and species co-occurrence (yellow) recorded by extensive sampling with both active surveillance and oviposition traps during the wet season months from 2016 through to 2018 in comparison to (B) Species occurrence data recorded from 2005 through 2017 through active surveillance by the Ministry of Health in Panama.

### Genomic evidence for local adaptation in *Ae. aegypti* in response to environmental heterogeneity across Panama

The spatial environmental heterogeneity of Panama coupled with the recent population changes associated with the introduction of *Ae. albopictus* provides a framework to ask if there was any evidence that local adaption of *Ae. aegypti* might allow population persistence. If so, we would expect populations of *Ae. aegypti* to harbour genomic loci with a signal of selection that are correlated to the local environmental conditions. These loci are expected to be present in regions of *Aedes* co-existence.

As a first step, we applied redundancy analysis (RDA) to jointly identify candidate outlier loci and to assess how candidate variation was partitioned among the different environmental variables. In this analysis, we tested a number of environmental variables including Normalized Difference Vegetation Index (NDVI), average rainfall, average humidity, average minimum and maximum temperature, and human population density. RDA identified 1,154 candidate SNP’s with a genomic signal of local adaptation, which we used to visualise putatively adaptive variation on ordination plots. Overall, there was a partitioning of alleles dependant on dry tropical and wet tropical conditions. For example, the position of sampled individuals on the RDA ordination plots, in relation to the depicted environmental variables, revealed that the candidate genotypes of *Ae. aegypti* from the wet tropical regions of Almirante and Changuinola in Bocas del Toro province were positively associated with humidity and average rainfall. Those from the wet tropical region of Chiriquí Grande in Bocas del Toro were also positively associated with increasing NDVI vegetation index and negatively associated with higher temperatures (Fig 3A). In comparison, the candidate genotypes of individuals from dry tropical regions of Panamá province (i.e., Princesa Mía, Lluvia de Oro, Nuevo Chorrillo), Los Santos (i.e., La Villa de Los Santos, Pedasí), Darién (i.e., Metetí) and David in Chiriquí province were somewhat positively influenced by both temperature variables and negatively associated with wet and vegetated conditions. Putatively adaptive variation in individuals from the province of Colón (i.e., Sabanitas and Portobelo), locations which receive high rainfall but higher temperatures and lower vegetation cover than in Bocas del Toro province, were associated with intermediate temperature and vegetation conditions.

**Fig 3.**
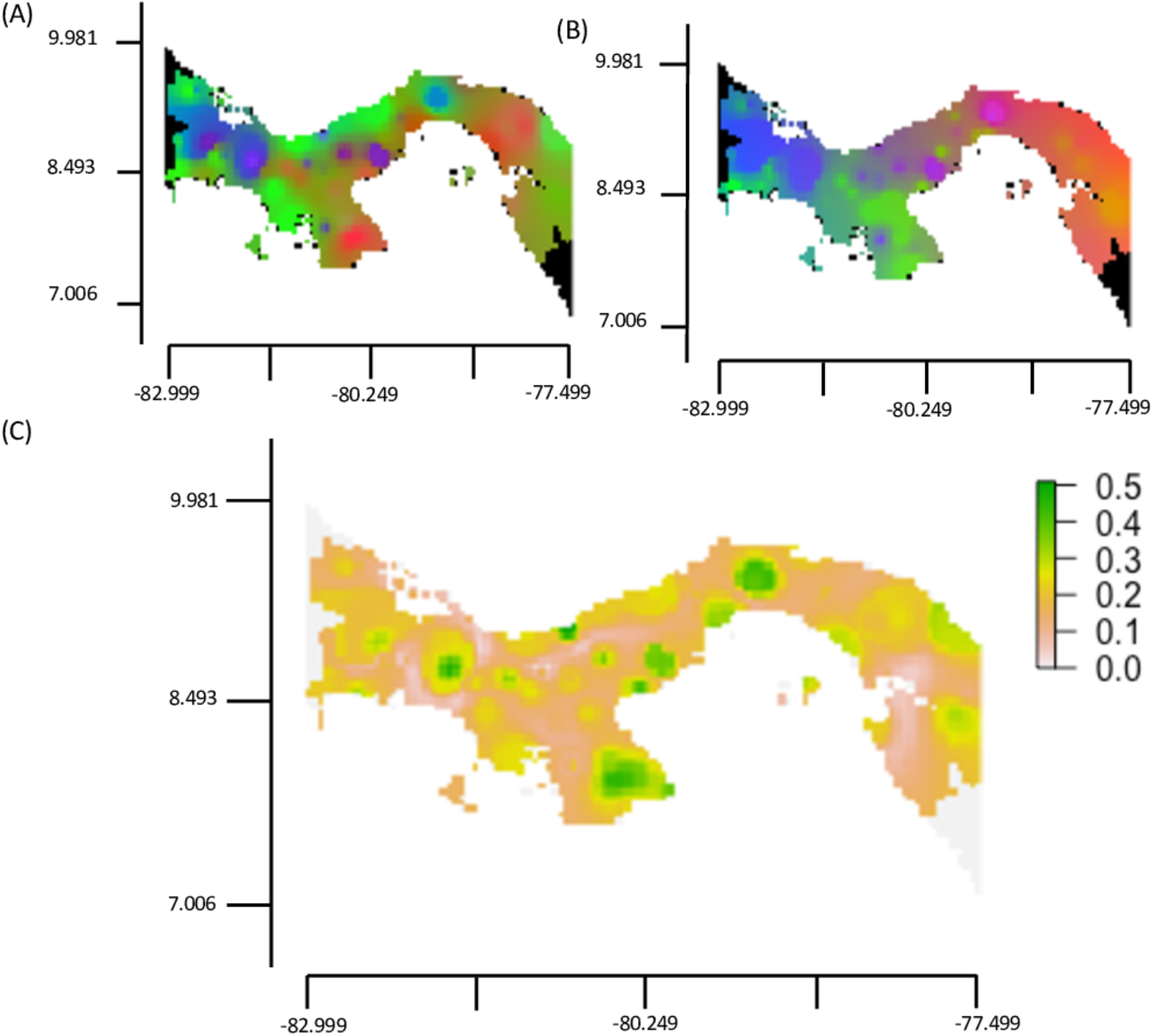
Putative adaptive variation in *Ae. aegypti* is partitioned between wet and dry tropical environments and associated with temperature and vegetation indices: (A) Ordination triplot of the first two constrained ordination axes of the redundancy analysis representing SNP’s either positively or negatively associated with the environmental variables as depicted by the position of the arrows. *Ae. aegypti* from the wettest region (blue) and driest region (red) are highlighted. (B) Compositional turnover splines for GDM analysis for the reference loci that are putatively neutral (dashed line) and the 128 candidate loci with a signal of local adaptation (black line) in association with NDVI vegetation index (NDVI), average minimum temperature (MinT), average maximum temperature (MaxT), average humidity (Humidity), average rainfall (Rain) and human population density (HPopD). A change in allele frequency relative to the reference loci is seen in the putatively adaptive alleles with increasing values of NDVI and MinT, marked in bold with an asterix.

RDA is robust in detecting adaptive processes that result from weak, multilocus effects across a range of demographic scenarios and sampling designs [25]. However, a proportion of the 1,154 candidate loci identified through this single analysis were likely false positives. Thus, rather than reflecting local adaptation, the strongly skewed frequency differences could be reflective of demographic processes such as hierarchical population structure, isolation by distance, allele surfing on range expansion and background selection, as well as, coincidental associations of allele frequencies to environmental variation or even covariance to other environmental factors not included in the analysis [26]. To further refine our identification of putatively adaptive loci, we identified candidates using two additional methods, PCAdapt and Latent Factor Mixed Model analysis (LFMM). Both are considered less sensitive to confounding demography due to their ability to account for population structure or unobserved spatial autocorrelation in the data [27]. The three methods identified different numbers of putatively adaptive loci. For example, compared to the 1,154 outlier SNP’s identified by RDA, PCAdapt identified 352 SNP’s (S3 Fig), whereas LFMM analysis identified 3,426 outlier SNP’s with a signature of selection widespread across the genome and associated with the environment respectively (S4 Fig).

Across all three methods there were 128 SNP’s consistently identified as outliers, providing greater confidence that these loci are located in or close to genomic regions possibly involved in local adaptation. These candidate SNPs fell into 15 distinct clusters, suggesting that linkage disequilibrium was driving some of the observed patterns (S5 Fig). The 128 SNPs fell into 17 genes, 11 of which are annotated as involved in structural functions, enzyme activity and metabolism (S1 Table). None of these genes are known to be involved in the development of insecticide resistance in populations of *Aedes* mosquitoes.

We further narrowed down which of the environmental variables contributed most to the partitioning of genomic variation using a combination of Generalised Dissimilarity Modelling (GDM) and Gradient Forests (GF) analyses. Both approaches allowed us to visualize the allelic turnover of these putatively adaptive loci in relation to each environmental variable. The environmental variables that contributed the greatest variance to both GDM and GF models on analysis of the 128 candidate loci were minimum and maximum temperature (S2 Table, S6 Fig). GDM analysis revealed that an increase in average minimum temperature accompanied a large change in putatively adaptive allele frequencies, visualised as a smooth curve accumulating in a steeper incline at the higher temperature range (Fig 3B). In comparison, GF turnover plots show a steeper incline at the mid-range for both average minimum and maximum temperature (S7 & S8 Fig). GDM analysis also revealed a distinct frequency change in putatively adaptive alleles with increasing NDVI vegetation index, although the change in allele frequency was relatively minor compared to that of minimum temperature (Fig 3B). In comparison, a low to negligible difference in allele frequency was observed in association with average rainfall, average humidity and human population density. Therefore, the variation in putatively adaptive allele frequencies between populations from dry tropical and wet tropical environments of Panama appears largely driven by differences in temperature and NDVI vegetation index.

### The geographic distribution of candidate adaptive alleles in relation to the present distribution of *Ae. albopictus*

Across our 128 candidate SNP’s, we used GDM and GF analysis to visualise the change in frequencies across Panama, and therefore the geographical landscape features which increase or decrease the genomic signature of local adaptation in relation to the environment. GDM analysis presented a smoother turnover in the geographical distribution of putatively adaptive loci than that of putatively neutral loci as indicated by a smoother transition in the colour palette between proximal geographic locations (Fig 4A & 4B). For example, there was similarity in the colouring and therefore allele composition between wet tropical regions along the Caribbean coast (i.e., the mainland/islands of Bocas del Toro, Chiriquí, and both the inland and Caribbean coastal regions stretching from Bocas del Toro through Veraguas to Colón). Similarly, there was greater continuity between dry tropical areas including David in Chiriquí, the eastern Azuero Peninsula (i.e., La Villa de Los Santos and Pedasí), the Pacific coastal regions stretching from the Azuero Peninsula through Coclé to Panamá, and the Darién (i.e., Metetí), indicating that these environments share putatively adaptive alleles. Patterns in the data were less distinct for GF analysis but the geographical distribution of putatively adaptive variation agreed with the GDM analysis in that there was a continuity in the allele composition between the eastern Azuero Peninsula and dry tropical Pacific coastal regions, distinct from the wet tropical regions along the Caribbean coast (S9 Fig).

**Fig 4.**
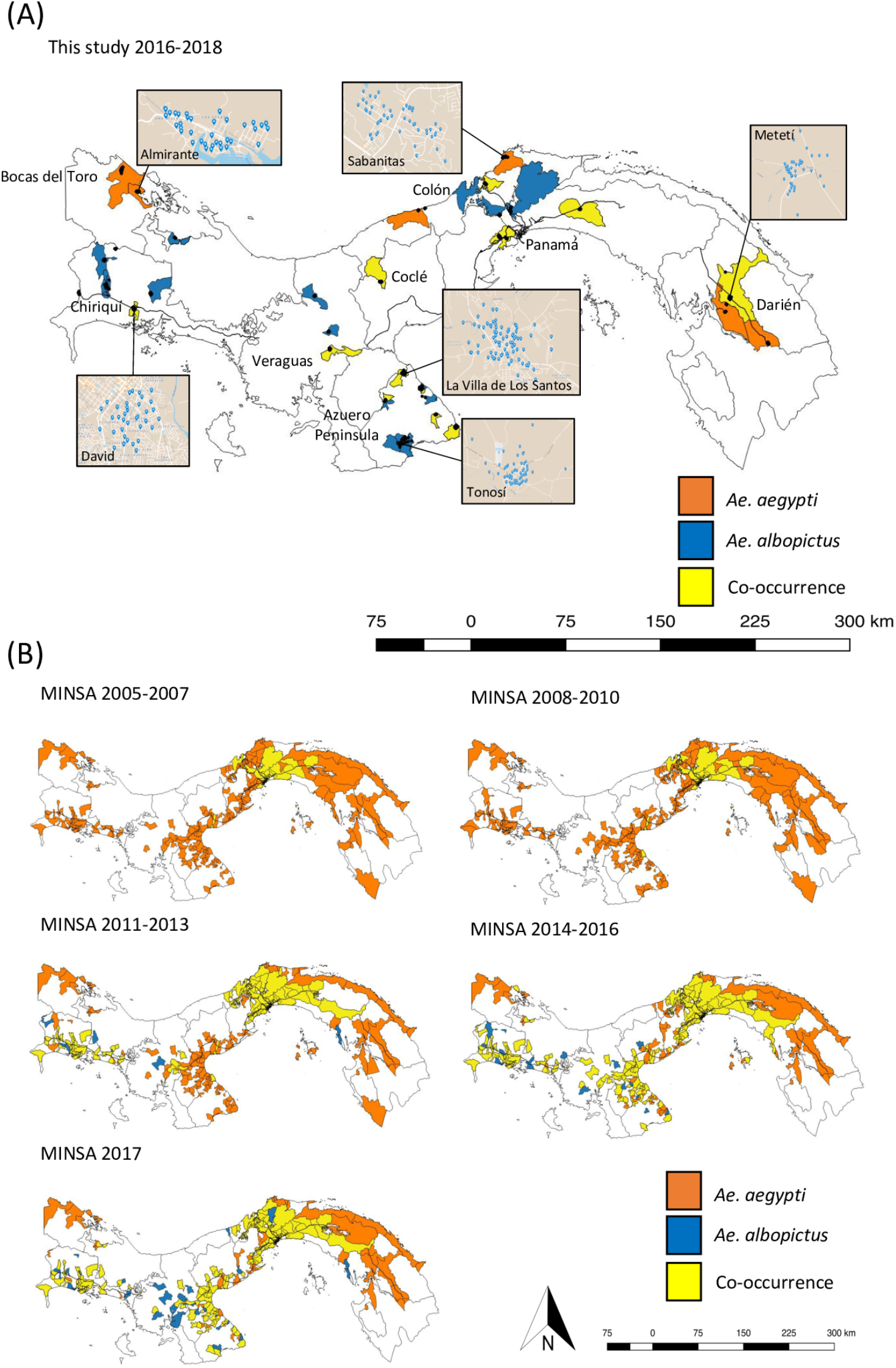
Patches of local adaptation are revealed on comparison of putative neutral and adaptive variation across geographical space: RGB maps of compositional allele frequency turner over across geographical space based on GDM analysis of (A) putatively neutral loci, (B) the 128 candidate loci with a signal of local adaptation and (C) the difference in allele compositional turnover between the putatively neutral reference loci and putatively adaptive candidate dataset using a Procrustes superimposition on the PCA ordinations. On maps (A) and (B), the dissimilarity between allele composition is depicted by an increasing divergent colour spectrum. Locations with a similar allele composition are a similar colour. On map (C), the scale represents the distance between the allele compositional turnover of the reference and candidate SNP datasets, with higher distances indicating areas that are potentially experiencing local adaptation.

Allele frequency turnover as predicted under neutral conditions and a scenario of local adaptation involving the candidate loci were compared across geographical space to identify locations that show the greatest disparity. These reflected the populations within Panama expected to be experiencing a strong genomic signal of local adaptation. Their comparison revealed multiple patches of potential local adaptation widespread across Panama, with a palpable patch occurring in the Azuero Peninsula, as indicated by a high distance between the patterns of predicted compositional allele frequency turnover (Fig 4C). A genomic signal of local adaptation was not identified in the region of Bocas del Toro. Since this region has a strong population structure and distinct climate within Panama, it is likely that the co-correlation of population structure and environmental variation across our sampling design hindered the inference of possible local adaptation in this case. This conclusion was supported by FastStructure analysis of the 128 putatively adaptive loci, which revealed that *Ae. aegypti* from the wet tropical region Bocas del Toro has a distinct allele composition composed of alleles assigned to a distinct composition of K populations, including unique alleles in addition to those shared broadly across the dry tropical regions of Panama (S10 Fig). Although the Talamanca mountain range was documented as a natural geographical barrier to dispersal across the region of Bocas del Toro for some *Anopheles* mosquitoes [28], this was not expected to hinder gene flow in *Ae. aegypti*, since human-assisted movement of this mosquito occurs via the local transport network [18]. Partitioning of the genomic data into 6=K populations revealed that Sabanitas on the Caribbean coast, which is subject to intermediate climate conditions, shared some of the distinct alleles present in Bocas del Toro. Moreover, individuals from the Azuero Peninsula, the driest and least vegetated region of Panama, were also somewhat distinct from other sampled regions since they had reduced levels of admixture.

Comparison of the geographical distribution of putatively locally adapted *Ae. aegypti* as revealed by GDM analysis and the species distribution data revealed that both *Ae. aegypti* and *Ae. albopictus* tended to co-occur in regions where *Ae. aegypti* have divergent candidate loci, despite evidence for species replacement elsewhere (Fig 5). Notably long-term co-existence was documented with the Pacific regions of Panama City, Coclé, the eastern Azuero Peninsula and potentially David in Pacific Chiriquí, where patches of local adaptation in *Ae. aegypti* were identified.

**Fig 5.**
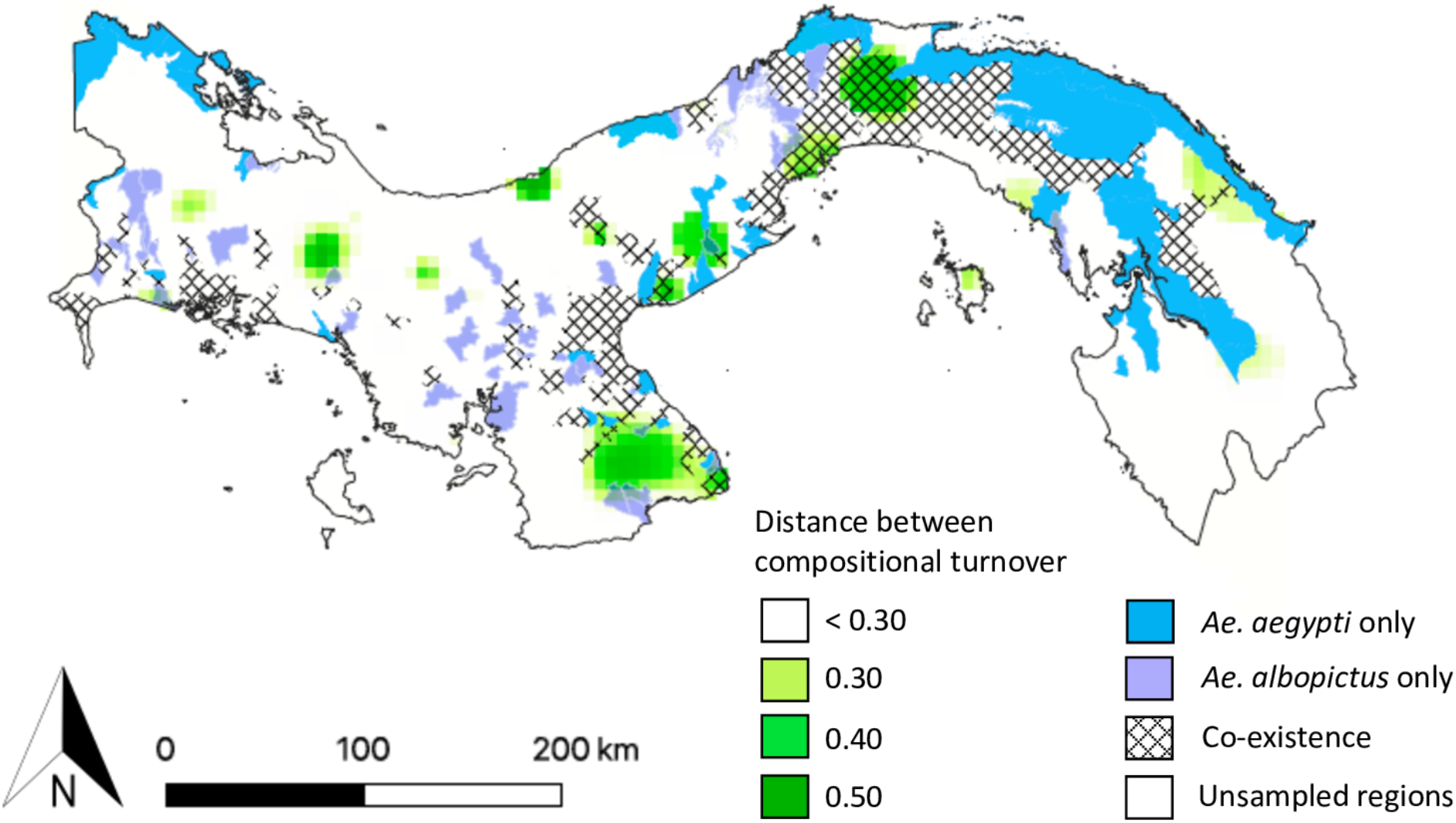
Putatively adaptive loci are predicted in *Ae. aegypti* populations within areas of species co-existence. Corregimientos with *Aedes* co-occurrence (dashed areas) are shown based on the most recent species distributions recorded during this study in 2018 and in other sampled regions by MINSA in 2017. The co-occurrence data is overlaid onto the compositional turnover of the reference and candidate SNP dataset from Figure 4., with values greater than 0.30 shown. Green coloured areas represent regions with a greater predicted distance between the allele composition of the reference and candidate datasets, indicating the potential presence of locally adapted *Ae. aegypti*. Corregimientos where only *Ae. aegypti* (blue) and *Ae. albopictus* (purple) were present are also indicated.

## Discussion

We combined genomic and ecological data to investigate whether *Ae. aegypti* have a signal of local adaptation to the environment, and to investigate whether this variation could influence species persistence on invasion by the recently introduced competitor *Ae. albopictus*. We first documented how fine-scale genomic variation within *Ae. aegypti* is distributed across a complex environment [11]. On a regional scale, Panamanian populations of *Ae. aegypti* are genetically similar to other Central and Caribbean American populations highlighting high dispersal potential and recent gene flow in this invasive species; however, this similarity belies a more complex local genomic architecture. Across Panama, genomic variation was not structured randomly, with the isolated Bocas del Toro region showing significant overall population differentiation. Across the rest of Panama, populations are more homogeneous suggesting high levels of gene flow, likely facilitated by the dispersal of *Aedes* mosquitoes in used tyres that are traded along the Pan-American highway [18]. Nonetheless, a subset of genomic variation was differentially distributed with evidence of localised adaptation across a relatively small number of SNPs and over a relatively fine geographical scale. Genomic variation in these SNPs was strongly correlated with temperature and NDVI vegetation index. Both these abiotic variables were previously identified as important in predicting large-scale *Aedes* distribution patterns [12]. Temperature is important for egg laying, development and survival of *Ae. aegypti* in larval habitats [29] and likely to promote selection to thermal tolerance at the adult stage to resist diurnal and inter-seasonal variation [30]. Vegetation is considered an important variable that contributes to oviposition cues [31], feeding dynamics [32] and microhabitat characteristics such as local moisture supply and shade [33,34]. Although correlational, the genomic patterns raise an important question: Is population persistence in the face of an ongoing invasion by *Ae. albopictus* the result of local adaptation?

The possibility of climatically adapted populations of *Ae. aegypti* is not without precedence. Data on a wide range of organisms with varying dispersal abilities [35–41] demonstrate that even well-connected populations can adapt to environmental differences and habitat heterogeneity across narrow spatial scales. Similar to other landscape genomics studies on plants [42–44], insects [45] and vertebrates [46], we have found a signal of local environmental adaptation across a small number of loci. The inability to identify more putative regions under selection may be the result of the analytical difficulties weak multilocus signatures from the genomic differentiation introduced by genetic drift and demography [25,47]. However, selection on just a few loci with large effects is expected when migration is high since large effect loci are better able to resist the homogenising effects of gene flow [48]. These few regions are expected to have a strong impact on fitness in one environment over the other because the allele with the highest fitness is expected to spread to all populations if this condition is not met [48].

The pattern of recent population distribution change in *Ae. aegypti* in response to the introduction of *Ae. albopictus* was also consistent with local adaption. Similar to studies from the South Eastern USA and Bermuda [49–54], we have found that species co-occurrence is condition dependant, with the long-term persistence of *Ae. aegypti* occurring throughout many areas despite invasion by *Ae. albopictus* 7 to 15 years ago. Previous studies have suggested *Ae. aegypti* is able to persist in dry climate conditions and/or urban environments because they are better adapted [15 and refs within]. The eggs of *Ae. aegypti* are more tolerant to higher temperatures and desiccation in comparison to the eggs of *Ae. albopictus*, which are able to survive lower temperatures through diapause [15,55]. Consistent with the prediction that local environmental adaptation contributes to *Ae. aegypti* persistence, we found putatively adaptive loci within the dry tropical Pacific regions of Chiriquí (David), Coclé, the eastern Azuero Peninsula and provincial Panamá where both species co-occur. There was also genetic evidence for local adaptation in the isolated wet tropical region of Bocas del Toro and Costa Abajo near Colon, but whether this variation will allow *Ae. aegypti* to resist invasion by *Ae. albopictus* is unknown, given that *Ae. albopictus* was only recorded in Costa Abajo in 2018 and has not yet reached Bocas del Toro. Alternatively, the present patterns of species co-existence could simply reflect the abilities of *Aedes* species to exploit a different ecological niche without involving local environmental adaptation. Nonetheless, this doesn’t reconcile the fact that *Ae. aegypti* is no longer found in many areas where candidate adaptive alleles were not detected. Our findings provide us with clear testable hypotheses moving forward. For example, if the genomic regions we identified are adaptive, then we expect genotype specific survival under different environmental conditions, which can be tested in a common garden with reciprocal transplant experiment in the presence of an ecological competitor.

The presence of locally adapted populations of *Ae. aegypti* could have a significant impact on the future arboviral disease landscape. Climate variables, most notably precipitation and temperature associated with altitudinal and latitudinal clines, are able to drive population differentiation in both *Anopheles* mosquitoes and *Drosophila* flies [56–59]. In the former, the Anopheles gambiae species complex is hypothesised to have radiated through ecological speciation driven by adaptation to aridity and in response to larval habitat competition. This has led to a series of ecotypes with semi-permeable species boundaries [60]. The resulting differences among ecotypes in anthropophily and the adult resting behaviour has a significant impact on malaria transmission risk [61]. Thus, at the most basic level, differentially adapted population variants of *Ae. aegypti* across Panama, could have different abilities to vector arboviral disease [62–67]. In addition, environmental adaptation would need to be considered in spatially predictive models. Currently, species geographic distribution or disease prediction models incorporate a set of environmental parameters coupled with a predicted outcome on mosquito biology and abundance without considering adaptive response [68,69].

Assuming that the whole population will respond to environmental precursors as a homogenous unit is erroneous when local adaptation is present and considering adaptability as a parameter, in combination with the environmental response, will improve the accuracy of future projections [12,70]. Furthermore, the presence of locally adapted populations threatens the efficiency of gene drive systems aimed at promoting disease resistance within mosquito populations. This is because environmental differences between sites, as well as physical geographical barriers, will restrict mosquito dispersal and therefore limit the spread of beneficial alleles or inherited bacteria [71]. However, if locally adaptive alleles are well-characterised, this knowledge could also potentially be exploited. A more tailored approach could improve gene drive efficiency, since locally adapted individuals are theoretically more likely to survive to pass on the intended benefit to the next generation.

If local environmental adaptation is proven to influence *Aedes* co-occurrence, then this could facilitate the emergence of sylvatic arboviral disease. *Ae. albopictus* is an opportunistic feeder, able to utilise a wide range of peri-domestic habitats outside of its native range [72,73] and the species could act as an efficient bridge vector for emergent zoonotic diseases from the forest [73]. The addition of the specialised commensal *Ae. aegypti*, provides the opportunity for any emergent epidemic to spread and be maintained within the urban population [8–11]. This scenario may have happened recently, where yellow fever virus re-emerged from forest reservoirs in Brazil [74]. In this case, the re-emergence was a function of both ecological changes and vaccination frequency. Unlike yellow fever virus, there is no vaccination against dengue, Zika, or chikungunya, reinforcing the role that ecological changes will likely play in future epidemics.

### Conclusion

The identification of small number of putatively adaptive genomic intervals provides exceptional experimental opportunities to determine 1) If these regions are in fact under selection, 2) How selection might be acting if our hypothesis is true. Defining species fitness in association with our candidate loci will allow us to untangle the interplay between genomic process, the environment, species competition and how these resolve the spatial distribution and abundance of medically important *Ae. aegypti*. Advances will be used to improve the accuracy of disease prediction models and characterise the genomic basis of adaptations with the capacity to alter the epidemiological landscape.

## Materials and Methods

### Mosquito Sampling

*Aedes* mosquitoes were collected through active surveillance and oviposition traps placed across 35 settlements and nine provinces of Panama from 2016 and 2018 (S3 Table). Immature stages of *Aedes* from each trap were reared to adulthood as separate collections in the laboratory, identified using the morphological key of Rueda *et al.* [75] and stored in absolute ethanol at −20°C.

### Genomics data

DNA was extracted from 70 *Ae. aegypti* (Fig 1A), representing populations subject to different environmental conditions using a modified phenol chloroform method [76]. To identify putative regions involved in the local adaption of *Ae. aegypti*, 26.74 Mb of the AaeL3 exome were targeted for capture. For each sample, 100 ng DNA was mechanically sheared to fragment sizes of ~ 350-500 base pairs and processed to add Illumina adapters using the Kapa Hyperprep kit. Amplified libraries were assessed on a Bioanalyser and Qubit before 24 uniquely barcoded individuals each were pooled to a combined mass of 1 μg to create three libraries of 24 individuals for hybridization. Sequence capture of exonic regions was performed on each pool according to the NimbleGen SeqCap EZ HyperCap workflow and using custom probes designed by Roche for the regions we specified (S1 Dataset).

Low quality base calls (<20) and Illumina adapters were trimmed from sequence ends with TrimGalore [77], before alignment to the *Ae. aegypti* AaeL5 reference genome with Burrows-Wheeler aligner [78]. Read duplicates were removed with BamUtil. Sequence reads were processed according to the GATK best practise recommendations, trained with a hard-filtered subset of SNPs using online recommendations (https://gatkforums.broadinstitute.org/gatk/discussion/2806/howto-apply-hard-filters-to-a-call-set). SNPs were called with a heterozygosity prior 0.0014 synonymous to previously reported values of theta [24]. Filters applied to the resulting SNP dataset included a minimum quality of 30, minimum depth of 30, minimum mean depth of 20, maximum 95 % missing data across individuals and a minor allele frequency ≥ 0.01. Indels were additionally removed to reduce uncertainty in true variable sites by poor alignment to the reference genome.

### Environmental Data

Climate variables including average rainfall, average humidity, average minimum and maximum temperature difference, average minimum temperature and average maximum temperature were obtained for each collection site from interpolated raster layers composed of values reported by Empresa de Transmisión Eléctrica Panameña (ETESA). All available data points from 2010 to 2017 representing 50-60 meteorological stations across Panama were averaged. NDVI vegetation indexes for Panama were obtained from MODIS Vegetation Indices 16-day L3 Global 250m products (NASA, USA) with values averaged over all available images from 2010 to 2017. Human population density values were obtained from Instituto Nacional de Estadística y Censo 2010. Raster layers for Generalised Dissimilarity Models and Gradient Forest analyses were created for each variable by inverse distance interpolation across the extent of Panama to a resolution of 0.05 pixels in QGIS version 2.18.15 [79].

The collinearity and covariance of the environmental data was assessed the R Stats package [80]. One variable, average minimum and maximum temperature difference was removed from analysis because it was highly correlated with the other temperature variables (>0.8 correlation coefficient). All other variable comparisons had a correlation coefficient below 0.7 and were retained for analysis (S4 Table).

### Analysis of population structure

FastStructure was also applied to all loci to infer the ancestry proportions of K modelled populations [81]. The optimal model complexity (K*e) was chosen to be two populations using the python script chooseK.py and confirmed by a PCA of all loci performed with the R package PCAdapt [82](see Analysis of local environmental adaptation below). FastStructure analysis with a logistic prior was also applied to 2,630 SNP’s shared with a worldwide SNP dataset representing *Ae. aegypti* from 26 different countries [19–23].

### Species distribution analysis

Historical data on species distributions from 2005 to 2017 was obtained from the Panamanian Ministry of Health (MINSA). This data was obtained through active surveillance of settlements regardless of time of year. A binomial Generalised Linear Model was performed to test for an association between the presence and absence of *Ae. aegypti* with the presence and absence of *Ae. albopictus* using the species occurrence data obtained from both MINSA and our own sampling using the Stats package in R [80]. The proportion of sampling sites positive for *Ae. aegypti* and *Ae. albopictus* presence from 2005 through 2018 were calculated by combining our mosquito surveillance data with that obtained from MINSA. Maps of the species distribution of *Ae. aegypti* and *Ae. albopictus* were produced in QGIS [79].

### Analysis of local environmental adaptation

To identify loci with a signal of selection differentiated across regional environmental conditions, three methods with different underlying algorithms and assumptions were applied. Two EAA approaches, redundancy analysis (RDA) and latent factor mixed models (LFMM) were implemented to identify loci associated with environmental predictors. RDA uses multivariate regression to detect genomic variation across environmental predictors as expected from a multilocus signature of selection [25]. In comparison, LFMM is a univariate approach which models background variation using latent factors, while simultaneously correlating the observed genotype frequencies of individuals to each environmental variable [83]. Before implementation of RDA, missing genotype values were imputed as the most common across all individuals. Loci which are strongly correlated to environmental predictors were then identified through multivariate linear regression of the genomic data with the environmental variables followed by constrained ordination of the fitted values as implemented with the RDA function in the R package Vegan [84]. Multi collinearity of the data was verified to be low as indicated by genomic inflation factors ranging from 1.31-5.80. Candidate loci were then identified as those which contribute most to the significant axes as determined by F statistics [85]. To account for population structure, we applied two latent factors to our LFMM analysis based on the PCA and scree plots of proportion of explained variance produced with PCAdapt (see below). As per recommendations to improve power, we filtered our data before analysis to include only sites with an MAF > 5 % and analysed our data with five separate LFMM runs, each with 20,000 cycles after an initial burn-in period of 10,000 cycles. Median Z-scores were calculated from the five runs and Bonferroni corrected for multiple tests, before loci significantly correlated with environmental variables were identified based on a false discovery rate of 10 % using the Benjamini-Hochberg procedure outlined in the program documentation. Visualisation of the Bonferroni adjusted probability values for the loci correlated with each environmental factor revealed that the majority of probability values were at a flat distribution while those correlated with environmental variables were within a peak close to 0, indicating that confounding factors were under control. In addition to the two EAA analyses, PCAdapt was applied to identify loci putatively under selection pressure because they deviate from the typical distribution of the test statistic Z [82]. Two K populations were chosen to account for neutral population structure in the data based on scree plots of the proportion of explained variance and visual inspection of PCA and STRUCTURE plots which revealed that populations from the region of Bocas del Toro form a distinct genomic grouping (Fig 1, S11 Fig).

### Distribution of candidate loci across geographical space

Both putatively neutral and adaptive genomic variation was visualised across geographic space using Generalised Dissimilarity Modelling (GDM) and Gradient Forests (GF) analysis [86]. GDM is a regression-based approach which maps allelic turnover using non-linear functions of environmental distance in relation to FST genetic distance. In comparison, GF uses a machine learning regression tree approach. Through subsetting the genomic and environmental data, the algorithm determines the degree of change for each allele along an environmental gradient and calculates the resulting split importance. Allelic turnover was investigated for both a set of reference SNP’s, not expected to be under selective pressure, as well as the loci putatively involved in local adaptation as jointly identified by LFMM, PCAdapt analysis and RDA. SNP’s representative of neutral variation included those not identified as a candidate outlier by any of the three methods. So as to reduce the dataset and avoid inclusion of strongly linked loci, SNP’s were thinned by a distance of 10 KB, an appropriate cut-off as indicated by the calculation of R2 linkage disequilibrium values for this dataset (S12 Fig).

To perform GDM analysis, the R program StAMPP [87] was used to generate the input FST matrixes and BBmisc [88] used to rescale the distances between 0 and 1. Environmental and genetic distance data were converted to GDM format and analysis performed using the R package GDM [89]. GF analysis [90] was implemented on a matrix of minor allele frequencies for each SNP for both the reference and candidate datasets, obtained through VCFtools [91]. Both SNP datasets only included loci present in at least 11 of 14 populations to ensure robust regression. The model was fitted with 2,000 regression trees, a correlation threshold of 0.5 and variable importance computed by conditional permutation with a distribution maximum of 1.37. Both analyses included Moran’s eigenvector map (MEM) variables which are weightings derived from the geographic coordinates of sampling locations used to model unmeasured environmental variation and geographic distance analogous to latent factors [86]. To visualise the patterns in allele variation across space, PCA was used to reduce the variability into three factors. The difference in genomic composition was mapped across the landscape of Panama by assigning the three centred principle components to RGB colours; similar genomic composition across space is indicated by a similar colour shade. The difference in allele turnover for the reference and candidate dataset was characterised to explore whether allelic turnover was greater than predicted under neutral expectations. Exploration was achieved by comparing and visualising the compositional turnover of allele frequencies for both reference and candidate SNP dataset across geographical space using a Procrustes superimposition on the PCA ordinations.

## Acknowledgements

We are grateful to Panama’s Ministry of Environment (*Mi Ambiente*) and the Panamanian community for supporting our scientific collections of insects in Panama, to Yamileth Chin and Marta Vargas for their guidance in the laboratory and Jose R. Rovira, Carmelo Gómez Martínez and Alejandro Almanza for their assistance in the field.

## Author contributions

The study was designed by KLB, WOM and JRL, and conducted by KLB, WOM and JRL. Sample preparation and laboratory procedures were conducted by KLB. KLB and JRL performed the data analysis and figure preparation. KLB wrote the manuscript with contributions from WOM and JRL.

## Funding

This work was supported by MINSA, the Zika project, the Secretariat for Science, Technology and Innovation (SENACYT) through the research grant IDDS15-047, and from the National System of Investigation (SNI) award to JRL. Research activity by KLB was supported by the Smithsonian Institution Fellowship Program, George Burch Fellowship, The Edward M. and Jeanne C. Kashian Family Foundation Inc and Mr Nicholas Logothetis of Chartwell Consulting Group Inc.

## Ethics Statement

Field research conducted in Panama was authorised under the Ministerio de Ambiente permit number SE/A-67-2016-9.

## Data availability

SNP data is available in the Sequence Read Archive data repository XXX

## Competing interests

The authors received funding from The Edward M. and Jeanne C. Kashian Family Foundation Inc. and Nicholas Logothetis of Chartwell Consulting Group Inc. The funders had no role in study design, data collection and analysis, decision to publish, or preparation of the manuscript. There are no patents, products in development or marketed products associated with this research to declare.

