## Supplementary Material for "The impact of biological invasion and genomic local adaptation on the geographical distribution of *Aedes aegypti* in Panama"

#### Supplementary Figures

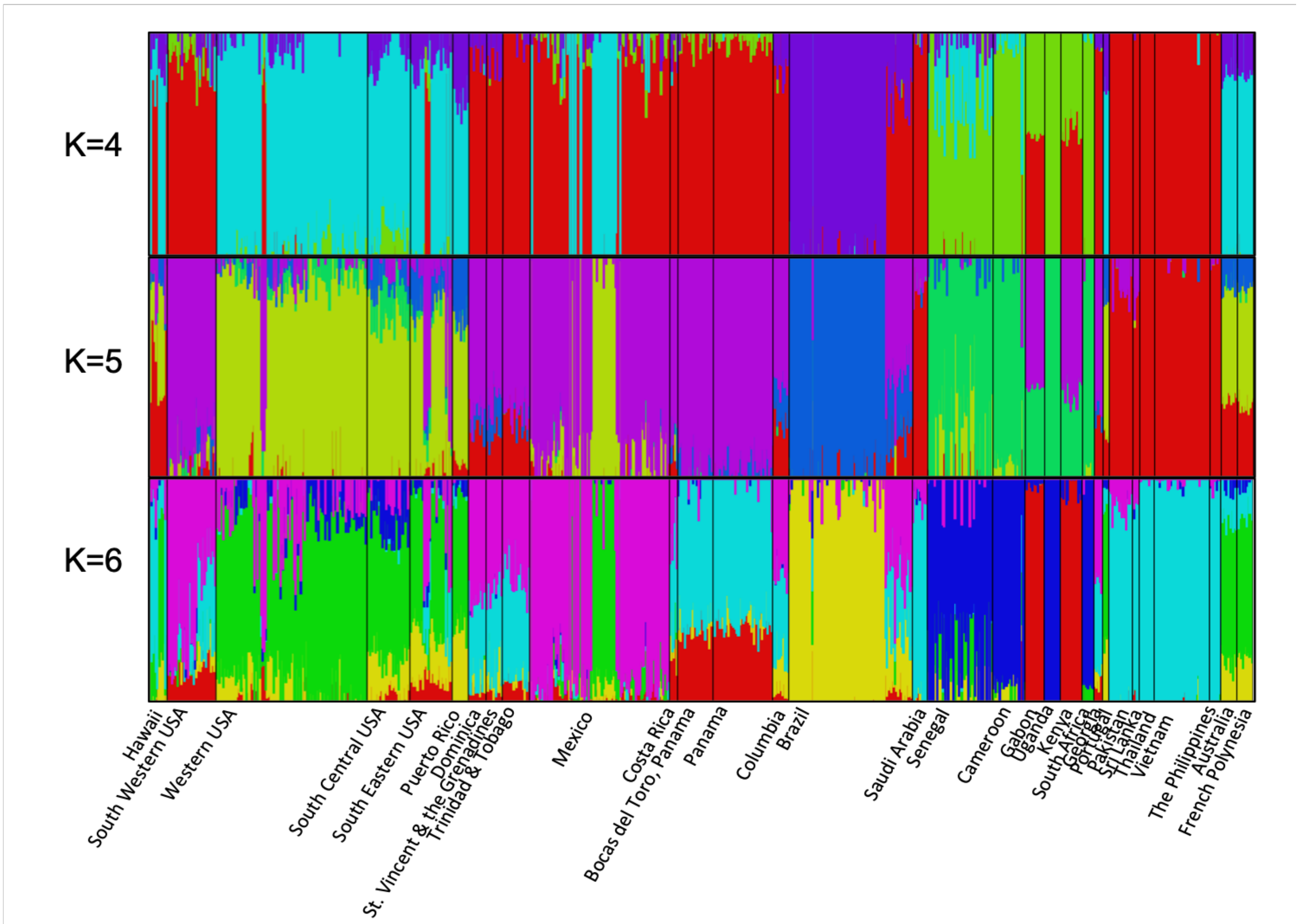


**S1 Fig.** FastStructure plots comparing 2,630 SNP's from individuals of *Ae. aegypti* from Panama to 26 other worldwide locations. Plots are provided for between 4=K populations, the best number of model components used to explain structure in data, and 6=K populations, reflecting the model complexity that maximizes the marginal likelihood. FastStructure assigns each individual to one or more K populations, as indicated by its colour. Genetically similar populations share the same colour or similar admixture composition on comparison within each separate plot.

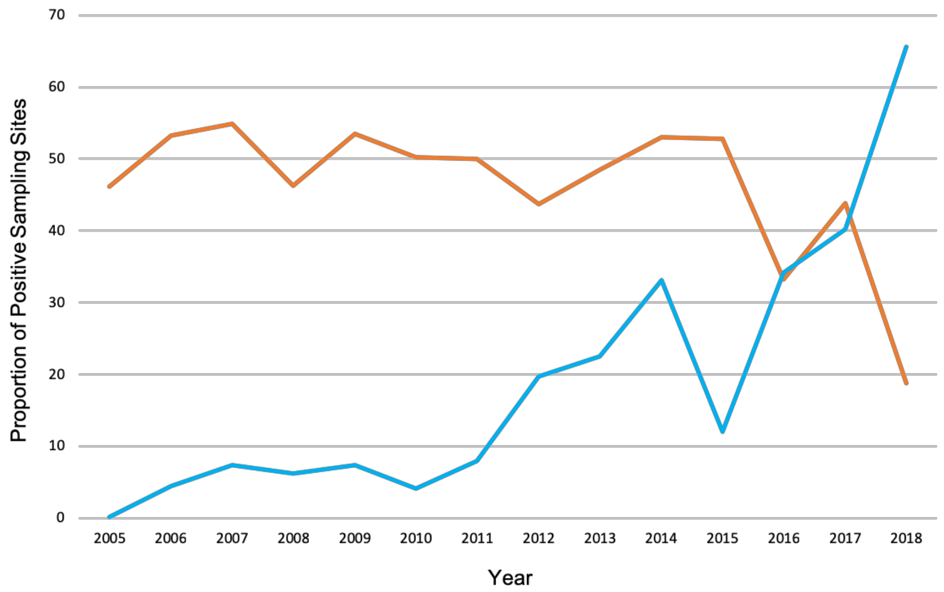


**S2 Fig.** The proportion of sampling sites positive for the presence of *Ae. aegypti* (orange) and *Ae. albopictus* (blue) from 2005 through 2018.

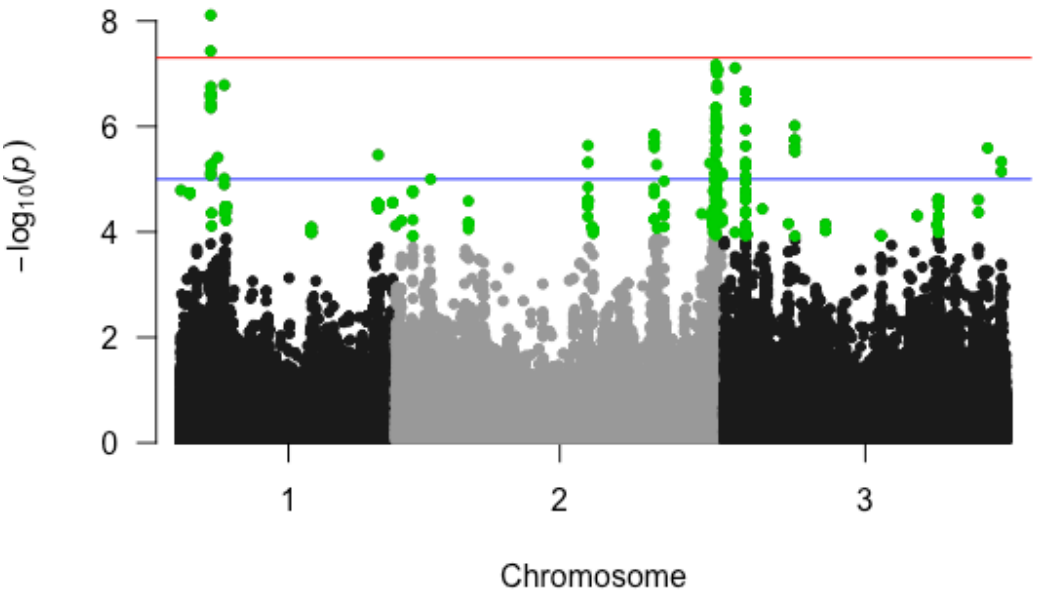


**S3 Fig.** Manhattan plot showing the location of putative adaptive loci across each of the three *Ae. aegypti* chromosomes as identified by PCAdapt. Loci above the red line, which is the threshold for genome-wide significance, show highly significant signals of selection, while those above the blue line are suggestive of selection. Those identified by the analysis as candidate loci with a signal of local adaptation are highlighted in green.

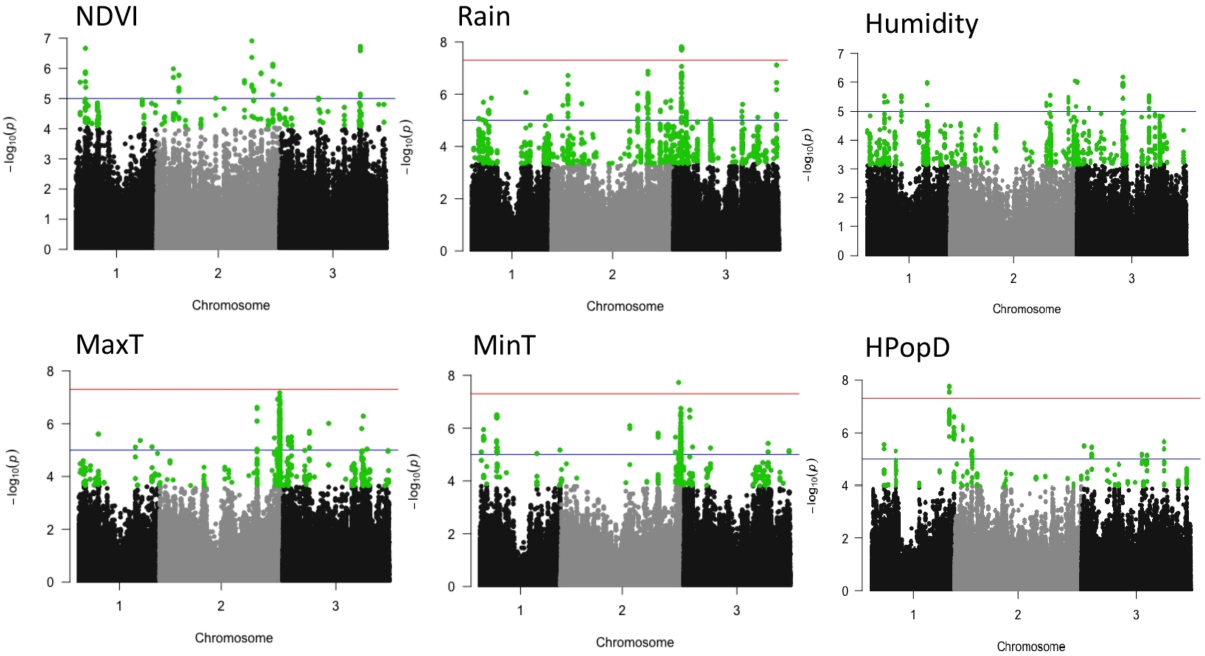


**S4 Fig.** Manhattan plots showing the distribution of genomic loci correlated to NDVI vegetation index (NDVI), average rainfall (Rain), average humidity, average minimum rainfall (MinT), average maximum temperature (MaxT) and human population density (HPopD) across the three chromosomes of the *Ae. aegypti* genome. Loci above the red line, which is the threshold for genome-wide significance, show highly significant signals of selection, while those above the blue line are suggestive of selection. Those identified by the analysis as candidate loci with a signal of local adaptation based on a false discovery rate of 10 % are highlighted in green.

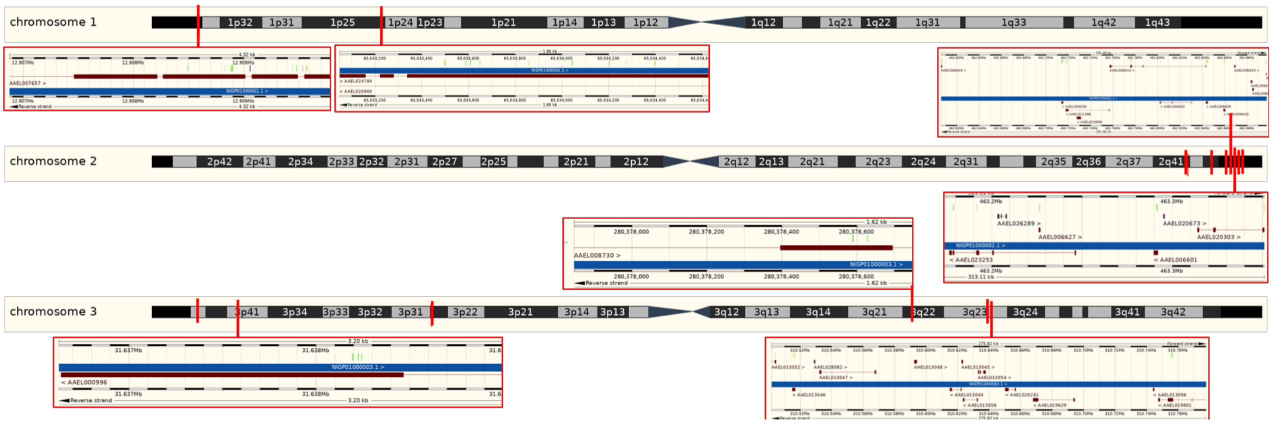


**S5 Fig.** The location of the 128 candidate loci within each of the three chromosomes of the *Ae. aegypti* genome that were identified as putatively involved in local adaptation and shared across selection and EAA analysis. Genomic locations with more than one SNP are expanded to show the position of potentially linked loci, which are annotated as coloured lines above the gene architecture (red bars) and genomic contig (blue bar). Images were created using the genome browser tool provided by VectorBase (www.vectorbase.org).

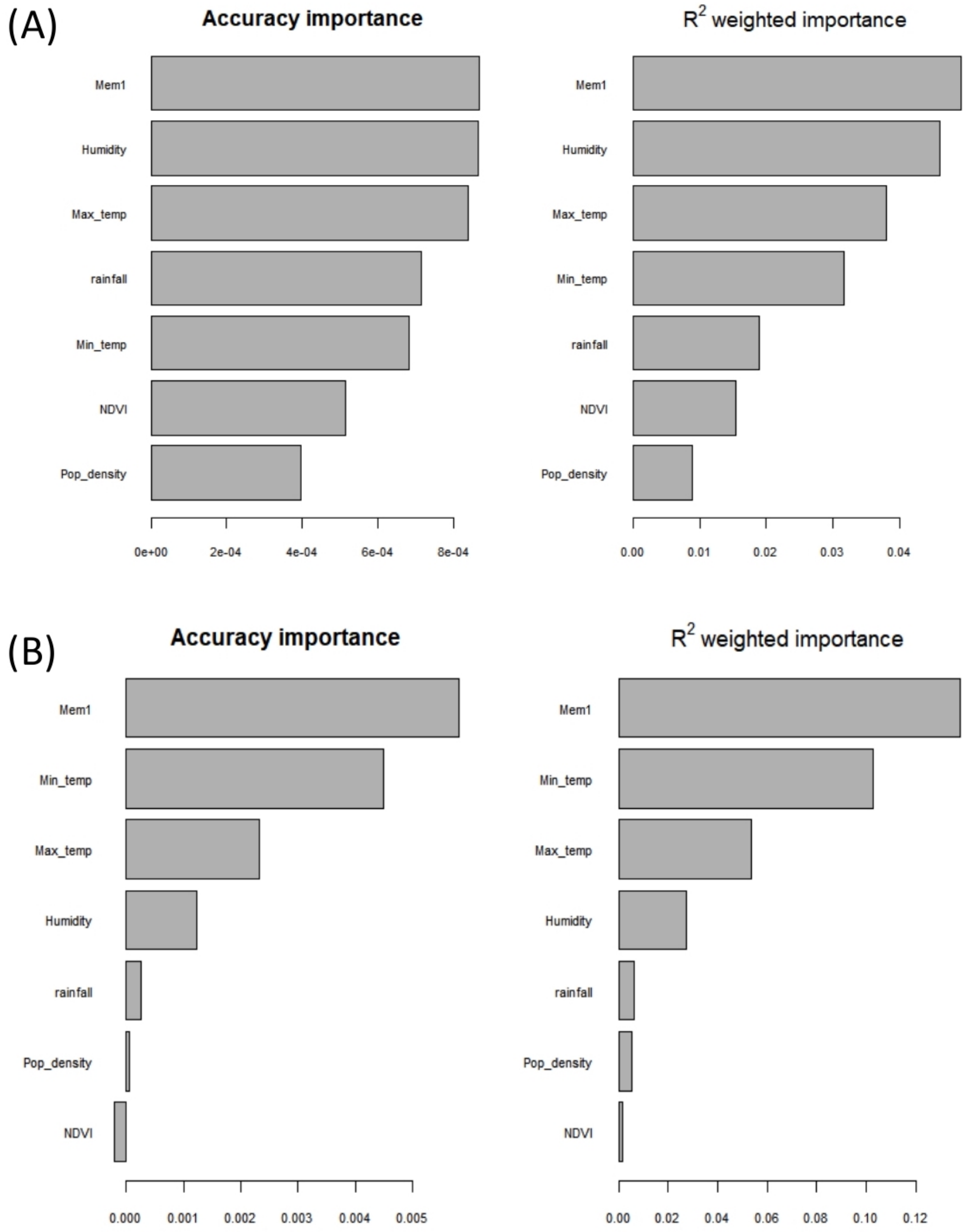


**S6 Fig.** Variable importance of the environmental variables in the GF model for a. reference and b. candidate loci.

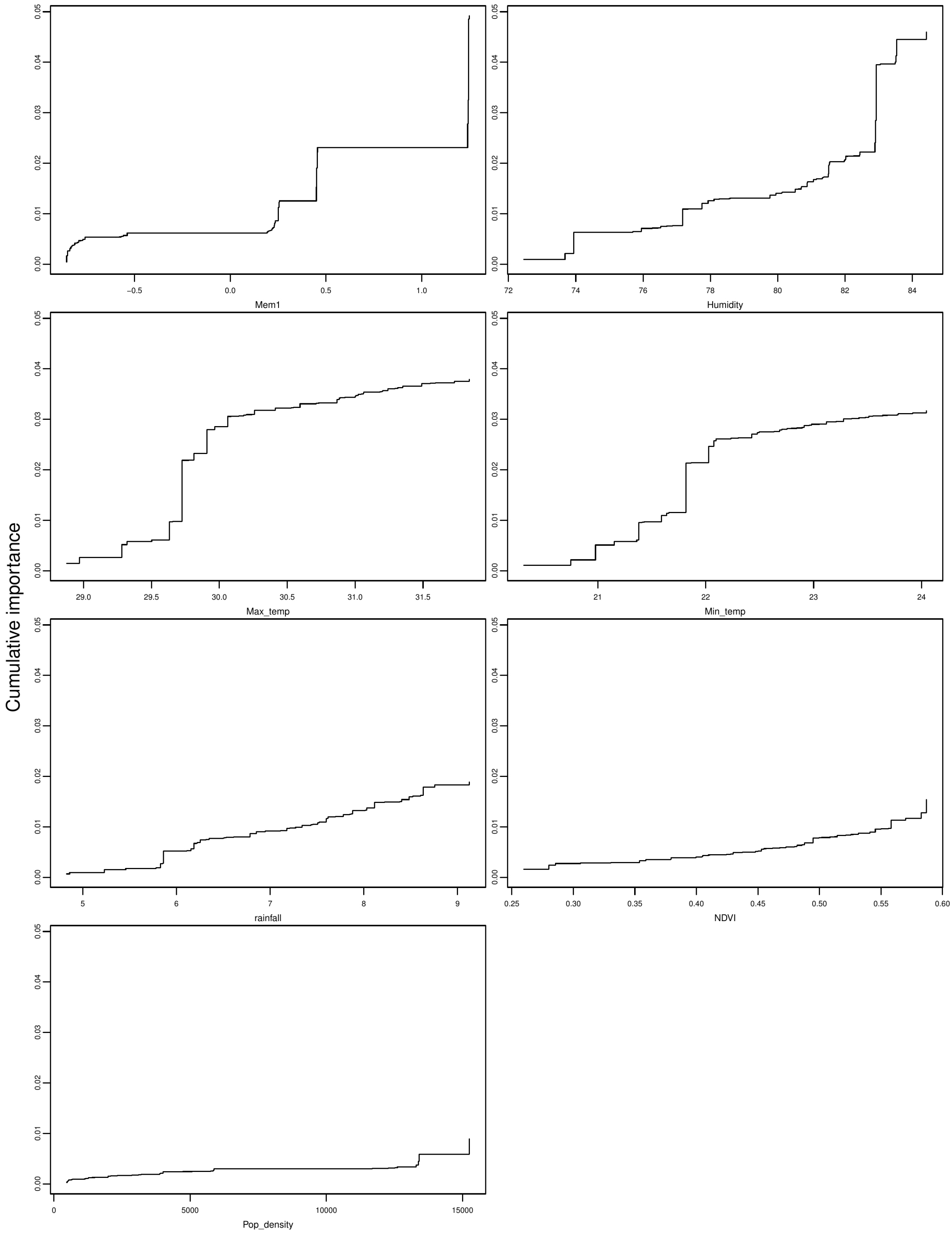


**S7 Fig.** Aggregate compositional turnover plots for GF analysis depicting allele change of the reference loci across each environmental gradient. The maximum height of the line gives the overall change in allele frequency and therefore the relative variable importance.

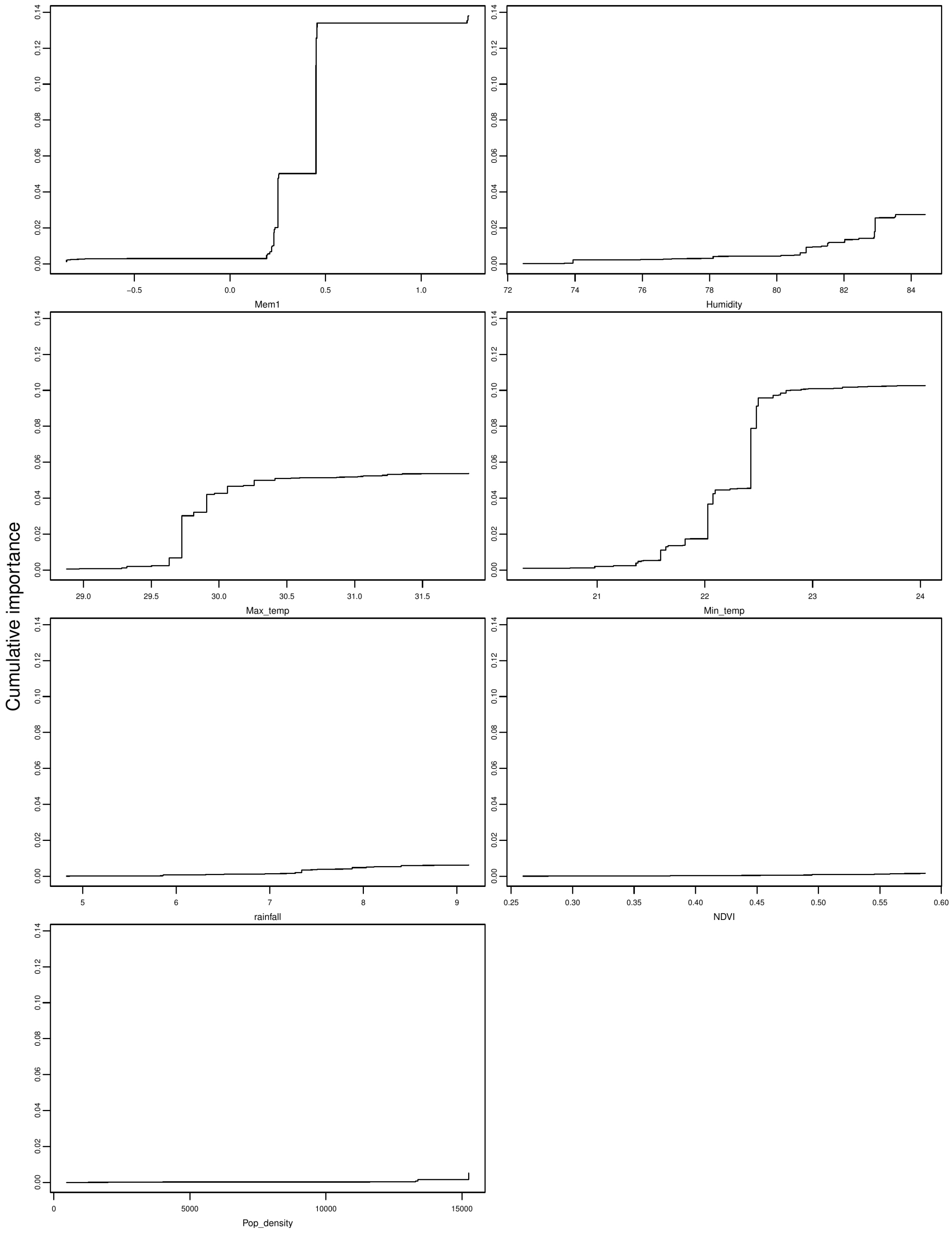


**S8 Fig.** Aggregate compositional turnover plots for GF analysis depicting allele change of the candidate loci across each environmental gradient. The maximum height of the line gives the overall change in allele frequency and therefore the relative variable importance.

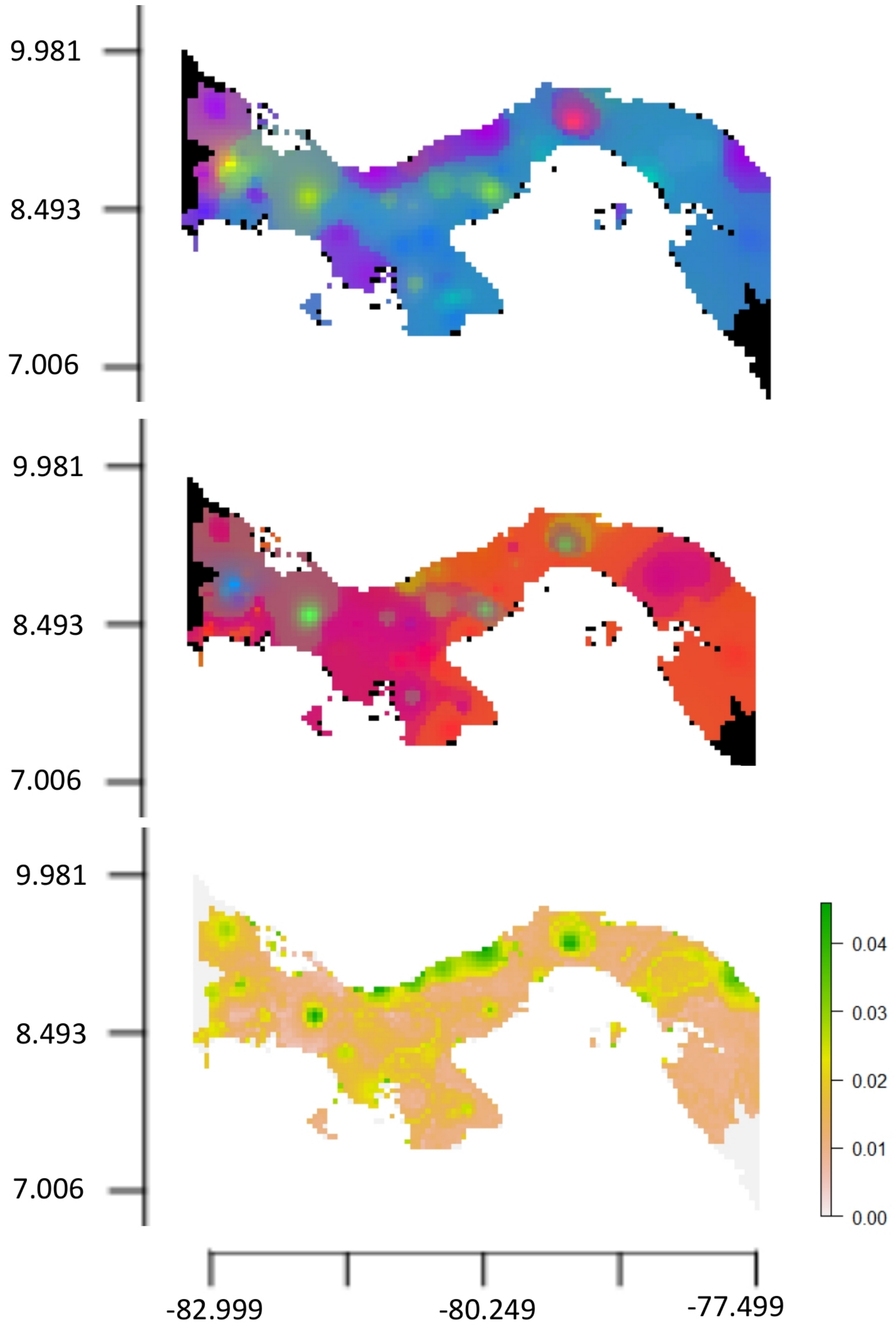


**S9 Fig.** RGB maps of compositional allele frequency turner over across geographical space based on GF analysis for reference loci (above), candidate loci (centre) and the difference in allele compositional turnover between the reference and candidate dataset using a Procrustes superimposition on the PCA ordinations (below). In the above and centre map, dissimilarity between allele composition is depicted by an increasing divergent colour spectrum. Locations with a similar allele composition are a similar colour. On the below map, the scale represents the distance between the allele composition of the reference and candidate SNP datasets, with higher distances indicating areas that are potentially experiencing local adaptation.

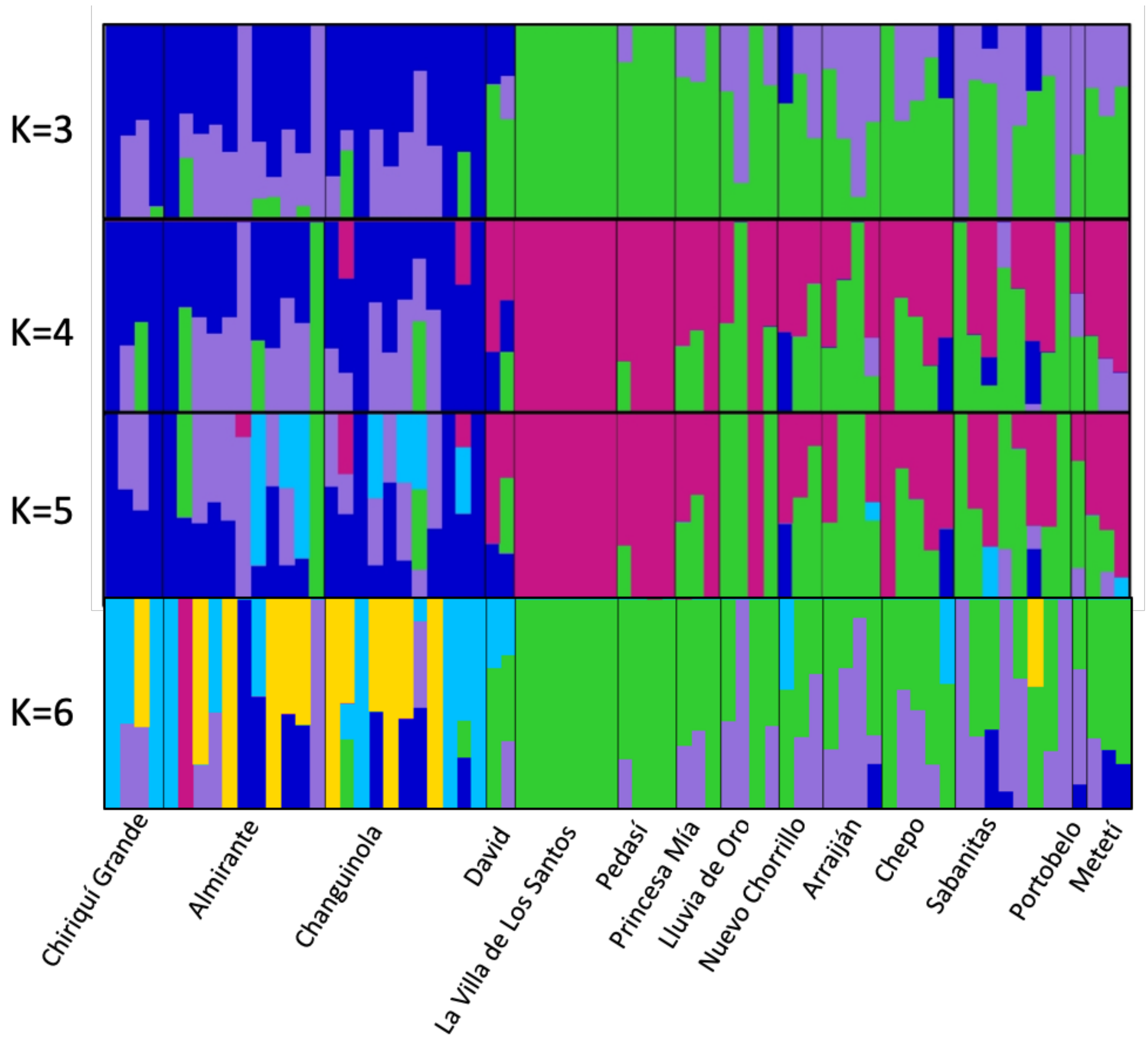


**S10 Fig.** FastStructure plot of 128 candidate loci with a signal of local adaptation for between 3=K (the number of model components used to explain structure in data) and 6=K populations (the model complexity that maximizes the marginal likelihood). FastStructure assigns each individual to one or more K populations, as indicated by its colour. Genetically similar populations share the same colour or similar admixture composition on comparison within each separate plot.

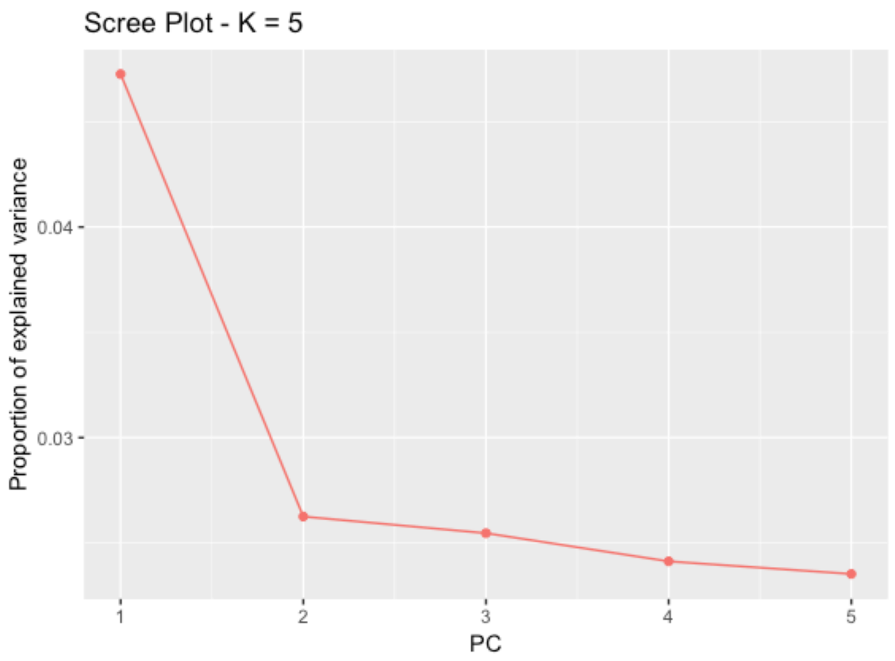


**S11 Fig.** Scree plot produced in PCAdapt showing the selection of K2 populations explains the variance in the genomic data of *Ae. aegypti*.

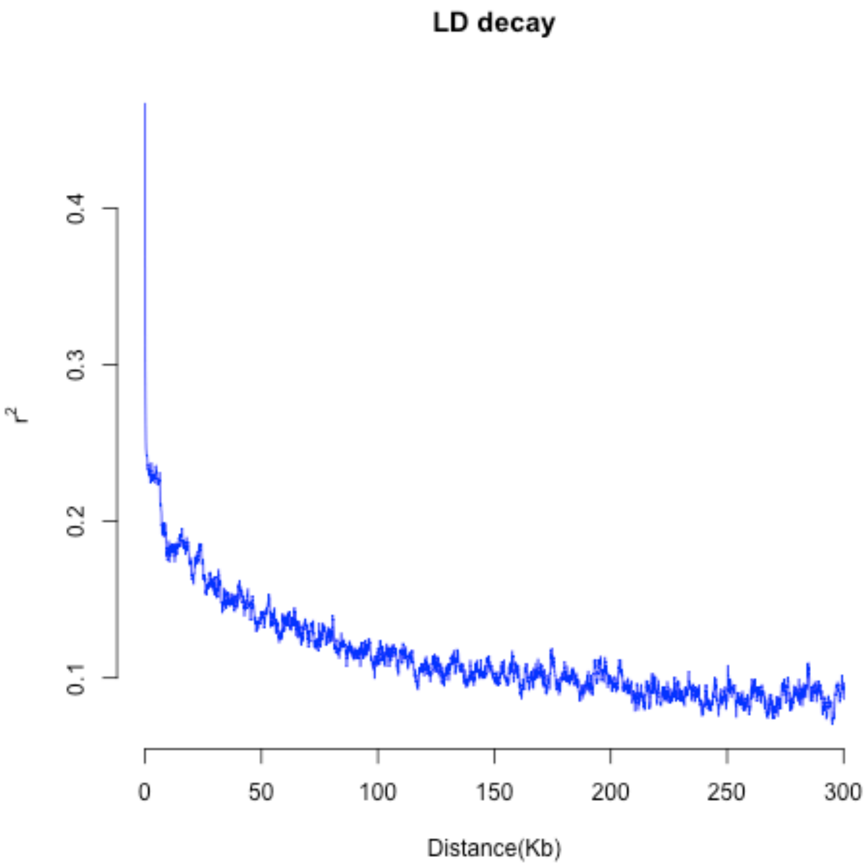


**S12 Fig.** LD decay plot of $R_2$ linkage disequilibrium across all loci the SNP dataset.

### Supplementary Tables

**S1 Table.** The annotated functions of 17 genes that include the 128 candidate loci involved in the local adaptation of *Ae. aegypti* available in Vectorbase.

| Gene stable ID | Gene description | Chromosome/scaffold name | Gene start (bp) | Gene end (bp) | GO term name |
|---|---|---|---|---|---|
| AAEL007657 | low-density lipoprotein receptor (ldl) | 1 | 12873445 | 12913096 | integral component of membrane |
| | | | | | calcium ion binding |
| AAEL001710 | | 2 | 441958067 | 442766939 | protein localization involved in establishment of planar polarity |
| | | | | | adherens junction |
| AAEL019568 | | 2 | 453369647 | 453398010 | ATP binding |
| | | | | | S-adenosylmethionine biosynthetic process |
| | | | | | methionine adenosyltransferase activity |
| AAEL020638 | | 2 | 459261854 | 459288485 | kinase activity |
| | | | | | protein binding |
| | | | | | protein serine/threonine kinase activity |
| | | | | | protein-containing complex binding |
| AAEL006609 | zinc finger protein | 2 | 462601938 | 462606861 | nucleic acid binding |
| AAEL006610 | | 2 | 462757004 | 462843291 | protein binding |
| AAEL006607 | juvenile hormone-inducible protein, putative | 2 | 462884609 | 462886033 | |
| AAEL006630 | | 2 | 462897060 | 462898678 | |
| AAEL006613 | pickpocket | 2 | 463027538 | 463039524 | membrane |
| | | | | | integral component of membrane |
| | | | | | sodium ion transport |
| | | | | | sodium channel activity |
| AAEL006627 | serine-type enodpeptidase | 2 | 463226014 | 463226973 | proteolysis |
| | | | | | serine-type endopeptidase activity |
| AAEL020303 | | 2 | 463313906 | 463367902 | ATP binding |
| | | | | | integral component of membrane |
| | | | | | transmembrane transport |
| | | | | | ATPase activity |
| AAEL002683 | aldehyde oxidase | 2 | 464805900 | 464819668 | molybdopterin cofactor binding |
| | | | | | electron transfer activity |
| | | | | | oxidoreductase activity |
| | | | | | oxidation-reduction process |
| | | | | | flavin adenine dinucleotide binding |
| | | | | | xanthine oxidase activity |
| | | | | | xanthine dehydrogenase activity |
| | | | | | metal ion binding |
| | | | | | iron ion binding |
| | | | | | iron-sulfur cluster binding |
| | | | | | 2 iron, 2 sulfur cluster binding |
| | | | | | FAD binding |
| AAEL002684 | dachshund | 2 | 465614172 | 465764066 | |
| AAEL000010 | 60S ribosomal protein L36 (Rpl36-1) | 3 | 102915808 | 102917099 | structural constituent of ribosome |
| | | | | | translation |
| | | | | | ribosome |
| | | | | | intracellular |
| AAEL008730 | anillin/rhotekin (rtkn) | 3 | 280355185 | 280379917 | |
| AAEL005305 | | 3 | 308006495 | 308442172 | |
| AAEL013052 | ubiquitin carboxyl-terminal hydrolase isozyme | 3 | 310492268 | 310504566 | ubiquitin-dependent protein catabolic process |
| | | | | | thiol-dependent ubiquitin-specific protease activity |
| | | | | | intracellular |

**S2 Table.** Variable importance explained by each variable in the full GDM model for the reference and candidate dataset including the geographic parameter MEM, NDVI vegetation index, rainfall, humidity, human population density (HPopD), average maximum temperature (MaxT) and average minimum temperature (MinT). Variable importance is measured as the percent change in deviance explained by the full model and the deviance explained by a model fit with the variable permuted.

| | Reference loci | Candidate loci |
|---|---|---|
| All variables | 23.23 | 53.37 |
| Geographic | 1.70 | 7.93 |
| NDVI | 0.00 | NA |
| Rainfall | 9.25 | 0.01 |
| Humidity | 17.81 | 0.53 |
| HPopD | 0.10 | 0.00 |
| MaxT | 9.46 | 3.38 |
| MinT | 3.17 | 1.53 |

**S3 Table.** The sampling localities and presence absence data for *Aedes* mosquitoes.

| Sample Code | Province | Locality | Latitude | Longitude | Year | *Ae. aegypti* presence | *Ae. albopictus* presence |
|---|---|---|---|---|---|---|---|
| BT1_1 | Bocas del Toro | Chiriquí Grande | 8.94399 | -82.11361 | 2016 | 0 | 0 |
| BT1_10 | Bocas del Toro | Chiriquí Grande | 8.94939 | -82.11790 | 2016 | 0 | 0 |
| BT1_11 | Bocas del Toro | Chiriquí Grande | 8.94945 | -82.11875 | 2016 | 0 | 0 |
| BT1_12 | Bocas del Toro | Chiriquí Grande | 8.94990 | -82.11907 | 2016 | 0 | 0 |
| BT1_13 | Bocas del Toro | Chiriquí Grande | 8.95018 | -82.11984 | 2016 | 0 | 1 |
| BT1_14 | Bocas del Toro | Chiriquí Grande | 8.95074 | -82.11977 | 2016 | 0 | 0 |
| BT1_16 | Bocas del Toro | Chiriquí Grande | 8.95194 | -82.12145 | 2016 | 0 | 0 |
| BT1_17 | Bocas del Toro | Chiriquí Grande | 8.95242 | -82.12221 | 2016 | 0 | 0 |
| BT1_18 | Bocas del Toro | Chiriquí Grande | 8.95228 | -82.12260 | 2016 | 0 | 0 |
| BT1_19 | Bocas del Toro | Chiriquí Grande | 8.95226 | -82.12425 | 2016 | 0 | 0 |
| BT1_2 | Bocas del Toro | Chiriquí Grande | 8.94473 | -82.11446 | 2016 | 0 | 0 |
| BT1_20 | Bocas del Toro | Chiriquí Grande | 8.94739 | -82.11976 | 2016 | 0 | 0 |
| BT1_21 | Bocas del Toro | Chiriquí Grande | 8.94827 | -82.12073 | 2016 | 0 | 0 |
| BT1_22 | Bocas del Toro | Chiriquí Grande | 8.94934 | -82.12201 | 2016 | 0 | 0 |
| BT1_23 | Bocas del Toro | Chiriquí Grande | 8.95082 | -82.12344 | 2016 | 0 | 0 |
| BT1_3 | Bocas del Toro | Chiriquí Grande | 8.94596 | -82.11524 | 2016 | 0 | 0 |
| BT1_4 | Bocas del Toro | Chiriquí Grande | 8.94754 | -82.11591 | 2016 | 0 | 0 |
| BT1_5 | Bocas del Toro | Chiriquí Grande | 8.94806 | -82.11718 | 2016 | 0 | 0 |
| BT1_6 | Bocas del Toro | Chiriquí Grande | 8.94812 | -82.11801 | 2016 | 0 | 0 |
| BT1_7 | Bocas del Toro | Chiriquí Grande | 8.94771 | -82.11731 | 2016 | 0 | 0 |
| BT1_8 | Bocas del Toro | Chiriquí Grande | 8.94978 | -82.11705 | 2016 | 0 | 0 |
| BT1_9 | Bocas del Toro | Chiriquí Grande | 8.94903 | -82.11730 | 2016 | 0 | 0 |
| BT2_1 | Bocas del Toro | Almirante | 9.29017 | -82.39680 | 2016 | 1 | 0 |
| BT2_10 | Bocas del Toro | Almirante | 9.29465 | -82.38886 | 2016 | 0 | 0 |

| | | | | | | | | |
|---|---|---|---|---|---|---|---|---|
| BT2_11 | Bocas del Toro | Almirante | 9.29425 | -82.38973 | 2016 | | 0 | 0 |
| BT2_12 | Bocas del Toro | Almirante | 9.29472 | -82.39139 | 2016 | | 0 | 0 |
| BT2_13 | Bocas del Toro | Almirante | 9.29659 | -82.40825 | 2016 | | 0 | 0 |
| BT2_14 | Bocas del Toro | Almirante | 9.29651 | -82.40711 | 2016 | | 1 | 0 |
| BT2_15 | Bocas del Toro | Almirante | 9.29423 | -82.40693 | 2016 | | 0 | 0 |
| BT2_16 | Bocas del Toro | Almirante | 9.29736 | -82.40921 | 2016 | | 0 | 0 |
| BT2_17 | Bocas del Toro | Almirante | 9.29450 | -82.40608 | 2016 | | 1 | 0 |
| BT2_18 | Bocas del Toro | Almirante | 9.29099 | -82.40363 | 2016 | | 1 | 0 |
| BT2_19 | Bocas del Toro | Almirante | 9.29288 | -82.40395 | 2016 | | 0 | 0 |
| BT2_2 | Bocas del Toro | Almirante | 9.28995 | -82.39794 | 2016 | | 0 | 0 |
| BT2_20 | Bocas del Toro | Almirante | 9.29491 | -82.39933 | 2016 | | 1 | 0 |
| BT2_3 | Bocas del Toro | Almirante | 9.29147 | -82.39667 | 2016 | | 0 | 0 |
| BT2_4 | Bocas del Toro | Almirante | 9.29107 | -82.39888 | 2016 | | 0 | 0 |
| BT2_5 | Bocas del Toro | Almirante | 9.28974 | -82.39985 | 2016 | | 0 | 0 |
| BT2_6 | Bocas del Toro | Almirante | 9.28996 | -82.40099 | 2016 | | 1 | 0 |
| BT2_7 | Bocas del Toro | Almirante | 9.29073 | -82.40112 | 2016 | | 0 | 0 |
| BT2_8 | Bocas del Toro | Almirante | 9.29240 | -82.39980 | 2016 | | 0 | 0 |
| BT2_9 | Bocas del Toro | Almirante | 9.29365 | -82.38852 | 2016 | | 0 | 0 |
| BT4_1 | Bocas del Toro | Changuinola | 9.47032 | -82.51345 | 2016 | | 1 | 0 |
| BT4_10 | Bocas del Toro | Changuinola | 9.46439 | -82.51952 | 2016 | | 1 | 0 |
| BT4_11 | Bocas del Toro | Changuinola | 9.45115 | -82.52225 | 2016 | | 1 | 0 |
| BT4_12 | Bocas del Toro | Changuinola | 9.45086 | -82.52531 | 2016 | | 0 | 0 |
| BT4_13 | Bocas del Toro | Changuinola | 9.44962 | -82.52017 | 2016 | | 0 | 0 |
| BT4_14 | Bocas del Toro | Changuinola | 9.44684 | -82.52113 | 2016 | | 1 | 0 |
| BT4_15 | Bocas del Toro | Changuinola | 9.44655 | -82.52294 | 2016 | | 1 | 0 |
| BT4_16 | Bocas del Toro | Changuinola | 9.44623 | -82.52434 | 2016 | | 0 | 0 |
| BT4_17 | Bocas del Toro | Changuinola | 9.44153 | -82.51825 | 2016 | | 1 | 0 |
| BT4_18 | Bocas del Toro | Changuinola | 9.43146 | -82.51944 | 2016 | | 0 | 0 |
| BT4_19 | Bocas del Toro | Changuinola | 9.43579 | -82.51982 | 2016 | | 1 | 0 |
| BT4_2 | Bocas del Toro | Changuinola | 9.47191 | -82.51326 | 2016 | | 1 | 0 |
| BT4_20 | Bocas del Toro | Changuinola | 9.43413 | -82.51972 | 2016 | | 0 | 0 |
| BT4_21 | Bocas del Toro | Changuinola | 9.43192 | -82.51957 | 2016 | | 0 | 0 |
| BT4_22 | Bocas del Toro | Changuinola | 9.42888 | -82.51981 | 2016 | | 0 | 0 |
| BT4_3 | Bocas del Toro | Changuinola | 9.47204 | -82.51458 | 2016 | | 1 | 0 |
| BT4_4 | Bocas del Toro | Changuinola | 9.47294 | -82.51033 | 2016 | | 1 | 0 |
| BT4_5 | Bocas del Toro | Changuinola | 9.47470 | -82.51148 | 2016 | | 0 | 0 |
| BT4_6 | Bocas del Toro | Changuinola | 9.47369 | -82.51311 | 2016 | | 0 | 0 |
| BT4_7 | Bocas del Toro | Changuinola | 9.47455 | -82.50920 | 2016 | | 1 | 0 |
| BT4_8 | Bocas del Toro | Changuinola | 9.47723 | -82.51037 | 2016 | | 1 | 0 |
| BT4_9 | Bocas del Toro | Changuinola | 9.46633 | -82.51819 | 2016 | | 0 | 0 |
| CH1_1 | Chiriquí | San Vincente | 8.59778 | -82.64417 | 2016 | | 0 | 1 |
| CH1_10 | Chiriquí | San Vincente | 8.59611 | -82.63556 | 2016 | | 0 | 1 |
| CH1_11 | Chiriquí | San Vincente | 8.60000 | -82.63444 | 2016 | | 0 | 1 |

| | | | | | | | |
|---|---|---|---|---|---|---|---|
| CH1_12 | Chiriquí | San Vincente | 8.60222 | -82.63661 | 2016 | 0 | 1 |
| CH1_13 | Chiriquí | San Vincente | 8.60694 | -82.63194 | 2016 | 0 | 0 |
| CH1_2 | Chiriquí | San Vincente | 8.59583 | -82.63639 | 2016 | 0 | 1 |
| CH1_3 | Chiriquí | San Vincente | 8.58417 | -82.63722 | 2016 | 0 | 1 |
| CH1_4 | Chiriquí | San Vincente | 8.58611 | -82.63694 | 2016 | 0 | 0 |
| CH1_5 | Chiriquí | San Vincente | 8.59444 | -82.62722 | 2016 | 0 | 1 |
| CH1_6 | Chiriquí | San Vincente | 8.57583 | -82.63611 | 2016 | 0 | 1 |
| CH1_7 | Chiriquí | San Vincente | 8.57861 | -82.63806 | 2016 | 0 | 1 |
| CH1_8 | Chiriquí | San Vincente | 8.58863 | -82.63667 | 2016 | 0 | 1 |
| CH1_9 | Chiriquí | San Vincente | 8.59417 | -82.63611 | 2016 | 0 | 1 |
| CH2_1 | Chiriquí | Bongo Abajo | 8.57639 | -82.61667 | 2016 | 0 | 0 |
| CH2_2 | Chiriquí | Bongo Abajo | 8.57472 | -82.61806 | 2016 | 0 | 0 |
| CH2_3 | Chiriquí | Bongo Abajo | 8.57361 | -82.61778 | 2016 | 0 | 1 |
| CH2_4 | Chiriquí | Bongo Abajo | 8.57306 | -82.61778 | 2016 | 0 | 0 |
| CH2_5 | Chiriquí | Bongo Abajo | 8.56917 | -82.61500 | 2016 | 0 | 1 |
| CH2_6 | Chiriquí | Bongo Abajo | 8.56833 | -82.61361 | 2016 | 0 | 0 |
| CH2_7 | Chiriquí | Bongo Abajo | 8.56694 | -82.61222 | 2016 | 0 | 1 |
| CH2_8 | Chiriquí | Bongo Abajo | 8.56583 | -82.61250 | 2016 | 0 | 1 |
| CH2_9 | Chiriquí | Bongo Abajo | 8.63167 | -82.61250 | 2016 | 0 | 1 |
| CH3_1 | Chiriquí | Volcán | 8.78250 | -82.64194 | 2016 | 0 | 0 |
| CH3_10 | Chiriquí | Volcán | 8.77722 | -82.65000 | 2016 | 0 | 0 |
| CH3_11 | Chiriquí | Volcán | 8.77278 | -82.64861 | 2016 | 0 | 0 |
| CH3_12 | Chiriquí | Volcán | 8.77917 | -82.64369 | 2016 | 0 | 0 |
| CH3_13 | Chiriquí | Volcán | 8.77528 | -82.64778 | 2016 | 0 | 0 |
| CH3_2 | Chiriquí | Volcán | 8.78861 | -82.64944 | 2016 | 0 | 1 |
| CH3_3 | Chiriquí | Volcán | 8.78472 | -82.64750 | 2016 | 0 | 0 |
| CH3_4 | Chiriquí | Volcán | 8.78694 | -82.64750 | 2016 | 0 | 1 |
| CH3_5 | Chiriquí | Volcán | 8.78306 | -82.65250 | 2016 | 0 | 1 |
| CH3_6 | Chiriquí | Volcán | 8.78972 | -82.64361 | 2016 | 0 | 0 |
| CH3_7 | Chiriquí | Volcán | 8.78250 | -82.65167 | 2016 | 0 | 0 |
| CH3_8 | Chiriquí | Volcán | 8.77917 | -82.65472 | 2016 | 0 | 0 |
| CH3_9 | Chiriquí | Volcán | 8.78306 | -82.64500 | 2016 | 0 | 1 |
| CH4_1 | Chiriquí | Cerro Punta | 8.86861 | -82.56000 | 2016 | 0 | 0 |
| CH4_10 | Chiriquí | Cerro Punta | 8.86944 | -82.56639 | 2016 | 0 | 0 |
| CH4_11 | Chiriquí | Cerro Punta | 8.86917 | -82.56750 | 2016 | 0 | 0 |
| CH4_2 | Chiriquí | Cerro Punta | 8.86639 | -82.55917 | 2016 | 0 | 0 |
| CH4_3 | Chiriquí | Cerro Punta | 8.86722 | -82.55872 | 2016 | 0 | 0 |
| CH4_4 | Chiriquí | Cerro Punta | 8.86861 | -82.56028 | 2016 | 0 | 0 |
| CH4_5 | Chiriquí | Cerro Punta | 8.86889 | -82.56028 | 2016 | 0 | 0 |
| CH4_6 | Chiriquí | Cerro Punta | 8.86917 | -82.56056 | 2016 | 0 | 0 |
| CH4_7 | Chiriquí | Cerro Punta | 8.86972 | -82.56139 | 2016 | 0 | 0 |
| CH4_8 | Chiriquí | Cerro Punta | 8.86044 | -82.56250 | 2016 | 0 | 0 |
| CH4_9 | Chiriquí | Cerro Punta | 8.86917 | -82.56417 | 2016 | 0 | 0 |

| | | | | | | | |
|---|---|---|---|---|---|---|---|
| CH5_1 | Chiriquí | Gualaca | 8.53691 | -82.29826 | 2016 | 0 | 0 |
| CH5_10 | Chiriquí | Gualaca | 8.53895 | -82.29831 | 2016 | 0 | 1 |
| CH5_11 | Chiriquí | Gualaca | 8.53684 | -82.30409 | 2016 | 0 | 0 |
| CH5_12 | Chiriquí | Gualaca | 8.53658 | -82.30148 | 2016 | 0 | 0 |
| CH5_13 | Chiriquí | Gualaca | 8.53307 | -82.30199 | 2016 | 0 | 0 |
| CH5_14 | Chiriquí | Gualaca | 8.52971 | -82.30306 | 2016 | 0 | 0 |
| CH5_15 | Chiriquí | Gualaca | 8.52889 | -82.30106 | 2016 | 0 | 1 |
| CH5_16 | Chiriquí | Gualaca | 8.52618 | -82.30067 | 2016 | 0 | 0 |
| CH5_17 | Chiriquí | Gualaca | 8.52426 | -82.29956 | 2016 | 0 | 0 |
| CH5_18 | Chiriquí | Gualaca | 8.52509 | -82.29769 | 2016 | 0 | 1 |
| CH5_19 | Chiriquí | Gualaca | 8.52754 | -82.29865 | 2016 | 0 | 0 |
| CH5_2 | Chiriquí | Gualaca | 8.53505 | -82.29842 | 2016 | 0 | 0 |
| CH5_20 | Chiriquí | Gualaca | 8.53001 | -82.29874 | 2016 | 0 | 0 |
| CH5_3 | Chiriquí | Gualaca | 8.53362 | -82.29791 | 2016 | 0 | 0 |
| CH5_4 | Chiriquí | Gualaca | 8.53174 | -82.29622 | 2016 | 0 | 1 |
| CH5_5 | Chiriquí | Gualaca | 8.52857 | -82.29668 | 2016 | 0 | 0 |
| CH5_6 | Chiriquí | Gualaca | 8.53117 | -82.29746 | 2016 | 0 | 0 |
| CH5_7 | Chiriquí | Gualaca | 8.53147 | -82.30013 | 2016 | 0 | 0 |
| CH5_8 | Chiriquí | Gualaca | 8.53433 | -82.30030 | 2016 | 0 | 0 |
| CH5_9 | Chiriquí | Gualaca | 8.53638 | -82.30007 | 2016 | 0 | 1 |
| CH6_1 | Chiriquí | Paso Canoas | 8.53451 | -82.83715 | 2016 | 0 | 0 |
| CH6_10 | Chiriquí | Paso Canoas | 8.54198 | -82.83913 | 2016 | 0 | 1 |
| CH6_11 | Chiriquí | Paso Canoas | 8.54339 | -82.83893 | 2016 | 0 | 0 |
| CH6_12 | Chiriquí | Paso Canoas | 8.53864 | -82.83928 | 2016 | 0 | 1 |
| CH6_13 | Chiriquí | Paso Canoas | 8.53164 | -82.83695 | 2016 | 0 | 1 |
| CH6_14 | Chiriquí | Paso Canoas | 8.53069 | -82.83661 | 2016 | 0 | 1 |
| CH6_15 | Chiriquí | Paso Canoas | 8.52982 | -82.83653 | 2016 | 0 | 0 |
| CH6_16 | Chiriquí | Paso Canoas | 8.52621 | -82.83479 | 2016 | 0 | 0 |
| CH6_17 | Chiriquí | Paso Canoas | 8.52420 | -82.83292 | 2016 | 0 | 0 |
| CH6_18 | Chiriquí | Paso Canoas | 8.52477 | -82.83380 | 2016 | 0 | 0 |
| CH6_19 | Chiriquí | Paso Canoas | 8.53765 | -82.83777 | 2016 | 0 | 0 |
| CH6_2 | Chiriquí | Paso Canoas | 8.53560 | -82.83738 | 2016 | 0 | 0 |
| CH6_20 | Chiriquí | Paso Canoas | 8.54917 | -82.83752 | 2016 | 0 | 0 |
| CH6_3 | Chiriquí | Paso Canoas | 8.53659 | -82.83690 | 2016 | 0 | 0 |
| CH6_4 | Chiriquí | Paso Canoas | 8.53606 | -82.83514 | 2016 | 0 | 0 |
| CH6_5 | Chiriquí | Paso Canoas | 8.53750 | -82.83510 | 2016 | 0 | 0 |
| CH6_6 | Chiriquí | Paso Canoas | 8.53792 | -82.83708 | 2016 | 1 | 0 |
| CH6_7 | Chiriquí | Paso Canoas | 8.53871 | -82.83768 | 2016 | 0 | 0 |
| CH6_8 | Chiriquí | Paso Canoas | 8.53863 | -82.83656 | 2016 | 0 | 0 |
| CH6_9 | Chiriquí | Paso Canoas | 8.53935 | -82.83892 | 2016 | 0 | 0 |
| CH7_1 | Chiriquí | David | 8.42182 | -82.42606 | 2016 | 0 | 0 |
| CH7_10 | Chiriquí | David | 8.42427 | -82.42539 | 2016 | 1 | 0 |
| CH7_11 | Chiriquí | David | 8.42429 | -82.42293 | 2016 | 0 | 0 |

| | | | | | | | |
|---|---|---|---|---|---|---|---|
| CH7_12 | Chiriquí | David | 8.42485 | -82.41992 | 2016 | 0 | 1 |
| CH7_13 | Chiriquí | David | 8.42218 | -82.42005 | 2016 | 0 | 0 |
| CH7_14 | Chiriquí | David | 8.41989 | -82.42118 | 2016 | 0 | 0 |
| CH7_15 | Chiriquí | David | 8.41874 | -82.42333 | 2016 | 1 | 0 |
| CH7_16 | Chiriquí | David | 8.41643 | -82.42270 | 2016 | 0 | 1 |
| CH7_17 | Chiriquí | David | 8.41771 | -82.42129 | 2016 | 0 | 1 |
| CH7_18 | Chiriquí | David | 8.41619 | -82.41977 | 2016 | 0 | 0 |
| CH7_19 | Chiriquí | David | 8.41821 | -82.41808 | 2016 | 0 | 1 |
| CH7_2 | Chiriquí | David | 8.42024 | -82.42592 | 2016 | 0 | 0 |
| CH7_20 | Chiriquí | David | 8.42032 | -82.41947 | 2016 | 0 | 1 |
| CH7_3 | Chiriquí | David | 8.41800 | -82.42690 | 2016 | 0 | 0 |
| CH7_4 | Chiriquí | David | 8.41570 | -82.42717 | 2016 | 0 | 0 |
| CH7_5 | Chiriquí | David | 8.41253 | -82.42680 | 2016 | 0 | 0 |
| CH7_6 | Chiriquí | David | 8.41150 | -82.42918 | 2016 | 0 | 0 |
| CH7_7 | Chiriquí | David | 8.41515 | -82.42934 | 2016 | 0 | 0 |
| CH7_8 | Chiriquí | David | 8.41849 | -82.42971 | 2016 | 0 | 0 |
| CH7_9 | Chiriquí | David | 8.42098 | -82.42838 | 2016 | 1 | 0 |
| CL1_1 | Coclé | Copé | 8.62065 | -80.57953 | 2016 | 1 | 1 |
| CL1_10 | Coclé | Copé | 8.61979 | -80.58417 | 2016 | 0 | 0 |
| CL1_11 | Coclé | Copé | 8.62126 | -80.58864 | 2016 | 0 | 0 |
| CL1_12 | Coclé | Copé | 8.62100 | -80.59002 | 2016 | 0 | 0 |
| CL1_13 | Coclé | Copé | 8.61803 | -80.58942 | 2016 | 0 | 0 |
| CL1_14 | Coclé | Copé | 8.62024 | -80.58998 | 2016 | 0 | 0 |
| CL1_15 | Coclé | Copé | 8.61813 | -80.58151 | 2016 | 1 | 1 |
| CL1_16 | Coclé | Copé | 8.61891 | -80.57971 | 2016 | 0 | 0 |
| CL1_17 | Coclé | Copé | 8.61877 | -80.57926 | 2016 | 0 | 0 |
| CL1_18 | Coclé | Copé | 8.61948 | -80.57762 | 2016 | 0 | 1 |
| CL1_19 | Coclé | Copé | 8.61943 | -80.57666 | 2016 | 1 | 0 |
| CL1_2 | Coclé | Copé | 8.62123 | -80.57967 | 2016 | 0 | 1 |
| CL1_20 | Coclé | Copé | 8.62061 | -80.57609 | 2016 | 0 | 1 |
| CL1_3 | Coclé | Copé | 8.62352 | -80.58004 | 2016 | 0 | 1 |
| CL1_4 | Coclé | Copé | 8.62468 | -80.58094 | 2016 | 0 | 1 |
| CL1_5 | Coclé | Copé | 8.62148 | -80.57904 | 2016 | 0 | 0 |
| CL1_6 | Coclé | Copé | 8.61866 | -80.58061 | 2016 | 0 | 0 |
| CL1_7 | Coclé | Copé | 8.61894 | -80.58160 | 2016 | 0 | 0 |
| CL1_8 | Coclé | Copé | 8.61941 | -80.58271 | 2016 | 0 | 0 |
| CL1_9 | Coclé | Copé | 8.61904 | -80.58234 | 2016 | 1 | 0 |
| CN1_1 | Colón | Miguel De La Borda | 9.15222 | -80.30639 | 2016 | 1 | 0 |
| CN1_10 | Colón | Miguel De La Borda | 9.15389 | -80.30556 | 2016 | 1 | 0 |
| CN1_11 | Colón | Miguel De La Borda | 9.15472 | -80.30500 | 2016 | 0 | 0 |
| CN1_12 | Colón | Miguel De La Borda | 9.15444 | -80.30417 | 2016 | 0 | 0 |
| CN1_13 | Colón | Miguel De La Borda | 9.15528 | -80.30083 | 2016 | 0 | 0 |
| CN1_14 | Colón | Miguel De La Borda | 9.15500 | -80.30167 | 2016 | 0 | 0 |

| | | | | | | | |
|---|---|---|---|---|---|---|---|
| CN1_15 | Colón | Miguel De La Borda | 9.15500 | -80.30306 | 2016 | 0 | 0 |
| CN1_16 | Colón | Miguel De La Borda | 9.15528 | -80.30444 | 2016 | 0 | 0 |
| CN1_2 | Colón | Miguel De La Borda | 9.15194 | -80.30639 | 2016 | 0 | 0 |
| CN1_3 | Colón | Miguel De La Borda | 9.15278 | -80.30639 | 2016 | 1 | 0 |
| CN1_4 | Colón | Miguel De La Borda | 9.15278 | -80.30583 | 2016 | 0 | 0 |
| CN1_5 | Colón | Miguel De La Borda | 9.15222 | -80.30556 | 2016 | 0 | 0 |
| CN1_6 | Colón | Miguel De La Borda | 9.15417 | -80.30611 | 2016 | 0 | 0 |
| CN1_7 | Colón | Miguel De La Borda | 9.15250 | -80.30500 | 2016 | 0 | 0 |
| CN1_8 | Colón | Miguel De La Borda | 9.15278 | -80.30500 | 2016 | 0 | 0 |
| CN1_9 | Colón | Miguel De La Borda | 9.15361 | -80.30417 | 2016 | 0 | 0 |
| CN2_1 | Colón | Gobea | 9.16722 | -80.25472 | 2016 | 0 | 0 |
| CN2_2 | Colón | Gobea | 9.16778 | -80.25472 | 2016 | 0 | 0 |
| CN2_6 | Colón | Gobea | 9.16778 | -80.25056 | 2016 | 0 | 0 |
| CN2_7 | Colón | Gobea | 9.16778 | -80.25194 | 2016 | 0 | 0 |
| CN2_8 | Colón | Gobea | 9.16750 | -80.25306 | 2016 | 0 | 0 |
| CN3_1 | Colón | Portobelo | 7.76694 | -80.27278 | 2016 | 0 | 0 |
| CN3_10 | Colón | Portobelo | 9.55333 | -79.65667 | 2016 | 0 | 0 |
| CN3_11 | Colón | Portobelo | 9.55167 | -79.65528 | 2016 | 0 | 0 |
| CN3_12 | Colón | Portobelo | 9.55278 | -79.65500 | 2016 | 0 | 0 |
| CN3_13 | Colón | Portobelo | 9.55389 | -79.65139 | 2016 | 0 | 0 |
| CN3_14 | Colón | Portobelo | 9.55472 | -79.65083 | 2016 | 0 | 0 |
| CN3_15 | Colón | Portobelo | 9.55306 | -79.65000 | 2016 | 0 | 0 |
| CN3_2 | Colón | Portobelo | 9.55583 | -79.65194 | 2016 | 0 | 0 |
| CN3_3 | Colón | Portobelo | 9.55472 | -79.65139 | 2016 | 0 | 0 |
| CN3_4 | Colón | Portobelo | 9.55444 | -79.65306 | 2016 | 0 | 0 |
| CN3_5 | Colón | Portobelo | 9.55472 | -79.65389 | 2016 | 0 | 0 |
| CN3_6 | Colón | Portobelo | 9.55444 | -79.65500 | 2016 | 1 | 0 |
| CN3_7 | Colón | Portobelo | 9.55583 | -79.65500 | 2016 | 1 | 0 |
| CN3_8 | Colón | Portobelo | 9.55389 | -79.65861 | 2016 | 0 | 0 |
| CN3_9 | Colón | Portobelo | 9.55361 | -79.65750 | 2016 | 1 | 0 |
| CN4_1 | Colón | Nuevo Tonosí | 9.55111 | -79.63778 | 2016 | 0 | 0 |
| CN4_2 | Colón | Nuevo Tonosí | 9.55278 | -79.63972 | 2016 | 0 | 0 |
| CN4_3 | Colón | Nuevo Tonosí | 9.55250 | -79.64250 | 2016 | 0 | 0 |
| CN4_4 | Colón | Nuevo Tonosí | 9.55194 | -79.66556 | 2016 | 0 | 0 |
| CN4_5 | Colón | Nuevo Tonosí | 9.54833 | -79.67000 | 2016 | 0 | 0 |
| CN5_1 | Colón | Sabanitas | 9.35444 | -79.80222 | 2016 | 0 | 0 |
| CN5_10 | Colón | Sabanitas | 9.34694 | -79.80056 | 2016 | 0 | 1 |
| CN5_11 | Colón | Sabanitas | 9.34833 | -79.80083 | 2016 | 0 | 0 |
| CN5_12 | Colón | Sabanitas | 9.35444 | -79.81000 | 2016 | 1 | 0 |
| CN5_13 | Colón | Sabanitas | 9.35361 | -79.80861 | 2016 | 1 | 0 |
| CN5_14 | Colón | Sabanitas | 9.35250 | -79.80528 | 2016 | 0 | 0 |
| CN5_15 | Colón | Sabanitas | 9.35417 | -79.80778 | 2016 | 1 | 0 |
| CN5_16 | Colón | Sabanitas | 9.35528 | -79.80917 | 2016 | 1 | 0 |

| | | | | | | | |
|---|---|---|---|---|---|---|---|
| CN5_17 | Colón | Sabanitas | 9.35639 | -79.80667 | 2016 | 1 | 0 |
| CN5_18 | Colón | Sabanitas | 9.35194 | -79.80667 | 2016 | 1 | 0 |
| CN5_19 | Colón | Sabanitas | 9.35000 | -79.80806 | 2016 | 0 | 0 |
| CN5_2 | Colón | Sabanitas | 9.35417 | -79.80167 | 2016 | 1 | 0 |
| CN5_20 | Colón | Sabanitas | 9.34944 | -79.80778 | 2016 | 0 | 1 |
| CN5_3 | Colón | Sabanitas | 9.35389 | -79.80139 | 2016 | 1 | 1 |
| CN5_4 | Colón | Sabanitas | 9.35250 | -79.80250 | 2016 | 1 | 0 |
| CN5_5 | Colón | Sabanitas | 9.35167 | -79.80083 | 2016 | 1 | 0 |
| CN5_6 | Colón | Sabanitas | 9.35083 | -79.79833 | 2016 | 0 | 1 |
| CN5_7 | Colón | Sabanitas | 9.35167 | -79.79750 | 2016 | 0 | 1 |
| CN5_8 | Colón | Sabanitas | 9.35028 | -79.79750 | 2016 | 0 | 0 |
| CN5_9 | Colón | Sabanitas | 9.35028 | -79.79889 | 2016 | 1 | 1 |
| CN6_1 | Colón | Gamboa | 9.11867 | -79.69586 | 2016 | 0 | 0 |
| CN6_2 | Colón | Gamboa | 9.11967 | -79.69928 | 2016 | 0 | 0 |
| CN6_3 | Colón | Gamboa | 9.11967 | -79.69928 | 2016 | 0 | 0 |
| CN6_4 | Colón | Gamboa | 9.11967 | -79.69928 | 2016 | 0 | 0 |
| CN6_5 | Colón | Gamboa | 9.11867 | -79.69586 | 2016 | 0 | 0 |
| CN6_6 | Colón | Gamboa | 9.11867 | -79.69586 | 2016 | 0 | 0 |
| CN6_7 | Colón | Gamboa | 9.11867 | -79.69586 | 2016 | 0 | 0 |
| CN6_8 | Colón | Gamboa | 9.11786 | -79.69614 | 2016 | 0 | 0 |
| CN6_9 | Colón | Gamboa | 9.11867 | -79.69586 | 2016 | 0 | 0 |
| CN6_10 | Colón | Gamboa | 9.11867 | -79.69586 | 2016 | 0 | 1 |
| CN6_11 | Colón | Gamboa | 9.11867 | -79.69586 | 2016 | 0 | 0 |
| CN6_12 | Colón | Gamboa | 9.11867 | -79.69586 | 2016 | 0 | 0 |
| CN6_13 | Colón | Gamboa | 9.11867 | -79.69586 | 2016 | 0 | 0 |
| CN6_15 | Colón | Gamboa | 9.11742 | -79.69344 | 2016 | 0 | 0 |
| CN6_16 | Colón | Gamboa | 9.11978 | -79.69344 | 2016 | 0 | 0 |
| CN6_17 | Colón | Gamboa | 9.11800 | -79.69633 | 2016 | 0 | 0 |
| CN6_18 | Colón | Gamboa | 9.11844 | -79.69603 | 2016 | 0 | 0 |
| CN6_19 | Colón | Gamboa | 9.11686 | -79.69822 | 2016 | 0 | 0 |
| CN6_20 | Colón | Gamboa | 9.11953 | -79.69942 | 2016 | 0 | 0 |
| CN6_21 | Colón | Gamboa | 9.11867 | -79.69586 | 2016 | 0 | 0 |
| CN6_22 | Colón | Gamboa | 9.11967 | -79.69928 | 2016 | 0 | 0 |
| CN6_23 | Colón | Gamboa | 9.11967 | -79.69928 | 2016 | 0 | 0 |
| CN6_24 | Colón | Gamboa | 9.11967 | -79.69928 | 2016 | 0 | 0 |
| CN6_25 | Colón | Gamboa | 9.11867 | -79.69586 | 2016 | 0 | 0 |
| CN6_26 | Colón | Gamboa | 9.11867 | -79.69586 | 2016 | 0 | 0 |
| CN6_27 | Colón | Gamboa | 9.11867 | -79.69586 | 2016 | 0 | 0 |
| CN6_28 | Colón | Gamboa | 9.11786 | -79.69614 | 2016 | 0 | 0 |
| CN6_29 | Colón | Gamboa | 9.11867 | -79.69586 | 2016 | 0 | 0 |
| CN6_30 | Colón | Gamboa | 9.11867 | -79.69586 | 2016 | 0 | 0 |
| CN6_31 | Colón | Gamboa | 9.11867 | -79.69586 | 2016 | 0 | 0 |
| CN6_32 | Colón | Gamboa | 9.11867 | -79.69586 | 2016 | 0 | 0 |

| | | | | | | | |
|---|---|---|---|---|---|---|---|
| CN6_33 | Colón | Gamboa | 9.11867 | -79.69586 | 2016 | 0 | 0 |
| CN6_35 | Colón | Gamboa | 9.11742 | -79.69344 | 2016 | 0 | 0 |
| CN6_36 | Colón | Gamboa | 9.11978 | -79.69344 | 2016 | 0 | 0 |
| CN6_37 | Colón | Gamboa | 9.11800 | -79.69633 | 2016 | 0 | 0 |
| CN6_38 | Colón | Gamboa | 9.11844 | -79.69603 | 2016 | 0 | 0 |
| CN6_39 | Colón | Gamboa | 9.11686 | -79.69822 | 2016 | 0 | 0 |
| CN6_40 | Colón | Gamboa | 9.11953 | -79.69942 | 2016 | 0 | 0 |
| DA1_1 | Darién | Yaviza | 8.15444 | -77.69167 | 2016 | 1 | 0 |
| DA1_10 | Darién | Yaviza | 8.15750 | -77.69139 | 2016 | 1 | 0 |
| DA1_11 | Darién | Yaviza | 8.15917 | -77.69111 | 2016 | 1 | 0 |
| DA1_12 | Darién | Yaviza | 8.15972 | -77.69417 | 2016 | 1 | 0 |
| DA1_13 | Darién | Yaviza | 8.16028 | -77.69194 | 2016 | 1 | 0 |
| DA1_14 | Darién | Yaviza | 8.16111 | -77.69222 | 2016 | 1 | 0 |
| DA1_15 | Darién | Yaviza | 8.16083 | -77.69194 | 2016 | 1 | 0 |
| DA1_16 | Darién | Yaviza | 8.16000 | -77.69167 | 2016 | 1 | 0 |
| DA1_17 | Darién | Yaviza | 8.15694 | -77.69250 | 2016 | 1 | 0 |
| DA1_18 | Darién | Yaviza | 8.15750 | -77.69306 | 2016 | 1 | 0 |
| DA1_19 | Darién | Yaviza | 8.15639 | -77.69250 | 2016 | 1 | 0 |
| DA1_2 | Darién | Yaviza | 8.15556 | -77.69083 | 2016 | 1 | 0 |
| DA1_20 | Darién | Yaviza | 8.15639 | -77.69306 | 2016 | 1 | 0 |
| DA1_21 | Darién | Yaviza | 8.16528 | -77.69833 | 2016 | 1 | 0 |
| DA1_22 | Darién | Yaviza | 8.16556 | -77.69861 | 2016 | 1 | 0 |
| DA1_23 | Darién | Yaviza | 8.16339 | -77.69500 | 2016 | 1 | 0 |
| DA1_24 | Darién | Yaviza | 8.16750 | -77.69369 | 2016 | 0 | 0 |
| DA1_25 | Darién | Yaviza | 8.16667 | -77.69369 | 2016 | 0 | 0 |
| DA1_26 | Darién | Yaviza | 8.16556 | -77.69722 | 2016 | 1 | 0 |
| DA1_27 | Darién | Yaviza | 8.16611 | -77.69722 | 2016 | 0 | 0 |
| DA1_28 | Darién | Yaviza | 8.16583 | -77.69806 | 2016 | 0 | 0 |
| DA1_29 | Darién | Yaviza | 8.16472 | -77.69778 | 2016 | 0 | 0 |
| DA1_3 | Darién | Yaviza | 8.15611 | -77.69139 | 2016 | 1 | 0 |
| DA1_30 | Darién | Yaviza | 8.16417 | -77.69750 | 2016 | 0 | 0 |
| DA1_4 | Darién | Yaviza | 8.15528 | -77.69139 | 2016 | 1 | 0 |
| DA1_5 | Darién | Yaviza | 8.15500 | -77.69222 | 2016 | 0 | 0 |
| DA1_6 | Darién | Yaviza | 8.15722 | -77.69056 | 2016 | 1 | 0 |
| DA1_7 | Darién | Yaviza | 8.15750 | -77.69139 | 2016 | 1 | 0 |
| DA1_8 | Darién | Yaviza | 8.15833 | -77.69139 | 2016 | 1 | 0 |
| DA1_9 | Darién | Yaviza | 8.15778 | -77.69194 | 2016 | 1 | 0 |
| DA2_1 | Darién | Metetí | 8.50972 | -77.98056 | 2016 | 1 | 0 |
| DA2_10 | Darién | Metetí | 8.49917 | -77.98000 | 2016 | 1 | 0 |
| DA2_11 | Darién | Metetí | 8.49750 | -77.98111 | 2016 | 1 | 0 |
| DA2_12 | Darién | Metetí | 8.49833 | -77.98167 | 2016 | 0 | 0 |
| DA2_13 | Darién | Metetí | 8.49750 | -77.98139 | 2016 | 1 | 0 |
| DA2_14 | Darién | Metetí | 8.49889 | -77.98222 | 2016 | 0 | 0 |

| | | | | | | | |
|---|---|---|---|---|---|---|---|
| DA2_15 | Darién | Metetí | 8.49972 | -77.98306 | 2016 | 0 | 0 |
| DA2_16 | Darién | Metetí | 8.50056 | -77.98333 | 2016 | 0 | 0 |
| DA2_17 | Darién | Metetí | 8.50083 | -77.98278 | 2016 | 0 | 0 |
| DA2_18 | Darién | Metetí | 8.49972 | -77.98222 | 2016 | 0 | 0 |
| DA2_19 | Darién | Metetí | 8.50028 | -77.98139 | 2016 | 0 | 0 |
| DA2_2 | Darién | Metetí | 8.15556 | -77.69083 | 2016 | 1 | 0 |
| DA2_20 | Darién | Metetí | 8.49500 | -77.98167 | 2016 | 0 | 0 |
| DA2_21 | Darién | Metetí | 8.49444 | -77.98111 | 2016 | 0 | 0 |
| DA2_22 | Darién | Metetí | 8.49389 | -77.98028 | 2016 | 0 | 0 |
| DA2_23 | Darién | Metetí | 8.49306 | -77.98028 | 2016 | 0 | 0 |
| DA2_24 | Darién | Metetí | 8.49361 | -77.98139 | 2016 | 0 | 0 |
| DA2_25 | Darién | Metetí | 8.49250 | -77.98194 | 2016 | 0 | 0 |
| DA2_3 | Darién | Metetí | 8.50222 | -77.98056 | 2016 | 1 | 0 |
| DA2_4 | Darién | Metetí | 8.50139 | -77.97972 | 2016 | 1 | 0 |
| DA2_5 | Darién | Metetí | 8.50167 | -77.97833 | 2016 | 1 | 0 |
| DA2_6 | Darién | Metetí | 8.50222 | -77.97750 | 2016 | 1 | 0 |
| DA2_7 | Darién | Metetí | 8.50194 | -77.97778 | 2016 | 0 | 0 |
| DA2_8 | Darién | Metetí | 8.50139 | -77.97861 | 2016 | 0 | 0 |
| DA2_9 | Darién | Metetí | 8.48444 | -77.97917 | 2016 | 1 | 0 |
| DA3_1 | Darién | Bijagual | 8.45000 | -78.00139 | 2016 | 0 | 0 |
| DA3_2 | Darién | Bijagual | 8.45028 | -78.00333 | 2016 | 0 | 0 |
| DA3_3 | Darién | Bijagual | 8.45028 | -78.00306 | 2016 | 0 | 0 |
| DA3_4 | Darién | Bijagual | 8.45056 | -78.00361 | 2016 | 0 | 0 |
| DA3_5 | Darién | Bijagual | 8.45056 | -78.00111 | 2016 | 0 | 0 |
| DA3_6 | Darién | Bijagual | 8.45111 | -78.00361 | 2016 | 0 | 0 |
| DA3_7 | Darién | Bijagual | 8.45083 | -78.00417 | 2016 | 0 | 0 |
| DA3_8 | Darién | Bijagual | 8.45139 | -78.00444 | 2016 | 0 | 0 |
| DA4_1 | Darién | Bajo Iglesias | 8.39778 | -78.00861 | 2016 | 1 | 0 |
| DA4_10 | Darién | Bajo Iglesias | 8.39583 | -78.01361 | 2016 | 0 | 0 |
| DA4_11 | Darién | Bajo Iglesias | 8.69369 | -78.01139 | 2016 | 0 | 0 |
| DA4_12 | Darién | Bajo Iglesias | 8.39583 | -78.01139 | 2016 | 1 | 0 |
| DA4_13 | Darién | Bajo Iglesias | 8.39528 | -78.01417 | 2016 | 0 | 0 |
| DA4_14 | Darién | Bajo Iglesias | 8.39667 | -78.01056 | 2016 | 1 | 0 |
| DA4_15 | Darién | Bajo Iglesias | 8.39583 | -78.01028 | 2016 | 0 | 0 |
| DA4_2 | Darién | Bajo Iglesias | 8.39861 | -78.00917 | 2016 | 1 | 0 |
| DA4_3 | Darién | Bajo Iglesias | 8.39917 | -78.00861 | 2016 | 0 | 0 |
| DA4_4 | Darién | Bajo Iglesias | 8.40056 | -78.00694 | 2016 | 1 | 0 |
| DA4_5 | Darién | Bajo Iglesias | 8.39833 | -78.00944 | 2016 | 1 | 0 |
| DA4_6 | Darién | Bajo Iglesias | 8.39778 | -78.01194 | 2016 | 0 | 0 |
| DA4_7 | Darién | Bajo Iglesias | 8.39833 | -78.01250 | 2016 | 0 | 0 |
| DA4_8 | Darién | Bajo Iglesias | 8.39917 | -78.01306 | 2016 | 0 | 0 |
| DA4_9 | Darién | Bajo Iglesias | 8.39556 | -78.01528 | 2016 | 0 | 0 |
| LS1_1 | Los Santos | La Villa | 7.93833 | -80.41694 | 2016 | 1 | 0 |

| | | | | | | | |
|---|---|---|---|---|---|---|---|
| LS1_10 | Los Santos | La Villa | 7.93194 | -80.41722 | 2016 | 0 | 0 |
| LS1_11 | Los Santos | La Villa | 7.93306 | -80.41500 | 2016 | 0 | 1 |
| LS1_12 | Los Santos | La Villa | 7.93556 | -80.41583 | 2016 | 1 | 1 |
| LS1_13 | Los Santos | La Villa | 7.93583 | -80.41361 | 2016 | 1 | 1 |
| LS1_14 | Los Santos | La Villa | 7.93639 | -80.41250 | 2016 | 1 | 0 |
| LS1_15 | Los Santos | La Villa | 7.93861 | -80.41167 | 2016 | 1 | 0 |
| LS1_16 | Los Santos | La Villa | 7.93528 | -80.41194 | 2016 | 1 | 1 |
| LS1_17 | Los Santos | La Villa | 7.93389 | -80.41167 | 2016 | 1 | 0 |
| LS1_18 | Los Santos | La Villa | 7.93444 | -80.41417 | 2016 | 1 | 1 |
| LS1_19 | Los Santos | La Villa | 7.93306 | -80.41333 | 2016 | 1 | 0 |
| LS1_2 | Los Santos | La Villa | 7.93917 | -80.41667 | 2016 | 1 | 0 |
| LS1_20 | Los Santos | La Villa | 7.93167 | -80.41000 | 2016 | 1 | 1 |
| LS1_3 | Los Santos | La Villa | 7.94139 | -80.41111 | 2016 | 1 | 1 |
| LS1_4 | Los Santos | La Villa | 7.93806 | -80.41361 | 2016 | 1 | 0 |
| LS1_5 | Los Santos | La Villa | 7.93750 | -80.41472 | 2016 | 1 | 1 |
| LS1_6 | Los Santos | La Villa | 7.93639 | -80.41583 | 2016 | 0 | 0 |
| LS1_7 | Los Santos | La Villa | 7.93361 | -80.41833 | 2016 | 0 | 0 |
| LS1_8 | Los Santos | La Villa | 7.93556 | -80.42222 | 2016 | 0 | 1 |
| LS1_9 | Los Santos | La Villa | 7.93222 | -80.41917 | 2016 | 0 | 1 |
| LS2_1 | Los Santos | Paritilla | 7.63166 | -80.16861 | 2016 | 0 | 0 |
| LS2_10 | Los Santos | Paritilla | 7.62916 | -80.17889 | 2016 | 1 | 1 |
| LS2_11 | Los Santos | Paritilla | 7.62972 | -80.17611 | 2016 | 0 | 1 |
| LS2_12 | Los Santos | Paritilla | 7.63138 | -80.17111 | 2016 | 0 | 0 |
| LS2_13 | Los Santos | Paritilla | 7.63027 | -80.17250 | 2016 | 0 | 1 |
| LS2_14 | Los Santos | Paritilla | 7.62944 | -80.17417 | 2016 | 1 | 1 |
| LS2_15 | Los Santos | Paritilla | 7.62916 | -80.17167 | 2016 | 0 | 0 |
| LS2_16 | Los Santos | Paritilla | 7.63083 | -80.17028 | 2016 | 1 | 1 |
| LS2_17 | Los Santos | Paritilla | 7.63083 | -80.17167 | 2016 | 0 | 0 |
| LS2_18 | Los Santos | Paritilla | 7.63000 | -80.17389 | 2016 | 1 | 1 |
| LS2_19 | Los Santos | Paritilla | 7.63111 | -80.17472 | 2016 | 0 | 1 |
| LS2_2 | Los Santos | Paritilla | 7.63027 | -80.17028 | 2016 | 0 | 1 |
| LS2_20 | Los Santos | Paritilla | 7.62944 | -80.17472 | 2016 | 0 | 1 |
| LS2_3 | Los Santos | Paritilla | 7.62916 | -80.17056 | 2016 | 0 | 0 |
| LS2_4 | Los Santos | Paritilla | 7.62833 | -80.16972 | 2016 | 0 | 1 |
| LS2_5 | Los Santos | Paritilla | 7.62805 | -80.17083 | 2016 | 0 | 1 |
| LS2_6 | Los Santos | Paritilla | 7.62888 | -80.17278 | 2016 | 1 | 1 |
| LS2_7 | Los Santos | Paritilla | 7.62888 | -80.17389 | 2016 | 1 | 1 |
| LS2_8 | Los Santos | Paritilla | 7.62833 | -80.17639 | 2016 | 0 | 1 |
| LS2_9 | Los Santos | Paritilla | 7.62833 | -80.17722 | 2016 | 0 | 1 |
| LS3_1 | Los Santos | Tonosí | 7.40500 | -80.44306 | 2016 | 0 | 0 |
| LS3_10 | Los Santos | Tonosí | 7.41000 | -80.43528 | 2016 | 0 | 1 |
| LS3_11 | Los Santos | Tonosí | 7.41222 | -80.43583 | 2016 | 0 | 1 |
| LS3_12 | Los Santos | Tonosí | 7.40972 | -80.42111 | 2016 | 0 | 1 |

| | | | | | | | |
|---|---|---|---|---|---|---|---|
| LS3_13 | Los Santos | Tonosí | 7.40917 | -80.44000 | 2016 | 0 | 0 |
| LS3_14 | Los Santos | Tonosí | 7.41056 | -80.44278 | 2016 | 0 | 1 |
| LS3_15 | Los Santos | Tonosí | 7.40722 | -80.44278 | 2016 | 0 | 1 |
| LS3_16 | Los Santos | Tonosí | 7.40667 | -80.44194 | 2016 | 0 | 1 |
| LS3_17 | Los Santos | Tonosí | 7.40444 | -80.44139 | 2016 | 0 | 1 |
| LS3_18 | Los Santos | Tonosí | 7.40444 | -80.43806 | 2016 | 0 | 0 |
| LS3_19 | Los Santos | Tonosí | 7.40750 | -80.43750 | 2016 | 0 | 1 |
| LS3_2 | Los Santos | Tonosí | 7.40472 | -80.44083 | 2016 | 0 | 1 |
| LS3_20 | Los Santos | Tonosí | 7.40833 | -80.43778 | 2016 | 0 | 1 |
| LS3_3 | Los Santos | Tonosí | 7.40472 | -80.43917 | 2016 | 0 | 1 |
| LS3_4 | Los Santos | Tonosí | 7.40639 | -80.43806 | 2016 | 0 | 0 |
| LS3_5 | Los Santos | Tonosí | 7.40694 | -80.44111 | 2016 | 0 | 1 |
| LS3_6 | Los Santos | Tonosí | 7.40556 | -80.44000 | 2016 | 0 | 1 |
| LS3_7 | Los Santos | Tonosí | 7.40583 | -80.44000 | 2016 | 0 | 1 |
| LS3_8 | Los Santos | Tonosí | 7.40583 | -80.43556 | 2016 | 0 | 1 |
| LS3_9 | Los Santos | Tonosí | 7.40778 | -80.43583 | 2016 | 0 | 0 |
| LS4_1 | Los Santos | El Cacao | 7.44194 | -80.41139 | 2016 | 0 | 1 |
| LS4_10 | Los Santos | El Cacao | 7.44667 | -80.40444 | 2016 | 0 | 1 |
| LS4_11 | Los Santos | El Cacao | 7.45028 | -80.40389 | 2016 | 0 | 1 |
| LS4_12 | Los Santos | El Cacao | 7.45194 | -80.40306 | 2016 | 0 | 1 |
| LS4_13 | Los Santos | El Cacao | 7.45639 | -80.40333 | 2016 | 0 | 0 |
| LS4_14 | Los Santos | El Cacao | 7.45250 | -80.40167 | 2016 | 0 | 0 |
| LS4_15 | Los Santos | El Cacao | 7.45528 | -80.40028 | 2016 | 0 | 0 |
| LS4_16 | Los Santos | El Cacao | 7.44972 | -80.40028 | 2016 | 0 | 1 |
| LS4_17 | Los Santos | El Cacao | 7.45472 | -80.40028 | 2016 | 0 | 1 |
| LS4_18 | Los Santos | El Cacao | 7.45639 | -80.40056 | 2016 | 0 | 1 |
| LS4_19 | Los Santos | El Cacao | 7.45750 | -80.40111 | 2016 | 0 | 1 |
| LS4_2 | Los Santos | El Cacao | 7.44361 | -80.41111 | 2016 | 0 | 1 |
| LS4_20 | Los Santos | El Cacao | 7.45889 | -80.40139 | 2016 | 0 | 0 |
| LS4_3 | Los Santos | El Cacao | 7.44500 | -80.40944 | 2016 | 0 | 1 |
| LS4_4 | Los Santos | El Cacao | 7.44583 | -80.40833 | 2016 | 0 | 1 |
| LS4_5 | Los Santos | El Cacao | 7.44861 | -80.40611 | 2016 | 0 | 1 |
| LS4_6 | Los Santos | El Cacao | 7.44833 | -80.40361 | 2016 | 0 | 1 |
| LS4_7 | Los Santos | El Cacao | 7.45611 | -80.40528 | 2016 | 0 | 1 |
| LS4_8 | Los Santos | El Cacao | 7.45611 | -80.40528 | 2016 | 0 | 0 |
| LS4_9 | Los Santos | El Cacao | 7.44861 | -80.40444 | 2016 | 0 | 1 |
| LS6_1 | Los Santos | Macaracas | 7.74000 | -80.54694 | 2016 | 0 | 1 |
| LS6_10 | Los Santos | Macaracas | 7.72861 | -80.55139 | 2016 | 0 | 0 |
| LS6_11 | Los Santos | Macaracas | 7.72944 | -80.55056 | 2016 | 0 | 0 |
| LS6_12 | Los Santos | Macaracas | 7.73000 | -80.54806 | 2016 | 0 | 0 |
| LS6_13 | Los Santos | Macaracas | 7.73194 | -80.55000 | 2016 | 0 | 1 |
| LS6_14 | Los Santos | Macaracas | 7.73333 | -80.54944 | 2016 | 0 | 1 |
| LS6_15 | Los Santos | Macaracas | 7.73333 | -80.55167 | 2016 | 0 | 1 |

| | | | | | | | |
|---|---|---|---|---|---|---|---|
| LS6_16 | Los Santos | Macaracas | 7.73556 | -80.55000 | 2016 | 0 | 0 |
| LS6_17 | Los Santos | Macaracas | 7.73583 | -80.54806 | 2016 | 0 | 0 |
| LS6_18 | Los Santos | Macaracas | 7.73778 | -80.54972 | 2016 | 0 | 0 |
| LS6_19 | Los Santos | Macaracas | 7.74056 | -80.54917 | 2016 | 0 | 0 |
| LS6_2 | Los Santos | Macaracas | 7.73806 | -80.54861 | 2016 | 0 | 1 |
| LS6_20 | Los Santos | Macaracas | 7.74250 | -80.54972 | 2016 | 0 | 0 |
| LS6_3 | Los Santos | Macaracas | 7.73667 | -80.54972 | 2016 | 1 | 1 |
| LS6_4 | Los Santos | Macaracas | 7.73472 | -80.55167 | 2016 | 0 | 1 |
| LS6_5 | Los Santos | Macaracas | 7.73361 | -80.55222 | 2016 | 0 | 1 |
| LS6_6 | Los Santos | Macaracas | 7.73194 | -80.55250 | 2016 | 0 | 1 |
| LS6_7 | Los Santos | Macaracas | 7.72972 | -80.55306 | 2016 | 0 | 1 |
| LS6_8 | Los Santos | Macaracas | 7.72750 | -80.55194 | 2016 | 0 | 0 |
| LS6_9 | Los Santos | Macaracas | 7.72639 | -80.55111 | 2016 | 0 | 1 |
| PA1_1 | Panamá Oeste | Princesa Mía | 8.96444 | -79.70417 | 2016 | 0 | 0 |
| PA1_10 | Panamá Oeste | Princesa Mía | 8.96552 | -79.70322 | 2016 | 1 | 1 |
| PA1_11 | Panamá Oeste | Princesa Mía | 8.96521 | -79.70266 | 2016 | 0 | 1 |
| PA1_12 | Panamá Oeste | Princesa Mía | 8.96433 | -79.70174 | 2016 | 0 | 1 |
| PA1_13 | Panamá Oeste | Princesa Mía | 8.96574 | -79.70148 | 2016 | 0 | 1 |
| PA1_14 | Panamá Oeste | Princesa Mía | 8.96505 | -79.70087 | 2016 | 0 | 0 |
| PA1_15 | Panamá Oeste | Princesa Mía | 8.96548 | -79.70061 | 2016 | 1 | 0 |
| PA1_16 | Panamá Oeste | Princesa Mía | 8.96604 | -79.70110 | 2016 | 1 | 0 |
| PA1_2 | Panamá Oeste | Princesa Mía | 8.96556 | -79.70361 | 2016 | 0 | 1 |
| PA1_3 | Panamá Oeste | Princesa Mía | 8.96472 | -79.70306 | 2016 | 1 | 1 |
| PA1_4 | Panamá Oeste | Princesa Mía | 8.96556 | -79.70250 | 2016 | 1 | 1 |
| PA1_5 | Panamá Oeste | Princesa Mía | 8.96444 | -79.70111 | 2016 | 0 | 1 |
| PA1_6 | Panamá Oeste | Princesa Mía | 8.96611 | -79.70194 | 2016 | 0 | 0 |
| PA1_7 | Panamá Oeste | Princesa Mía | 8.96528 | -79.70028 | 2016 | 0 | 1 |
| PA1_8 | Panamá Oeste | Princesa Mía | 8.96528 | -79.70419 | 2016 | 0 | 1 |
| PA1_9 | Panamá Oeste | Princesa Mía | 8.96454 | -79.70370 | 2016 | 0 | 1 |
| PA2_1 | Panamá Oeste | Lluvia de Oro | 8.95844 | -79.70000 | 2016 | 1 | 1 |
| PA2_10 | Panamá Oeste | Lluvia de Oro | 8.95925 | -79.70103 | 2016 | 1 | 0 |
| PA2_11 | Panamá Oeste | Lluvia de Oro | 8.96088 | -79.70145 | 2016 | 0 | 0 |
| PA2_12 | Panamá Oeste | Lluvia de Oro | 8.95888 | -79.70048 | 2016 | 0 | 0 |
| PA2_13 | Panamá Oeste | Lluvia de Oro | 8.96059 | -79.70040 | 2016 | 1 | 1 |
| PA2_14 | Panamá Oeste | Lluvia de Oro | 8.96051 | -79.69973 | 2016 | 0 | 0 |
| PA2_15 | Panamá Oeste | Lluvia de Oro | 8.95935 | -79.69937 | 2016 | 0 | 0 |
| PA2_16 | Panamá Oeste | Lluvia de Oro | 8.96081 | -79.69877 | 2016 | 0 | 0 |
| PA2_17 | Panamá Oeste | Lluvia de Oro | 8.95944 | -79.69801 | 2016 | 0 | 0 |
| PA2_18 | Panamá Oeste | Lluvia de Oro | 8.96011 | -79.69734 | 2016 | 1 | 1 |
| PA2_19 | Panamá Oeste | Lluvia de Oro | 8.95927 | -79.69621 | 2016 | 0 | 1 |
| PA2_2 | Panamá Oeste | Lluvia de Oro | 8.95931 | -79.69944 | 2016 | 1 | 1 |
| PA2_3 | Panamá Oeste | Lluvia de Oro | 8.96083 | -79.69861 | 2016 | 1 | 1 |
| PA2_4 | Panamá Oeste | Lluvia de Oro | 8.95950 | -79.69858 | 2016 | 0 | 1 |

| | | | | | | | |
|---|---|---|---|---|---|---|---|
| PA2_5 | Panamá Oeste | Lluvia de Oro | 8.96050 | -79.69786 | 2016 | 0 | 0 |
| PA2_6 | Panamá Oeste | Lluvia de Oro | 8.95942 | -79.69761 | 2016 | 0 | 1 |
| PA2_7 | Panamá Oeste | Lluvia de Oro | 8.96033 | -79.69708 | 2016 | 1 | 1 |
| PA2_8 | Panamá Oeste | Lluvia de Oro | 8.95989 | -79.70200 | 2016 | 0 | 1 |
| PA2_9 | Panamá Oeste | Lluvia de Oro | 8.96036 | -79.70206 | 2016 | 0 | 0 |
| PA3_1 | Panamá Oeste | Nuevo Chorrillo | 8.95206 | -79.69908 | 2016 | 1 | 1 |
| PA3_10 | Panamá Oeste | Nuevo Chorrillo | 8.95534 | -79.69649 | 2016 | 1 | 0 |
| PA3_11 | Panamá Oeste | Nuevo Chorrillo | 8.95289 | -79.69631 | 2016 | 0 | 0 |
| PA3_12 | Panamá Oeste | Nuevo Chorrillo | 8.95207 | -79.69725 | 2016 | 1 | 0 |
| PA3_13 | Panamá Oeste | Nuevo Chorrillo | 8.95344 | -79.69570 | 2016 | 1 | 0 |
| PA3_14 | Panamá Oeste | Nuevo Chorrillo | 8.95595 | -79.69533 | 2016 | 1 | 1 |
| PA3_15 | Panamá Oeste | Nuevo Chorrillo | 8.95403 | -79.69458 | 2016 | 1 | 0 |
| PA3_16 | Panamá Oeste | Nuevo Chorrillo | 8.95373 | -79.69303 | 2016 | 0 | 0 |
| PA3_17 | Panamá Oeste | Nuevo Chorrillo | 8.95650 | -79.69417 | 2016 | 1 | 0 |
| PA3_18 | Panamá Oeste | Nuevo Chorrillo | 8.95246 | -79.69097 | 2016 | 0 | 0 |
| PA3_19 | Panamá Oeste | Nuevo Chorrillo | 8.95717 | -79.69293 | 2016 | 1 | 1 |
| PA3_2 | Panamá Oeste | Nuevo Chorrillo | 8.95367 | -79.69803 | 2016 | 0 | 0 |
| PA3_3 | Panamá Oeste | Nuevo Chorrillo | 8.95289 | -79.69631 | 2016 | 0 | 1 |
| PA3_4 | Panamá Oeste | Nuevo Chorrillo | 8.95217 | -79.69664 | 2016 | 1 | 1 |
| PA3_5 | Panamá Oeste | Nuevo Chorrillo | 8.95578 | -79.69583 | 2016 | 1 | 1 |
| PA3_6 | Panamá Oeste | Nuevo Chorrillo | 8.95272 | -79.69494 | 2016 | 1 | 1 |
| PA3_7 | Panamá Oeste | Nuevo Chorrillo | 8.95008 | -79.69175 | 2016 | 1 | 1 |
| PA3_8 | Panamá Oeste | Nuevo Chorrillo | 8.95239 | -79.69902 | 2016 | 0 | 0 |
| PA3_9 | Panamá Oeste | Nuevo Chorrillo | 8.95389 | -79.69785 | 2016 | 0 | 0 |
| PA5_1 | Panamá | Chilibre | 9.15616 | -79.61788 | 2016 | 0 | 0 |
| PA5_10 | Panamá | Chilibre | 9.14948 | -79.61697 | 2016 | 0 | 0 |
| PA5_11 | Panamá | Chilibre | 9.14828 | -79.61978 | 2016 | 0 | 0 |
| PA5_12 | Panamá | Chilibre | 9.14771 | -79.61732 | 2016 | 0 | 1 |
| PA5_13 | Panamá | Chilibre | 9.14618 | -79.61881 | 2016 | 0 | 0 |
| PA5_14 | Panamá | Chilibre | 9.14519 | -79.61664 | 2016 | 0 | 0 |
| PA5_15 | Panamá | Chilibre | 9.14309 | -79.61678 | 2016 | 0 | 0 |
| PA5_16 | Panamá | Chilibre | 9.14446 | -79.61326 | 2016 | 0 | 0 |
| PA5_17 | Panamá | Chilibre | 9.14187 | -79.61387 | 2016 | 0 | 0 |
| PA5_18 | Panamá | Chilibre | 9.14023 | -79.61479 | 2016 | 0 | 1 |
| PA5_19 | Panamá | Chilibre | 9.13892 | -79.61627 | 2016 | 0 | 0 |
| PA5_2 | Panamá | Chilibre | 9.15359 | -79.61896 | 2016 | 0 | 0 |
| PA5_20 | Panamá | Chilibre | 9.13814 | -79.61518 | 2016 | 0 | 0 |
| PA5_3 | Panamá | Chilibre | 9.15376 | -79.62108 | 2016 | 0 | 0 |
| PA5_4 | Panamá | Chilibre | 9.15690 | -79.62093 | 2016 | 0 | 0 |
| PA5_5 | Panamá | Chilibre | 9.15228 | -79.62144 | 2016 | 0 | 0 |
| PA5_6 | Panamá | Chilibre | 9.15205 | -79.61757 | 2016 | 0 | 1 |
| PA5_7 | Panamá | Chilibre | 9.15170 | -79.61542 | 2016 | 0 | 1 |
| PA5_8 | Panamá | Chilibre | 9.15055 | -79.61928 | 2016 | 0 | 1 |

| | | | | | | | |
|---|---|---|---|---|---|---|---|
| PA5_9 | Panamá | Chilibre | 9.14915 | -79.61461 | 2016 | 0 | 1 |
| VE1_1 | Veraguas | Santa Fe | 8.51691 | -81.07857 | 2016 | 0 | 0 |
| VE1_10 | Veraguas | Santa Fe | 8.51161 | -81.07525 | 2016 | 0 | 1 |
| VE1_11 | Veraguas | Santa Fe | 8.51278 | -81.07591 | 2016 | 0 | 0 |
| VE1_12 | Veraguas | Santa Fe | 8.51405 | -81.07616 | 2016 | 0 | 0 |
| VE1_13 | Veraguas | Santa Fe | 8.51360 | -81.07701 | 2016 | 0 | 1 |
| VE1_14 | Veraguas | Santa Fe | 8.51378 | -81.07911 | 2016 | 0 | 1 |
| VE1_15 | Veraguas | Santa Fe | 8.51206 | -81.07781 | 2016 | 0 | 1 |
| VE1_16 | Veraguas | Santa Fe | 8.51341 | -81.07996 | 2016 | 0 | 0 |
| VE1_17 | Veraguas | Santa Fe | 8.51461 | -81.08039 | 2016 | 0 | 0 |
| VE1_18 | Veraguas | Santa Fe | 8.51185 | -81.07956 | 2016 | 0 | 1 |
| VE1_19 | Veraguas | Santa Fe | 8.51164 | -81.07922 | 2016 | 0 | 1 |
| VE1_2 | Veraguas | Santa Fe | 8.51526 | -81.07801 | 2016 | 0 | 1 |
| VE1_20 | Veraguas | Santa Fe | 8.51137 | -81.07823 | 2016 | 0 | 1 |
| VE1_3 | Veraguas | Santa Fe | 8.51434 | -81.07873 | 2016 | 0 | 1 |
| VE1_4 | Veraguas | Santa Fe | 8.51316 | -81.08071 | 2016 | 0 | 1 |
| VE1_5 | Veraguas | Santa Fe | 8.51165 | -81.08109 | 2016 | 0 | 1 |
| VE1_6 | Veraguas | Santa Fe | 8.51069 | -81.08093 | 2016 | 0 | 0 |
| VE1_7 | Veraguas | Santa Fe | 8.51007 | -81.07960 | 2016 | 0 | 1 |
| VE1_8 | Veraguas | Santa Fe | 8.50972 | -81.07755 | 2016 | 0 | 0 |
| VE1_9 | Veraguas | Santa Fe | 8.50882 | -81.07616 | 2016 | 0 | 0 |
| VE2_1 | Veraguas | San Francisco | 8.25223 | -80.97392 | 2016 | 0 | 1 |
| VE2_10 | Veraguas | San Francisco | 8.24794 | -80.97143 | 2016 | 0 | 1 |
| VE2_11 | Veraguas | San Francisco | 8.24770 | -80.96917 | 2016 | 0 | 1 |
| VE2_12 | Veraguas | San Francisco | 8.24581 | -80.97030 | 2016 | 0 | 1 |
| VE2_13 | Veraguas | San Francisco | 8.24599 | -80.97159 | 2016 | 0 | 0 |
| VE2_14 | Veraguas | San Francisco | 8.24656 | -80.97196 | 2016 | 0 | 1 |
| VE2_15 | Veraguas | San Francisco | 8.24727 | -80.97441 | 2016 | 0 | 1 |
| VE2_16 | Veraguas | San Francisco | 8.24727 | -80.97636 | 2016 | 0 | 1 |
| VE2_17 | Veraguas | San Francisco | 8.24506 | -80.97897 | 2016 | 0 | 0 |
| VE2_18 | Veraguas | San Francisco | 8.24665 | -80.97857 | 2016 | 0 | 1 |
| VE2_19 | Veraguas | San Francisco | 8.24545 | -80.97525 | 2016 | 0 | 0 |
| VE2_2 | Veraguas | San Francisco | 8.25104 | -80.97423 | 2016 | 0 | 1 |
| VE2_20 | Veraguas | San Francisco | 8.24310 | -80.97321 | 2016 | 0 | 0 |
| VE2_3 | Veraguas | San Francisco | 8.24916 | -80.97403 | 2016 | 0 | 1 |
| VE2_4 | Veraguas | San Francisco | 8.24884 | -80.97284 | 2016 | 0 | 1 |
| VE2_5 | Veraguas | San Francisco | 8.25028 | -80.97299 | 2016 | 0 | 0 |
| VE2_6 | Veraguas | San Francisco | 8.24890 | -80.97174 | 2016 | 0 | 1 |
| VE2_7 | Veraguas | San Francisco | 8.24967 | -80.97168 | 2016 | 0 | 1 |
| VE2_8 | Veraguas | San Francisco | 8.25162 | -80.97085 | 2016 | 0 | 1 |
| VE2_9 | Veraguas | San Francisco | 8.25015 | -80.97084 | 2016 | 0 | 1 |
| VE3_1 | Veraguas | Santiago | 8.12072 | -80.96609 | 2016 | 1 | 1 |
| VE3_10 | Veraguas | Santiago | 8.11873 | -80.95938 | 2016 | 1 | 1 |

| | | | | | | | |
|---|---|---|---|---|---|---|---|
| VE3_11 | Veraguas | Santiago | 8.11947 | -80.96127 | 2016 | 0 | 0 |
| VE3_12 | Veraguas | Santiago | 8.12100 | -80.96274 | 2016 | 0 | 0 |
| VE3_13 | Veraguas | Santiago | 8.12348 | -80.96317 | 2016 | 0 | 1 |
| VE3_14 | Veraguas | Santiago | 8.12038 | -80.96392 | 2016 | 0 | 0 |
| VE3_15 | Veraguas | Santiago | 8.11672 | -80.96396 | 2016 | 0 | 0 |
| VE3_16 | Veraguas | Santiago | 8.11436 | -80.96518 | 2016 | 0 | 0 |
| VE3_17 | Veraguas | Santiago | 8.11454 | -80.96314 | 2016 | 0 | 0 |
| VE3_18 | Veraguas | Santiago | 8.11185 | -80.96177 | 2016 | 0 | 1 |
| VE3_19 | Veraguas | Santiago | 8.11045 | -80.96380 | 2016 | 0 | 0 |
| VE3_2 | Veraguas | Santiago | 8.12247 | -80.96580 | 2016 | 0 | 1 |
| VE3_20 | Veraguas | Santiago | 8.11263 | -80.96606 | 2016 | 0 | 0 |
| VE3_3 | Veraguas | Santiago | 8.12405 | -80.96680 | 2016 | 0 | 0 |
| VE3_4 | Veraguas | Santiago | 8.12551 | -80.96575 | 2016 | 0 | 0 |
| VE3_5 | Veraguas | Santiago | 8.12748 | -80.96636 | 2016 | 0 | 0 |
| VE3_6 | Veraguas | Santiago | 8.12735 | -80.96462 | 2016 | 0 | 0 |
| VE3_7 | Veraguas | Santiago | 8.12567 | -80.96345 | 2016 | 0 | 1 |
| VE3_8 | Veraguas | Santiago | 8.12389 | -80.96136 | 2016 | 0 | 0 |
| VE3_9 | Veraguas | Santiago | 8.12132 | -80.95953 | 2016 | 0 | 0 |
| BTO1_1 | Bocas del Toro | Chiriquí Grande | 8.94372 | -82.11327 | 2017 | 0 | 1 |
| BTO1_10 | Bocas del Toro | Chiriquí Grande | 8.95063 | -82.11995 | 2017 | 0 | 0 |
| BTO1_11 | Bocas del Toro | Chiriquí Grande | 8.95148 | -82.12096 | 2017 | 1 | 0 |
| BTO1_12 | Bocas del Toro | Chiriquí Grande | 8.95219 | -82.12235 | 2017 | 0 | 0 |
| BTO1_13 | Bocas del Toro | Chiriquí Grande | 8.95239 | -82.12265 | 2017 | 0 | 0 |
| BTO1_14 | Bocas del Toro | Chiriquí Grande | 8.95215 | -82.12448 | 2017 | 0 | 0 |
| BTO1_15 | Bocas del Toro | Chiriquí Grande | 8.94714 | -82.11953 | 2017 | 0 | 0 |
| BTO1_16 | Bocas del Toro | Chiriquí Grande | 8.94778 | -82.12021 | 2017 | 0 | 0 |
| BTO1_17 | Bocas del Toro | Chiriquí Grande | 8.94867 | -82.12121 | 2017 | 0 | 0 |
| BTO1_18 | Bocas del Toro | Chiriquí Grande | 8.94967 | -82.12230 | 2017 | 0 | 0 |
| BTO1_19 | Bocas del Toro | Chiriquí Grande | 8.95077 | -82.12348 | 2017 | 0 | 0 |
| BTO1_2 | Bocas del Toro | Chiriquí Grande | 8.94524 | -82.11494 | 2017 | 0 | 0 |
| BTO1_20 | Bocas del Toro | Chiriquí Grande | 8.95182 | -82.12466 | 2017 | 0 | 0 |
| BTO1_3 | Bocas del Toro | Chiriquí Grande | 8.94751 | -82.11600 | 2017 | 0 | 0 |
| BTO1_4 | Bocas del Toro | Chiriquí Grande | 8.94807 | -82.11807 | 2017 | 0 | 0 |
| BTO1_5 | Bocas del Toro | Chiriquí Grande | 8.94768 | -82.11731 | 2017 | 1 | 0 |
| BTO1_6 | Bocas del Toro | Chiriquí Grande | 8.94979 | -82.11706 | 2017 | 1 | 0 |
| BTO1_7 | Bocas del Toro | Chiriquí Grande | 8.94938 | -82.11793 | 2017 | 0 | 0 |
| BTO1_8 | Bocas del Toro | Chiriquí Grande | 8.94939 | -82.11881 | 2017 | 0 | 0 |
| BTO1_9 | Bocas del Toro | Chiriquí Grande | 8.95003 | -82.11981 | 2017 | 1 | 0 |
| BTO2_1 | Bocas del Toro | Almirante | 9.29122 | -82.39593 | 2017 | 1 | 0 |
| BTO2_10 | Bocas del Toro | Almirante | 9.29428 | -82.39318 | 2017 | 1 | 0 |
| BTO2_11 | Bocas del Toro | Almirante | 9.29525 | -82.39678 | 2017 | 1 | 0 |
| BTO2_12 | Bocas del Toro | Almirante | 9.29685 | -82.39820 | 2017 | 1 | 0 |
| BTO2_13 | Bocas del Toro | Almirante | 9.29546 | -82.40397 | 2017 | 1 | 0 |

| | | | | | | | |
|---|---|---|---|---|---|---|---|
| BTO2_14 | Bocas del Toro | Almirante | 9.29575 | -82.40470 | 2017 | 1 | 0 |
| BTO2_15 | Bocas del Toro | Almirante | 9.29636 | -82.40483 | 2017 | 1 | 0 |
| BTO2_16 | Bocas del Toro | Almirante | 9.29631 | -82.40650 | 2017 | 0 | 0 |
| BTO2_17 | Bocas del Toro | Almirante | 9.29725 | -82.40908 | 2017 | 0 | 0 |
| BTO2_18 | Bocas del Toro | Almirante | 9.29607 | -82.40906 | 2017 | 1 | 0 |
| BTO2_19 | Bocas del Toro | Almirante | 9.29371 | -82.40602 | 2017 | 1 | 0 |
| BTO2_2 | Bocas del Toro | Almirante | 9.29085 | -82.39806 | 2017 | 0 | 0 |
| BTO2_20 | Bocas del Toro | Almirante | 9.29314 | -82.40593 | 2017 | 1 | 0 |
| BTO2_3 | Bocas del Toro | Almirante | 9.29038 | -82.39964 | 2017 | 0 | 0 |
| BTO2_4 | Bocas del Toro | Almirante | 9.28994 | -82.40094 | 2017 | 1 | 0 |
| BTO2_5 | Bocas del Toro | Almirante | 9.29066 | -82.40112 | 2017 | 1 | 0 |
| BTO2_6 | Bocas del Toro | Almirante | 9.29124 | -82.40188 | 2017 | 0 | 0 |
| BTO2_7 | Bocas del Toro | Almirante | 9.29066 | -82.39038 | 2017 | 1 | 0 |
| BTO2_8 | Bocas del Toro | Almirante | 9.29459 | -82.38883 | 2017 | 0 | 0 |
| BTO2_9 | Bocas del Toro | Almirante | 9.29355 | -82.39104 | 2017 | 0 | 0 |
| BTO4_1 | Bocas del Toro | Changuinola | 9.47043 | -82.51347 | 2017 | 1 | 0 |
| BTO4_10 | Bocas del Toro | Changuinola | 9.46501 | -82.51825 | 2017 | 1 | 0 |
| BTO4_11 | Bocas del Toro | Changuinola | 9.46476 | -82.51961 | 2017 | 0 | 0 |
| BTO4_12 | Bocas del Toro | Changuinola | 9.45126 | -82.52164 | 2017 | 0 | 0 |
| BTO4_13 | Bocas del Toro | Changuinola | 9.45074 | -82.52531 | 2017 | 1 | 0 |
| BTO4_14 | Bocas del Toro | Changuinola | 9.45021 | -82.52376 | 2017 | 1 | 0 |
| BTO4_15 | Bocas del Toro | Changuinola | 9.45074 | -82.52254 | 2017 | 1 | 0 |
| BTO4_16 | Bocas del Toro | Changuinola | 9.44972 | -82.51992 | 2017 | 1 | 0 |
| BTO4_17 | Bocas del Toro | Changuinola | 9.45145 | -82.52000 | 2017 | 1 | 0 |
| BTO4_18 | Bocas del Toro | Changuinola | 9.45466 | -82.51648 | 2017 | 1 | 0 |
| BTO4_19 | Bocas del Toro | Changuinola | 9.45668 | -82.51567 | 2017 | 0 | 0 |
| BTO4_2 | Bocas del Toro | Changuinola | 9.47201 | -82.51465 | 2017 | 0 | 0 |
| BTO4_20 | Bocas del Toro | Changuinola | 9.45624 | -82.51524 | 2017 | 0 | 0 |
| BTO4_3 | Bocas del Toro | Changuinola | 9.47761 | -82.51069 | 2017 | 1 | 0 |
| BTO4_4 | Bocas del Toro | Changuinola | 9.47476 | -82.50778 | 2017 | 1 | 0 |
| BTO4_5 | Bocas del Toro | Changuinola | 9.47811 | -82.50884 | 2017 | 0 | 0 |
| BTO4_6 | Bocas del Toro | Changuinola | 9.47501 | -82.50895 | 2017 | 1 | 0 |
| BTO4_7 | Bocas del Toro | Changuinola | 9.47289 | -82.51032 | 2017 | 1 | 0 |
| BTO4_8 | Bocas del Toro | Changuinola | 9.47368 | -82.51317 | 2017 | 1 | 0 |
| BTO4_9 | Bocas del Toro | Changuinola | 9.46629 | -82.51824 | 2017 | 1 | 0 |
| CHI5_1 | Chiriquí | Gualaca | 8.53681 | -82.29827 | 2017 | 0 | 0 |
| CHI5_10 | Chiriquí | Gualaca | 8.52329 | -82.29837 | 2017 | 0 | 1 |
| CHI5_11 | Chiriquí | Gualaca | 8.52355 | -82.30174 | 2017 | 0 | 0 |
| CHI5_12 | Chiriquí | Gualaca | 8.52887 | -82.30254 | 2017 | 0 | 1 |
| CHI5_13 | Chiriquí | Gualaca | 8.52966 | -82.30426 | 2017 | 0 | 0 |
| CHI5_14 | Chiriquí | Gualaca | 8.53144 | -82.30273 | 2017 | 0 | 0 |
| CHI5_15 | Chiriquí | Gualaca | 8.53364 | -82.30144 | 2017 | 0 | 1 |
| CHI5_16 | Chiriquí | Gualaca | 8.53152 | -82.30043 | 2017 | 0 | 1 |

| | | | | | | | |
|---|---|---|---|---|---|---|---|
| CHI5_17 | Chiriquí | Gualaca | 8.53610 | -82.30018 | 2017 | 0 | 1 |
| CHI5_18 | Chiriquí | Gualaca | 8.53702 | -82.30411 | 2017 | 0 | 0 |
| CHI5_19 | Chiriquí | Gualaca | 8.53555 | -82.30244 | 2017 | 0 | 1 |
| CHI5_2 | Chiriquí | Gualaca | 8.53404 | -82.29825 | 2017 | 0 | 1 |
| CHI5_20 | Chiriquí | Gualaca | 8.53237 | -82.29972 | 2017 | 0 | 1 |
| CHI5_3 | Chiriquí | Gualaca | 8.53318 | -82.29775 | 2017 | 0 | 0 |
| CHI5_4 | Chiriquí | Gualaca | 8.53165 | -82.29621 | 2017 | 0 | 1 |
| CHI5_5 | Chiriquí | Gualaca | 8.52856 | -82.29667 | 2017 | 0 | 0 |
| CHI5_6 | Chiriquí | Gualaca | 8.52826 | -82.29741 | 2017 | 0 | 1 |
| CHI5_7 | Chiriquí | Gualaca | 8.52661 | -82.29647 | 2017 | 0 | 1 |
| CHI5_8 | Chiriquí | Gualaca | 8.52666 | -82.29798 | 2017 | 0 | 0 |
| CHI5_9 | Chiriquí | Gualaca | 8.52568 | -82.29861 | 2017 | 0 | 0 |
| CHI7_1 | Chiriquí | David | 8.41576 | -82.42724 | 2017 | 0 | 0 |
| CHI7_10 | Chiriquí | David | 8.41934 | -82.42655 | 2017 | 0 | 1 |
| CHI7_11 | Chiriquí | David | 8.41840 | -82.42369 | 2017 | 0 | 0 |
| CHI7_12 | Chiriquí | David | 8.41726 | -82.42583 | 2017 | 0 | 0 |
| CHI7_13 | Chiriquí | David | 8.42086 | -82.42602 | 2017 | 0 | 1 |
| CHI7_14 | Chiriquí | David | 8.42331 | -82.42231 | 2017 | 0 | 0 |
| CHI7_15 | Chiriquí | David | 8.42155 | -82.42305 | 2017 | 0 | 1 |
| CHI7_16 | Chiriquí | David | 8.42465 | -82.42465 | 2017 | 0 | 1 |
| CHI7_17 | Chiriquí | David | 8.42097 | -82.42846 | 2017 | 1 | 1 |
| CHI7_18 | Chiriquí | David | 8.42209 | -82.43100 | 2017 | 1 | 0 |
| CHI7_19 | Chiriquí | David | 8.42056 | -82.43245 | 2017 | 0 | 1 |
| CHI7_2 | Chiriquí | David | 8.41235 | -82.42681 | 2017 | 0 | 0 |
| CHI7_20 | Chiriquí | David | 8.41583 | -82.43278 | 2017 | 1 | 1 |
| CHI7_3 | Chiriquí | David | 8.41327 | -82.42594 | 2017 | 0 | 0 |
| CHI7_4 | Chiriquí | David | 8.41238 | -82.42454 | 2017 | 0 | 0 |
| CHI7_5 | Chiriquí | David | 8.41088 | -82.42372 | 2017 | 0 | 0 |
| CHI7_6 | Chiriquí | David | 8.41309 | -82.42322 | 2017 | 0 | 1 |
| CHI7_7 | Chiriquí | David | 8.41412 | -82.42660 | 2017 | 0 | 0 |
| CHI7_8 | Chiriquí | David | 8.41863 | -82.42811 | 2017 | 0 | 0 |
| CHI7_9 | Chiriquí | David | 8.41855 | -82.42970 | 2017 | 0 | 1 |
| COL3_1 | Colón | Portobelo | 9.55349 | -79.64700 | 2017 | 0 | 0 |
| COL3_10 | Colón | Portobelo | 9.55450 | -79.65353 | 2017 | 1 | 0 |
| COL3_11 | Colón | Portobelo | 9.55471 | -79.65381 | 2017 | 0 | 0 |
| COL3_12 | Colón | Portobelo | 9.55319 | -79.65465 | 2017 | 0 | 0 |
| COL3_13 | Colón | Portobelo | 9.55267 | -79.65509 | 2017 | 0 | 0 |
| COL3_14 | Colón | Portobelo | 9.55178 | -79.65514 | 2017 | 0 | 0 |
| COL3_15 | Colón | Portobelo | 9.55309 | -79.65549 | 2017 | 0 | 0 |
| COL3_16 | Colón | Portobelo | 9.55323 | -79.65682 | 2017 | 0 | 0 |
| COL3_17 | Colón | Portobelo | 9.54853 | -79.67016 | 2017 | 0 | 0 |
| COL3_18 | Colón | Portobelo | 9.54586 | -79.67137 | 2017 | 0 | 0 |
| COL3_19 | Colón | Portobelo | 9.54458 | -79.67311 | 2017 | 0 | 0 |

| | | | | | | | |
|---|---|---|---|---|---|---|---|
| COL3_2 | Colón | Portobelo | 9.55315 | -79.65012 | 2017 | 0 | 0 |
| COL3_3 | Colón | Portobelo | 9.55401 | -79.65071 | 2017 | 0 | 0 |
| COL3_4 | Colón | Portobelo | 9.55402 | -79.65098 | 2017 | 0 | 0 |
| COL3_5 | Colón | Portobelo | 9.55448 | -79.65130 | 2017 | 0 | 0 |
| COL3_6 | Colón | Portobelo | 9.55426 | -79.65158 | 2017 | 0 | 0 |
| COL3_7 | Colón | Portobelo | 9.55599 | -79.65195 | 2017 | 0 | 0 |
| COL3_8 | Colón | Portobelo | 9.55498 | -79.65178 | 2017 | 0 | 0 |
| COL3_9 | Colón | Portobelo | 9.55527 | -79.65250 | 2017 | 0 | 0 |
| COL5_1 | Colón | Sabanitas | 9.35212 | -79.80632 | 2017 | 1 | 1 |
| COL5_2 | Colón | Sabanitas | 9.35377 | -79.80870 | 2017 | 1 | 1 |
| COL5_3 | Colón | Sabanitas | 9.35481 | -79.80865 | 2017 | 1 | 0 |
| COL5_4 | Colón | Sabanitas | 9.35255 | -79.80522 | 2017 | 0 | 1 |
| COL5_5 | Colón | Sabanitas | 9.35362 | -79.80478 | 2017 | 1 | 0 |
| COL5_6 | Colón | Sabanitas | 9.35433 | -79.80586 | 2017 | 1 | 1 |
| COL5_7 | Colón | Sabanitas | 9.35584 | -79.80803 | 2017 | 0 | 0 |
| COL5_8 | Colón | Sabanitas | 9.35569 | -79.80716 | 2017 | 0 | 0 |
| COL5_9 | Colón | Sabanitas | 9.35039 | -79.80681 | 2017 | 0 | 1 |
| COL5_10 | Colón | Sabanitas | 9.35009 | -79.80798 | 2017 | 0 | 0 |
| COL5_11 | Colón | Sabanitas | 9.35068 | -79.79895 | 2017 | 1 | 1 |
| COL5_12 | Colón | Sabanitas | 9.35005 | -79.79758 | 2017 | 0 | 0 |
| COL5_13 | Colón | Sabanitas | 9.35089 | -79.79673 | 2017 | 0 | 0 |
| COL5_14 | Colón | Sabanitas | 9.34978 | -79.79811 | 2017 | 1 | 0 |
| COL5_15 | Colón | Sabanitas | 9.34790 | -79.79729 | 2017 | 0 | 0 |
| COL5_16 | Colón | Sabanitas | 9.34612 | -79.79597 | 2017 | 0 | 0 |
| COL5_17 | Colón | Sabanitas | 9.35078 | -79.80029 | 2017 | 0 | 0 |
| COL5_18 | Colón | Sabanitas | 9.35157 | -79.80105 | 2017 | 1 | 0 |
| COL5_19 | Colón | Sabanitas | 9.35442 | -79.80220 | 2017 | 0 | 0 |
| COL5_20 | Colón | Sabanitas | 9.35368 | -79.80125 | 2017 | 0 | 0 |
| COL5_21 | Colón | Sabanitas | 9.35208 | -79.80168 | 2017 | 0 | 0 |
| COL5_22 | Colón | Sabanitas | 9.35402 | -79.80179 | 2017 | 1 | 1 |
| COL5_23 | Colón | Sabanitas | 9.35400 | -79.80115 | 2017 | 1 | 1 |
| COL5_24 | Colón | Sabanitas | 9.35386 | -79.80271 | 2017 | 0 | 0 |
| COL5_25 | Colón | Sabanitas | 9.35219 | -79.80634 | 2017 | 1 | 0 |
| COL5_26 | Colón | Sabanitas | 9.35331 | -79.80805 | 2017 | 1 | 1 |
| COL5_27 | Colón | Sabanitas | 9.35379 | -79.80877 | 2017 | 0 | 0 |
| COL5_28 | Colón | Sabanitas | 9.35450 | -79.80994 | 2017 | 0 | 0 |
| COL5_29 | Colón | Sabanitas | 9.35295 | -79.80581 | 2017 | 1 | 1 |
| COL5_30 | Colón | Sabanitas | 9.35286 | -79.80456 | 2017 | 0 | 0 |
| COL5_31 | Colón | Sabanitas | 9.35363 | -79.80479 | 2017 | 1 | 0 |
| COL5_32 | Colón | Sabanitas | 9.35653 | -79.80912 | 2017 | 0 | 0 |
| COL5_33 | Colón | Sabanitas | 9.35569 | -79.80718 | 2017 | 0 | 0 |
| COL5_34 | Colón | Sabanitas | 9.35010 | -79.80703 | 2017 | 0 | 0 |
| COL5_35 | Colón | Sabanitas | 9.35178 | -79.80686 | 2017 | 1 | 0 |

| | | | | | | | |
|---|---|---|---|---|---|---|---|
| COL5_36 | Colón | Sabanitas | 9.35053 | -79.80329 | 2017 | 0 | 0 |
| COL6_1 | Colón | Gamboa | 9.11890 | -79.69860 | 2017 | 0 | 1 |
| COL6_2 | Colón | Gamboa | 9.11780 | -79.69700 | 2017 | 0 | 0 |
| COL6_3 | Colón | Gamboa | 9.11800 | -79.69633 | 2017 | 0 | 0 |
| COL6_4 | Colón | Gamboa | 9.11680 | -79.69800 | 2017 | 0 | 1 |
| COL6_5 | Colón | Gamboa | 9.11680 | -79.69800 | 2017 | 0 | 0 |
| COL6_6 | Colón | Gamboa | 9.11680 | -79.69840 | 2017 | 0 | 0 |
| COL6_7 | Colón | Gamboa | 9.11670 | -79.69890 | 2017 | 0 | 0 |
| COL6_8 | Colón | Gamboa | 9.11630 | -79.69800 | 2017 | 0 | 1 |
| COL6_9 | Colón | Gamboa | 9.11600 | -79.69700 | 2017 | 0 | 0 |
| COL6_10 | Colón | Gamboa | 9.11810 | -79.69800 | 2017 | 0 | 0 |
| COL6_11 | Colón | Gamboa | 9.11960 | -79.70110 | 2017 | 0 | 0 |
| COL6_12 | Colón | Gamboa | 9.11960 | -79.70110 | 2017 | 0 | 0 |
| COL6_13 | Colón | Gamboa | 9.11890 | -79.69860 | 2017 | 0 | 0 |
| COL6_14 | Colón | Gamboa | 9.11780 | -79.69700 | 2017 | 0 | 1 |
| COL6_15 | Colón | Gamboa | 9.11800 | -79.69633 | 2017 | 0 | 0 |
| COL6_16 | Colón | Gamboa | 9.11680 | -79.69800 | 2017 | 0 | 0 |
| COL6_17 | Colón | Gamboa | 9.11680 | -79.69800 | 2017 | 0 | 1 |
| COL6_18 | Colón | Gamboa | 9.11680 | -79.69840 | 2017 | 0 | 0 |
| COL6_19 | Colón | Gamboa | 9.11670 | -79.69890 | 2017 | 0 | 0 |
| COL6_20 | Colón | Gamboa | 9.11630 | -79.69800 | 2017 | 0 | 1 |
| COL6_21 | Colón | Gamboa | 9.11600 | -79.69700 | 2017 | 0 | 0 |
| COL6_22 | Colón | Gamboa | 9.11810 | -79.69800 | 2017 | 0 | 1 |
| COL6_23 | Colón | Gamboa | 9.11960 | -79.70110 | 2017 | 0 | 0 |
| COL6_24 | Colón | Gamboa | 9.11960 | -79.70110 | 2017 | 0 | 0 |
| DAR2_1 | Darién | Metetí | 8.50225 | -77.98127 | 2017 | 0 | 0 |
| DAR2_10 | Darién | Metetí | 8.49137 | -77.98262 | 2017 | 1 | 0 |
| DAR2_11 | Darién | Metetí | 8.49647 | -77.98402 | 2017 | 0 | 0 |
| DAR2_12 | Darién | Metetí | 8.49889 | -77.98247 | 2017 | 1 | 1 |
| DAR2_13 | Darién | Metetí | 8.50163 | -77.98375 | 2017 | 0 | 1 |
| DAR2_14 | Darién | Metetí | 8.50436 | -77.97404 | 2017 | 0 | 1 |
| DAR2_15 | Darién | Metetí | 8.50442 | -77.97133 | 2017 | 0 | 1 |
| DAR2_16 | Darién | Metetí | 8.50450 | -77.96847 | 2017 | 0 | 0 |
| DAR2_17 | Darién | Metetí | 8.50689 | -77.97587 | 2017 | 0 | 1 |
| DAR2_18 | Darién | Metetí | 8.51266 | -77.97822 | 2017 | 0 | 1 |
| DAR2_2 | Darién | Metetí | 8.50265 | -77.97826 | 2017 | 0 | 1 |
| DAR2_3 | Darién | Metetí | 8.49974 | -77.97696 | 2017 | 0 | 1 |
| DAR2_4 | Darién | Metetí | 8.49929 | -77.97421 | 2017 | 0 | 0 |
| DAR2_5 | Darién | Metetí | 8.49756 | -77.97101 | 2017 | 0 | 0 |
| DAR2_6 | Darién | Metetí | 8.49410 | -77.96969 | 2017 | 1 | 0 |
| DAR2_7 | Darién | Metetí | 8.49760 | -77.97995 | 2017 | 0 | 0 |
| DAR2_8 | Darién | Metetí | 8.49503 | -77.98141 | 2017 | 0 | 0 |
| DAR2_9 | Darién | Metetí | 8.49257 | -77.98041 | 2017 | 1 | 0 |

| | | | | | | | |
|---|---|---|---|---|---|---|---|
| LSA1_1 | Los Santos | La Villa | 7.93664 | -80.41697 | 2017 | 0 | 0 |
| LSA1_10 | Los Santos | La Villa | 7.93861 | -80.41169 | 2017 | 1 | 1 |
| LSA1_11 | Los Santos | La Villa | 7.93789 | -80.41103 | 2017 | 1 | 0 |
| LSA1_12 | Los Santos | La Villa | 7.93572 | -80.41133 | 2017 | 0 | 0 |
| LSA1_13 | Los Santos | La Villa | 7.93058 | -80.40700 | 2017 | 1 | 1 |
| LSA1_14 | Los Santos | La Villa | 7.93006 | -80.40772 | 2017 | 0 | 0 |
| LSA1_15 | Los Santos | La Villa | 7.92606 | -80.40503 | 2017 | 0 | 0 |
| LSA1_16 | Los Santos | La Villa | 7.93125 | -80.41225 | 2017 | 1 | 1 |
| LSA1_17 | Los Santos | La Villa | 7.93036 | -80.41294 | 2017 | 1 | 1 |
| LSA1_18 | Los Santos | La Villa | 7.93181 | -80.41603 | 2017 | 1 | 1 |
| LSA1_19 | Los Santos | La Villa | 7.93239 | -80.41794 | 2017 | 1 | 1 |
| LSA1_2 | Los Santos | La Villa | 7.93772 | -80.41756 | 2017 | 0 | 0 |
| LSA1_20 | Los Santos | La Villa | 7.93192 | -80.41753 | 2017 | 1 | 1 |
| LSA1_3 | Los Santos | La Villa | 7.93933 | -80.41725 | 2017 | 1 | 0 |
| LSA1_4 | Los Santos | La Villa | 7.94050 | -80.41681 | 2017 | 1 | 0 |
| LSA1_5 | Los Santos | La Villa | 7.94217 | -80.41767 | 2017 | 0 | 1 |
| LSA1_6 | Los Santos | La Villa | 7.94061 | -80.41517 | 2017 | 1 | 1 |
| LSA1_7 | Los Santos | La Villa | 7.93883 | -80.41550 | 2017 | 1 | 1 |
| LSA1_8 | Los Santos | La Villa | 7.93964 | -80.41306 | 2017 | 1 | 1 |
| LSA1_9 | Los Santos | La Villa | 7.94044 | -80.41064 | 2017 | 1 | 1 |
| LSA3_1 | Los Santos | Tonosí | 7.40692 | -80.44378 | 2017 | 0 | 1 |
| LSA3_10 | Los Santos | Tonosí | 7.40769 | -80.43667 | 2017 | 0 | 1 |
| LSA3_11 | Los Santos | Tonosí | 7.40944 | -80.43986 | 2017 | 0 | 1 |
| LSA3_12 | Los Santos | Tonosí | 7.41044 | -80.44322 | 2017 | 0 | 1 |
| LSA3_13 | Los Santos | Tonosí | 7.41100 | -80.44256 | 2017 | 0 | 1 |
| LSA3_14 | Los Santos | Tonosí | 7.41119 | -80.44567 | 2017 | 0 | 1 |
| LSA3_15 | Los Santos | Tonosí | 7.41083 | -80.43533 | 2017 | 0 | 1 |
| LSA3_16 | Los Santos | Tonosí | 7.40942 | -80.43528 | 2017 | 0 | 0 |
| LSA3_17 | Los Santos | Tonosí | 7.40850 | -80.43592 | 2017 | 0 | 1 |
| LSA3_18 | Los Santos | Tonosí | 7.40956 | -80.43667 | 2017 | 0 | 1 |
| LSA3_19 | Los Santos | Tonosí | 7.41158 | -80.43650 | 2017 | 0 | 0 |
| LSA3_2 | Los Santos | Tonosí | 7.40506 | -80.44314 | 2017 | 0 | 1 |
| LSA3_20 | Los Santos | Tonosí | 7.40692 | -80.44203 | 2017 | 0 | 1 |
| LSA3_3 | Los Santos | Tonosí | 7.40503 | -80.44169 | 2017 | 0 | 1 |
| LSA3_4 | Los Santos | Tonosí | 7.40647 | -80.44117 | 2017 | 0 | 0 |
| LSA3_5 | Los Santos | Tonosí | 7.40478 | -80.43992 | 2017 | 0 | 1 |
| LSA3_6 | Los Santos | Tonosí | 7.40569 | -80.43917 | 2017 | 0 | 1 |
| LSA3_7 | Los Santos | Tonosí | 7.40506 | -80.43739 | 2017 | 0 | 1 |
| LSA3_8 | Los Santos | Tonosí | 7.40542 | -80.43592 | 2017 | 0 | 1 |
| LSA3_9 | Los Santos | Tonosí | 7.40689 | -80.43747 | 2017 | 0 | 1 |
| LSA4_1 | Los Santos | El Cacao | 7.44064 | -80.41206 | 2017 | 0 | 1 |
| LSA4_10 | Los Santos | El Cacao | 7.44664 | -80.40453 | 2017 | 0 | 0 |
| LSA4_11 | Los Santos | El Cacao | 7.44797 | -80.40333 | 2017 | 0 | 1 |

| | | | | | | | |
|---|---|---|---|---|---|---|---|
| LSA4_12 | Los Santos | El Cacao | 7.44697 | -80.40306 | 2017 | 0 | 1 |
| LSA4_13 | Los Santos | El Cacao | 7.44939 | -80.40300 | 2017 | 0 | 1 |
| LSA4_14 | Los Santos | El Cacao | 7.45036 | -80.40375 | 2017 | 0 | 1 |
| LSA4_15 | Los Santos | El Cacao | 7.45189 | -80.40306 | 2017 | 0 | 1 |
| LSA4_16 | Los Santos | El Cacao | 7.45200 | -80.40036 | 2017 | 0 | 1 |
| LSA4_17 | Los Santos | El Cacao | 7.45081 | -80.40044 | 2017 | 0 | 0 |
| LSA4_18 | Los Santos | El Cacao | 7.45128 | -80.40414 | 2017 | 0 | 1 |
| LSA4_2 | Los Santos | El Cacao | 7.44186 | -80.41122 | 2017 | 0 | 1 |
| LSA4_3 | Los Santos | El Cacao | 7.44231 | -80.41097 | 2017 | 0 | 1 |
| LSA4_4 | Los Santos | El Cacao | 7.44508 | -80.40936 | 2017 | 0 | 0 |
| LSA4_5 | Los Santos | El Cacao | 7.44561 | -80.40908 | 2017 | 0 | 1 |
| LSA4_6 | Los Santos | El Cacao | 7.44856 | -80.40617 | 2017 | 0 | 0 |
| LSA4_7 | Los Santos | El Cacao | 7.44878 | -80.40456 | 2017 | 0 | 1 |
| LSA4_8 | Los Santos | El Cacao | 7.44844 | -80.40450 | 2017 | 0 | 1 |
| LSA4_9 | Los Santos | El Cacao | 7.44786 | -80.40433 | 2017 | 0 | 1 |
| LSA5_1 | Los Santos | Pedasi | 7.53342 | -80.02778 | 2017 | 1 | 1 |
| LSA5_10 | Los Santos | Pedasi | 7.52672 | -80.01819 | 2017 | 1 | 1 |
| LSA5_11 | Los Santos | Pedasi | 7.52750 | -80.01703 | 2017 | 1 | 1 |
| LSA5_12 | Los Santos | Pedasi | 7.52681 | -80.01828 | 2017 | 1 | 1 |
| LSA5_13 | Los Santos | Pedasi | 7.52800 | -80.01706 | 2017 | 1 | 1 |
| LSA5_14 | Los Santos | Pedasi | 7.52856 | -80.01869 | 2017 | 1 | 1 |
| LSA5_15 | Los Santos | Pedasi | 7.53106 | -80.01731 | 2017 | 0 | 1 |
| LSA5_16 | Los Santos | Pedasi | 7.53150 | -80.01922 | 2017 | 0 | 1 |
| LSA5_17 | Los Santos | Pedasi | 7.53372 | -80.01706 | 2017 | 1 | 0 |
| LSA5_18 | Los Santos | Pedasi | 7.53278 | -80.01775 | 2017 | 1 | 0 |
| LSA5_19 | Los Santos | Pedasi | 7.53161 | -80.01769 | 2017 | 1 | 1 |
| LSA5_2 | Los Santos | Pedasi | 7.53281 | -80.02972 | 2017 | 1 | 1 |
| LSA5_20 | Los Santos | Pedasi | 7.53014 | -80.01936 | 2017 | 1 | 1 |
| LSA5_3 | Los Santos | Pedasi | 7.53083 | -80.01831 | 2017 | 0 | 1 |
| LSA5_4 | Los Santos | Pedasi | 7.52972 | -80.01836 | 2017 | 1 | 1 |
| LSA5_5 | Los Santos | Pedasi | 7.52972 | -80.01683 | 2017 | 0 | 0 |
| LSA5_6 | Los Santos | Pedasi | 7.52889 | -80.01672 | 2017 | 1 | 0 |
| LSA5_7 | Los Santos | Pedasi | 7.53111 | -80.01917 | 2017 | 1 | 1 |
| LSA5_8 | Los Santos | Pedasi | 7.53056 | -80.01747 | 2017 | 1 | 1 |
| LSA5_9 | Los Santos | Pedasi | 7.52514 | -80.01919 | 2017 | 1 | 0 |
| PAN1_1 | Panamá Oeste | Princesa Mía | 8.96544 | -79.70419 | 2017 | 0 | 1 |
| PAN1_10 | Panamá Oeste | Princesa Mía | 8.96519 | -79.70088 | 2017 | 0 | 1 |
| PAN1_11 | Panamá Oeste | Princesa Mía | 8.96575 | -79.70074 | 2017 | 1 | 0 |
| PAN1_12 | Panamá Oeste | Princesa Mía | 8.96613 | -79.70138 | 2017 | 1 | 0 |
| PAN1_2 | Panamá Oeste | Princesa Mía | 8.96468 | -79.70362 | 2017 | 0 | 0 |
| PAN1_3 | Panamá Oeste | Princesa Mía | 8.96561 | -79.70400 | 2017 | 0 | 0 |
| PAN1_4 | Panamá Oeste | Princesa Mía | 8.96483 | -79.70318 | 2017 | 0 | 0 |
| PAN1_5 | Panamá Oeste | Princesa Mía | 8.96486 | -79.70271 | 2017 | 0 | 1 |

| | | | | | | | |
|---|---|---|---|---|---|---|---|
| PAN1_6 | Panamá Oeste | Princesa Mía | 8.96446 | -79.70205 | 2017 | 0 | 1 |
| PAN1_7 | Panamá Oeste | Princesa Mía | 8.96554 | -79.70212 | 2017 | 0 | 1 |
| PAN1_8 | Panamá Oeste | Princesa Mía | 8.96576 | -79.70152 | 2017 | 1 | 1 |
| PAN1_9 | Panamá Oeste | Princesa Mía | 8.96495 | -79.70167 | 2017 | 1 | 0 |
| PAN2_1 | Panamá Oeste | Lluvia de Oro | 8.96010 | -79.70202 | 2017 | 0 | 0 |
| PAN2_10 | Panamá Oeste | Lluvia de Oro | 8.96053 | -79.69965 | 2017 | 1 | 1 |
| PAN2_11 | Panamá Oeste | Lluvia de Oro | 8.96050 | -79.69923 | 2017 | 1 | 1 |
| PAN2_12 | Panamá Oeste | Lluvia de Oro | 8.95975 | -79.69833 | 2017 | 1 | 0 |
| PAN2_2 | Panamá Oeste | Lluvia de Oro | 8.96035 | -79.70180 | 2017 | 1 | 0 |
| PAN2_3 | Panamá Oeste | Lluvia de Oro | 8.95872 | -79.70157 | 2017 | 0 | 0 |
| PAN2_4 | Panamá Oeste | Lluvia de Oro | 8.95941 | -79.70195 | 2017 | 1 | 1 |
| PAN2_5 | Panamá Oeste | Lluvia de Oro | 8.96081 | -79.70141 | 2017 | 0 | 1 |
| PAN2_6 | Panamá Oeste | Lluvia de Oro | 8.95921 | -79.70106 | 2017 | 1 | 1 |
| PAN2_7 | Panamá Oeste | Lluvia de Oro | 8.96045 | -79.70089 | 2017 | 0 | 0 |
| PAN2_8 | Panamá Oeste | Lluvia de Oro | 8.95916 | -79.70050 | 2017 | 1 | 1 |
| PAN2_9 | Panamá Oeste | Lluvia de Oro | 8.96058 | -79.70038 | 2017 | 1 | 0 |
| PAN3_1 | Panamá Oeste | Nuevo Chorrillo | 8.95236 | -79.69890 | 2017 | 1 | 1 |
| PAN3_10 | Panamá Oeste | Nuevo Chorrillo | 8.95527 | -79.69450 | 2017 | 1 | 1 |
| PAN3_11 | Panamá Oeste | Nuevo Chorrillo | 8.95804 | -79.69421 | 2017 | 0 | 0 |
| PAN3_12 | Panamá Oeste | Nuevo Chorrillo | 8.95712 | -79.69354 | 2017 | 0 | 0 |
| PAN3_2 | Panamá Oeste | Nuevo Chorrillo | 8.95417 | -79.69722 | 2017 | 0 | 1 |
| PAN3_3 | Panamá Oeste | Nuevo Chorrillo | 8.95500 | -79.69629 | 2017 | 1 | 1 |
| PAN3_4 | Panamá Oeste | Nuevo Chorrillo | 8.95582 | -79.69528 | 2017 | 0 | 0 |
| PAN3_5 | Panamá Oeste | Nuevo Chorrillo | 8.95355 | -79.69507 | 2017 | 1 | 0 |
| PAN3_6 | Panamá Oeste | Nuevo Chorrillo | 8.95287 | -79.69628 | 2017 | 1 | 1 |
| PAN3_7 | Panamá Oeste | Nuevo Chorrillo | 8.95177 | -79.69705 | 2017 | 1 | 1 |
| PAN3_8 | Panamá Oeste | Nuevo Chorrillo | 8.95154 | -79.69840 | 2017 | 1 | 1 |
| PAN3_9 | Panamá Oeste | Nuevo Chorrillo | 8.95281 | -79.69784 | 2017 | 1 | 0 |
| PAN4_1 | Panamá | Chilibre | 9.16880 | -79.61697 | 2017 | 0 | 1 |
| PAN4_2 | Panamá | Chilibre | 9.16821 | -79.61460 | 2017 | 0 | 0 |
| PAN4_3 | Panamá | Chilibre | 9.16771 | -79.61249 | 2017 | 0 | 0 |
| PAN4_4 | Panamá | Chilibre | 9.16952 | -79.61151 | 2017 | 0 | 0 |
| PAN4_5 | Panamá | Chilibre | 9.17524 | -79.60968 | 2017 | 0 | 0 |
| PAN4_6 | Panamá | Chilibre | 9.17808 | -79.61234 | 2017 | 0 | 0 |
| PAN4_7 | Panamá | Chilibre | 9.17697 | -79.61236 | 2017 | 0 | 0 |
| PAN4_8 | Panamá | Chilibre | 9.17449 | -79.61092 | 2017 | 0 | 1 |
| PAN4_9 | Panamá | Chilibre | 9.17341 | -79.61113 | 2017 | 0 | 1 |
| PAN5_1 | Panamá | Arraiján | 8.94794 | -79.64919 | 2017 | 0 | 1 |
| PAN5_10 | Panamá | Arraiján | 8.94217 | -79.64014 | 2017 | 0 | 0 |
| PAN5_2 | Panamá | Arraiján | 8.94261 | -79.64602 | 2017 | 1 | 1 |
| PAN5_3 | Panamá | Arraiján | 8.94166 | -79.64613 | 2017 | 1 | 1 |
| PAN5_4 | Panamá | Arraiján | 8.94001 | -79.64573 | 2017 | 1 | 1 |
| PAN5_5 | Panamá | Arraiján | 8.94266 | -79.64424 | 2017 | 0 | 1 |

| | | | | | | | |
|---|---|---|---|---|---|---|---|
| PAN5_6 | Panamá | Arraiján | 8.94190 | -79.64379 | 2017 | 1 | 0 |
| PAN5_7 | Panamá | Arraiján | 8.94084 | -79.64120 | 2017 | 0 | 0 |
| PAN5_8 | Panamá | Arraiján | 8.94203 | -79.64092 | 2017 | 1 | 1 |
| PAN5_9 | Panamá | Arraiján | 8.94383 | -79.64043 | 2017 | 0 | 0 |
| PAN6_1 | Panamá | Chepo | 9.16706 | -79.09197 | 2017 | 1 | 1 |
| PAN6_10 | Panamá | Chepo | 9.16089 | -79.09720 | 2017 | 0 | 0 |
| PAN6_11 | Panamá | Chepo | 9.16073 | -79.09930 | 2017 | 0 | 1 |
| PAN6_12 | Panamá | Chepo | 9.16030 | -79.10192 | 2017 | 0 | 1 |
| PAN6_13 | Panamá | Chepo | 9.16200 | -79.09916 | 2017 | 0 | 0 |
| PAN6_14 | Panamá | Chepo | 9.16305 | -79.09672 | 2017 | 0 | 0 |
| PAN6_15 | Panamá | Chepo | 9.16592 | -79.09475 | 2017 | 0 | 0 |
| PAN6_16 | Panamá | Chepo | 9.16647 | -79.09570 | 2017 | 1 | 0 |
| PAN6_17 | Panamá | Chepo | 9.16527 | -79.09679 | 2017 | 1 | 0 |
| PAN6_18 | Panamá | Chepo | 9.16277 | -79.09877 | 2017 | 1 | 1 |
| PAN6_19 | Panamá | Chepo | 9.16450 | -79.09892 | 2017 | 0 | 0 |
| PAN6_2 | Panamá | Chepo | 9.16573 | -79.09284 | 2017 | 1 | 1 |
| PAN6_20 | Panamá | Chepo | 9.16643 | -79.09803 | 2017 | 0 | 1 |
| PAN6_3 | Panamá | Chepo | 9.16415 | -79.09185 | 2017 | 1 | 0 |
| PAN6_4 | Panamá | Chepo | 9.16155 | -79.09193 | 2017 | 0 | 0 |
| PAN6_5 | Panamá | Chepo | 9.16296 | -79.09338 | 2017 | 0 | 0 |
| PAN6_6 | Panamá | Chepo | 9.16219 | -79.09454 | 2017 | 0 | 0 |
| PAN6_7 | Panamá | Chepo | 9.16042 | -79.09392 | 2017 | 0 | 0 |
| PAN6_8 | Panamá | Chepo | 9.15905 | -79.09462 | 2017 | 1 | 1 |
| PAN6_9 | Panamá | Chepo | 9.15956 | -79.09610 | 2017 | 0 | 0 |
| LSA3_1 | Los Santos | Tonosí | 7.40694 | -80.44361 | 2018 | 0 | 1 |
| LSA3_2 | Los Santos | Tonosí | 7.40528 | -80.44306 | 2018 | 0 | 1 |
| LSA3_3 | Los Santos | Tonosí | 7.38169 | -80.44306 | 2018 | 0 | 1 |
| LSA3_4 | Los Santos | Tonosí | 7.40444 | -80.43833 | 2018 | 0 | 0 |
| LSA3_5 | Los Santos | Tonosí | 7.40556 | -80.43694 | 2018 | 0 | 1 |
| LSA3_6 | Los Santos | Tonosí | 7.40528 | -80.43972 | 2018 | 0 | 1 |
| LSA3_7 | Los Santos | Tonosí | 7.42500 | -80.43333 | 2018 | 0 | 1 |
| LSA3_8 | Los Santos | Tonosí | 7.41611 | -80.40722 | 2018 | 0 | 1 |
| LSA3_9 | Los Santos | Tonosí | 7.42583 | -80.37694 | 2018 | 0 | 0 |
| LSA3_10 | Los Santos | Tonosí | 7.42167 | -80.39333 | 2018 | 0 | 1 |
| LSA3_11 | Los Santos | Tonosí | 7.40917 | -80.43694 | 2018 | 0 | 1 |
| LSA3_12 | Los Santos | Tonosí | 7.44333 | -80.42806 | 2018 | 0 | 1 |
| LSA3_13 | Los Santos | Tonosí | 7.41000 | -80.44361 | 2018 | 0 | 1 |
| LSA3_14 | Los Santos | Tonosí | 7.40861 | -80.44083 | 2018 | 0 | 1 |
| LSA3_15 | Los Santos | Tonosí | 7.40917 | -80.43917 | 2018 | 0 | 1 |
| LSA4_1 | Los Santos | El Cacao | 7.44056 | -80.41194 | 2018 | 0 | 0 |
| LSA4_2 | Los Santos | El Cacao | 7.43444 | -80.35389 | 2018 | 0 | 0 |
| LSA4_3 | Los Santos | El Cacao | 7.44389 | -80.41028 | 2018 | 0 | 0 |
| LSA4_4 | Los Santos | El Cacao | 7.45806 | -80.37806 | 2018 | 0 | 1 |

| | | | | | | | |
|---|---|---|---|---|---|---|---|
| LSA4_5 | Los Santos | El Cacao | 7.44861 | -80.40611 | 2018 | 0 | 1 |
| LSA4_6 | Los Santos | El Cacao | 7.44639 | -80.40472 | 2018 | 0 | 1 |
| LSA4_7 | Los Santos | El Cacao | 7.45667 | -80.39778 | 2018 | 0 | 0 |
| LSA4_8 | Los Santos | El Cacao | 7.45222 | -80.39417 | 2018 | 0 | 0 |
| LSA4_9 | Los Santos | El Cacao | 7.44889 | -80.40361 | 2018 | 0 | 0 |
| LSA4_10 | Los Santos | El Cacao | 7.45806 | -80.37806 | 2018 | 0 | 1 |
| LSA4_11 | Los Santos | El Cacao | 7.45111 | -80.40417 | 2018 | 0 | 1 |
| LSA4_12 | Los Santos | El Cacao | 7.45194 | -80.40222 | 2018 | 0 | 1 |
| LSA4_13 | Los Santos | El Cacao | 7.45278 | -80.40028 | 2018 | 0 | 0 |
| LSA4_14 | Los Santos | El Cacao | 7.45083 | -80.40056 | 2018 | 0 | 0 |
| LSA4_15 | Los Santos | El Cacao | 7.44944 | -80.40028 | 2018 | 0 | 0 |
| LSA5_1 | Los Santos | Pedasí | 7.53333 | -80.01944 | 2018 | 0 | 1 |
| LSA5_2 | Los Santos | Pedasí | 7.53417 | -80.02917 | 2018 | 1 | 1 |
| LSA5_3 | Los Santos | Pedasí | 7.53417 | -80.03056 | 2018 | 0 | 1 |
| LSA5_4 | Los Santos | Pedasí | 7.53278 | -80.02972 | 2018 | 0 | 1 |
| LSA5_5 | Los Santos | Pedasí | 7.53417 | -80.02917 | 2018 | 0 | 1 |
| LSA5_6 | Los Santos | Pedasí | 7.53000 | -80.01694 | 2018 | 0 | 1 |
| LSA5_7 | Los Santos | Pedasí | 7.52389 | -80.02694 | 2018 | 1 | 0 |
| LSA5_8 | Los Santos | Pedasí | 7.52528 | -80.02611 | 2018 | 0 | 1 |
| LSA5_9 | Los Santos | Pedasí | 7.52778 | -80.02222 | 2018 | 0 | 1 |
| LSA5_10 | Los Santos | Pedasí | 7.52833 | -80.02111 | 2018 | 0 | 1 |
| LSA5_11 | Los Santos | Pedasí | 7.53083 | -80.02472 | 2018 | 0 | 0 |
| LSA5_12 | Los Santos | Pedasí | 7.53111 | -80.02472 | 2018 | 0 | 1 |
| LSA5_13 | Los Santos | Pedasí | 7.53361 | -80.02639 | 2018 | 0 | 1 |
| LSA5_14 | Los Santos | Pedasí | 7.53278 | -80.02556 | 2018 | 0 | 0 |
| LSA5_15 | Los Santos | Pedasí | 7.53028 | -80.02500 | 2018 | 0 | 1 |
| LSA6_1 | Los Santos | Guararé | 7.75306 | -80.25028 | 2018 | 0 | 1 |
| LSA6_2 | Los Santos | Guararé | 7.81583 | -80.27556 | 2018 | 0 | 1 |
| LSA6_3 | Los Santos | Guararé | 7.81806 | -80.27306 | 2018 | 0 | 1 |
| LSA6_4 | Los Santos | Guararé | 7.81444 | -80.28111 | 2018 | 0 | 1 |
| LSA6_5 | Los Santos | Guararé | 7.81972 | -80.27500 | 2018 | 0 | 1 |
| LSA6_6 | Los Santos | Guararé | 7.81917 | -80.27750 | 2018 | 0 | 1 |
| LSA6_7 | Los Santos | Guararé | 7.81528 | -80.28083 | 2018 | 1 | 1 |
| LSA6_10 | Los Santos | Guararé | 7.81639 | -80.28306 | 2018 | 1 | 0 |
| LSA6_11 | Los Santos | Guararé | 7.81500 | -80.28139 | 2018 | 0 | 1 |
| LSA6_12 | Los Santos | Guararé | 7.81583 | -80.28111 | 2018 | 0 | 1 |
| LSA6_13 | Los Santos | Guararé | 7.83111 | -80.27972 | 2018 | 0 | 0 |
| LSA6_14 | Los Santos | Guararé | 7.81972 | -80.27778 | 2018 | 1 | 1 |
| LSA6_15 | Los Santos | Guararé | 7.82278 | -80.27194 | 2018 | 0 | 0 |
| LSA1_1 | Los Santos | La Villa | 7.93639 | -80.41611 | 2018 | 1 | 0 |
| LSA1_2 | Los Santos | La Villa | 7.94694 | -80.42056 | 2018 | 0 | 0 |
| LSA1_3 | Los Santos | La Villa | 7.93944 | -80.41333 | 2018 | 1 | 0 |
| LSA1_4 | Los Santos | La Villa | 7.93944 | -80.41083 | 2018 | 1 | 1 |

| | | | | | | | |
|---|---|---|---|---|---|---|---|
| LSA1_5 | Los Santos | La Villa | 7.93833 | -80.41194 | 2018 | 1 | 0 |
| LSA1_6 | Los Santos | La Villa | 7.93750 | -80.41139 | 2018 | 1 | 0 |
| LSA1_7 | Los Santos | La Villa | 7.93639 | -80.41194 | 2018 | 0 | 0 |
| LSA1_8 | Los Santos | La Villa | 7.93444 | -80.40861 | 2018 | 0 | 1 |
| LSA1_9 | Los Santos | La Villa | 7.92889 | -80.40556 | 2018 | 1 | 1 |
| LSA1_10 | Los Santos | La Villa | 7.93000 | -80.40722 | 2018 | 1 | 1 |
| LSA1_11 | Los Santos | La Villa | 7.92722 | -80.41028 | 2018 | 1 | 1 |
| LSA1_12 | Los Santos | La Villa | 7.93056 | -80.40889 | 2018 | 0 | 0 |
| LSA1_14 | Los Santos | La Villa | 7.93250 | -80.41333 | 2018 | 1 | 0 |
| LSA1_15 | Los Santos | La Villa | 7.93278 | -80.41528 | 2018 | 0 | 1 |
| LSA3_1 | Los Santos | Tonosí | 7.41167 | -80.45083 | 2018 | 0 | 1 |
| LSA3_2 | Los Santos | Tonosí | 7.41167 | -80.45083 | 2018 | 0 | 0 |
| LSA3_3 | Los Santos | Tonosí | 7.40861 | -80.44944 | 2018 | 0 | 1 |
| LSA3_4 | Los Santos | Tonosí | 7.40806 | -80.44611 | 2018 | 0 | 1 |
| LSA3_5 | Los Santos | Tonosí | 7.40750 | -80.44167 | 2018 | 0 | 1 |
| LSA3_6 | Los Santos | Tonosí | 7.40861 | -80.43333 | 2018 | 0 | 1 |
| LSA3_7 | Los Santos | Tonosí | 7.41167 | -80.43750 | 2018 | 0 | 1 |
| LSA3_8 | Los Santos | Tonosí | 7.41528 | -80.43667 | 2018 | 0 | 1 |
| LSA3_9 | Los Santos | Tonosí | 7.41500 | -80.43694 | 2018 | 0 | 1 |
| LSA3_10 | Los Santos | Tonosí | 7.41750 | -80.43806 | 2018 | 0 | 1 |
| LSA3_11 | Los Santos | Tonosí | 7.41444 | -80.43889 | 2018 | 0 | 0 |
| LSA3_12 | Los Santos | Tonosí | 7.41139 | -80.44028 | 2018 | 0 | 1 |
| LSA3_13 | Los Santos | Tonosí | 7.41139 | -80.44028 | 2018 | 0 | 1 |
| LSA3_14 | Los Santos | Tonosí | 7.42000 | -80.45583 | 2018 | 0 | 1 |
| LSA3_15 | Los Santos | Tonosí | 7.41583 | -80.44417 | 2018 | 0 | 1 |
| LSA4_1 | Los Santos | El Cacao | 7.44528 | -80.42000 | 2018 | 0 | 0 |
| LSA4_2 | Los Santos | El Cacao | 7.44833 | -80.41861 | 2018 | 0 | 0 |
| LSA4_3 | Los Santos | El Cacao | 7.45000 | -80.41806 | 2018 | 0 | 0 |
| LSA4_4 | Los Santos | El Cacao | 7.45361 | -80.41500 | 2018 | 0 | 0 |
| LSA4_5 | Los Santos | El Cacao | 7.45361 | -80.41500 | 2018 | 0 | 1 |
| LSA4_6 | Los Santos | El Cacao | 7.45556 | -80.40750 | 2018 | 0 | 1 |
| LSA4_7 | Los Santos | El Cacao | 7.45611 | -80.40528 | 2018 | 0 | 1 |
| LSA4_8 | Los Santos | El Cacao | 7.45611 | -80.40528 | 2018 | 0 | 0 |
| LSA4_9 | Los Santos | El Cacao | 7.45861 | -80.40778 | 2018 | 0 | 0 |
| LSA4_10 | Los Santos | El Cacao | 7.45000 | -80.40611 | 2018 | 0 | 1 |
| LSA4_11 | Los Santos | El Cacao | 7.45333 | -80.40417 | 2018 | 0 | 1 |
| LSA4_12 | Los Santos | El Cacao | 7.45500 | -80.40000 | 2018 | 0 | 1 |
| LSA4_13 | Los Santos | El Cacao | 7.45000 | -80.40000 | 2018 | 0 | 1 |
| LSA4_14 | Los Santos | El Cacao | 7.46028 | -80.40000 | 2018 | 0 | 1 |
| LSA4_15 | Los Santos | El Cacao | 7.46083 | -80.40722 | 2018 | 0 | 1 |
| LSA5_1 | Los Santos | Pedasí | 7.53333 | -80.01861 | 2018 | 0 | 1 |
| LSA5_2 | Los Santos | Pedasí | 7.53361 | -80.02111 | 2018 | 0 | 0 |
| LSA5_3 | Los Santos | Pedasí | 7.53361 | -80.02194 | 2018 | 0 | 0 |

| | | | | | | | |
|---|---|---|---|---|---|---|---|
| LSA5_4 | Los Santos | Pedasí | 7.53333 | -80.02306 | 2018 | 0 | 1 |
| LSA5_5 | Los Santos | Pedasí | 7.54361 | -80.02194 | 2018 | 1 | 1 |
| LSA5_6 | Los Santos | Pedasí | 7.54028 | -80.02139 | 2018 | 0 | 1 |
| LSA5_7 | Los Santos | Pedasí | 7.53889 | -80.02444 | 2018 | 1 | 1 |
| LSA5_8 | Los Santos | Pedasí | 7.53528 | -80.02083 | 2018 | 0 | 0 |
| LSA5_9 | Los Santos | Pedasí | 7.53333 | -80.01667 | 2018 | 1 | 1 |
| LSA5_10 | Los Santos | Pedasí | 7.53361 | -80.01194 | 2018 | 0 | 0 |
| LSA5_11 | Los Santos | Pedasí | 7.53528 | -80.00972 | 2018 | 0 | 0 |
| LSA5_12 | Los Santos | Pedasí | 7.53556 | -80.00972 | 2018 | 0 | 0 |
| LSA5_13 | Los Santos | Pedasí | 7.54028 | -80.01361 | 2018 | 0 | 1 |
| LSA5_14 | Los Santos | Pedasí | 7.54028 | -80.01361 | 2018 | 0 | 1 |
| LSA5_15 | Los Santos | Pedasí | 7.54333 | -80.01556 | 2018 | 0 | 1 |
| LSA6_1 | Los Santos | Guararé | 7.82472 | -80.28806 | 2018 | 0 | 1 |
| LSA6_2 | Los Santos | Guararé | 7.82611 | -80.28056 | 2018 | 0 | 1 |
| LSA6_3 | Los Santos | Guararé | 7.81694 | -80.27722 | 2018 | 0 | 1 |
| LSA6_4 | Los Santos | Guararé | 7.82306 | -80.27694 | 2018 | 0 | 1 |
| LSA6_5 | Los Santos | Guararé | 7.82417 | -80.28028 | 2018 | 0 | 1 |
| LSA6_6 | Los Santos | Guararé | 7.82444 | -80.28694 | 2018 | 1 | 1 |
| LSA6_7 | Los Santos | Guararé | 7.82167 | -80.28083 | 2018 | 1 | 1 |
| LSA6_8 | Los Santos | Guararé | 7.82194 | -80.28528 | 2018 | 1 | 1 |
| LSA6_9 | Los Santos | Guararé | 7.82167 | -80.28528 | 2018 | 0 | 1 |
| LSA6_10 | Los Santos | Guararé | 7.82500 | -80.29194 | 2018 | 0 | 1 |
| LSA6_11 | Los Santos | Guararé | 7.81694 | -80.29167 | 2018 | 1 | 0 |
| LSA6_12 | Los Santos | Guararé | 7.81667 | -80.29056 | 2018 | 0 | 0 |
| LSA6_13 | Los Santos | Guararé | 7.81694 | -80.28833 | 2018 | 1 | 1 |
| LSA6_14 | Los Santos | Guararé | 7.82611 | -80.28944 | 2018 | 0 | 1 |
| LSA6_15 | Los Santos | Guararé | 7.83167 | -80.28889 | 2018 | 0 | 1 |
| LSA1_1 | Los Santos | La Villa | 7.94139 | -80.41694 | 2018 | 1 | 0 |
| LSA1_2 | Los Santos | La Villa | 7.94556 | -80.41667 | 2018 | 1 | 0 |
| LSA1_3 | Los Santos | La Villa | 7.94361 | -80.42222 | 2018 | 1 | 1 |
| LSA1_4 | Los Santos | La Villa | 7.94556 | -80.41778 | 2018 | 0 | 0 |
| LSA1_5 | Los Santos | La Villa | 7.94167 | -80.41944 | 2018 | 0 | 0 |
| LSA1_6 | Los Santos | La Villa | 7.94111 | -80.41833 | 2018 | 1 | 1 |
| LSA1_7 | Los Santos | La Villa | 7.94111 | -80.41861 | 2018 | 1 | 0 |
| LSA1_8 | Los Santos | La Villa | 7.94083 | -80.41833 | 2018 | 1 | 1 |
| LSA1_9 | Los Santos | La Villa | 7.93750 | -80.40944 | 2018 | 1 | 1 |
| LSA1_10 | Los Santos | La Villa | 7.93444 | -80.41056 | 2018 | 0 | 1 |
| LSA1_11 | Los Santos | La Villa | 7.93444 | -80.40028 | 2018 | 1 | 1 |
| LSA1_12 | Los Santos | La Villa | 7.94056 | -80.42056 | 2018 | 1 | 1 |
| LSA1_13 | Los Santos | La Villa | 7.94083 | -80.42194 | 2018 | 1 | 1 |
| LSA1_14 | Los Santos | La Villa | 7.94194 | -80.42667 | 2018 | 0 | 1 |
| LSA1_15 | Los Santos | La Villa | 7.94250 | -80.41694 | 2018 | 1 | 0 |
| LSA3_1 | Los Santos | Tonosí | 7.41167 | -80.45083 | 2018 | 0 | 0 |

| | | | | | | | |
|---|---|---|---|---|---|---|---|
| LSA3_2 | Los Santos | Tonosí | 7.41167 | -80.45083 | 2018 | 0 | 1 |
| LSA3_3 | Los Santos | Tonosí | 7.40861 | -80.44944 | 2018 | 0 | 0 |
| LSA3_4 | Los Santos | Tonosí | 7.40806 | -80.44611 | 2018 | 0 | 1 |
| LSA3_5 | Los Santos | Tonosí | 7.40750 | -80.44167 | 2018 | 0 | 1 |
| LSA3_6 | Los Santos | Tonosí | 7.40861 | -80.43333 | 2018 | 0 | 0 |
| LSA3_7 | Los Santos | Tonosí | 7.41167 | -80.43750 | 2018 | 0 | 1 |
| LSA3_8 | Los Santos | Tonosí | 7.41528 | -80.43667 | 2018 | 0 | 1 |
| LSA3_9 | Los Santos | Tonosí | 7.41500 | -80.43694 | 2018 | 0 | 0 |
| LSA3_10 | Los Santos | Tonosí | 7.41750 | -80.43806 | 2018 | 0 | 1 |
| LSA3_11 | Los Santos | Tonosí | 7.41444 | -80.43889 | 2018 | 0 | 0 |
| LSA3_12 | Los Santos | Tonosí | 7.41139 | -80.44028 | 2018 | 0 | 1 |
| LSA3_13 | Los Santos | Tonosí | 7.41139 | -80.44028 | 2018 | 0 | 1 |
| LSA3_14 | Los Santos | Tonosí | 7.42000 | -80.45583 | 2018 | 0 | 1 |
| LSA3_15 | Los Santos | Tonosí | 7.41583 | -80.44417 | 2018 | 0 | 1 |
| LSA4_1 | Los Santos | El Cacao | 7.44528 | -80.42000 | 2018 | 0 | 1 |
| LSA4_2 | Los Santos | El Cacao | 7.44833 | -80.41861 | 2018 | 0 | 0 |
| LSA4_3 | Los Santos | El Cacao | 7.45000 | -80.41806 | 2018 | 0 | 0 |
| LSA4_4 | Los Santos | El Cacao | 7.45361 | -80.41500 | 2018 | 0 | 1 |
| LSA4_5 | Los Santos | El Cacao | 7.45361 | -80.41500 | 2018 | 0 | 1 |
| LSA4_6 | Los Santos | El Cacao | 7.45556 | -80.40750 | 2018 | 0 | 1 |
| LSA4_7 | Los Santos | El Cacao | 7.45611 | -80.40528 | 2018 | 0 | 1 |
| LSA4_8 | Los Santos | El Cacao | 7.45611 | -80.40528 | 2018 | 0 | 1 |
| LSA4_9 | Los Santos | El Cacao | 7.45861 | -80.40778 | 2018 | 0 | 0 |
| LSA4_10 | Los Santos | El Cacao | 7.45000 | -80.40611 | 2018 | 0 | 0 |
| LSA4_11 | Los Santos | El Cacao | 7.45333 | -80.40417 | 2018 | 0 | 1 |
| LSA4_12 | Los Santos | El Cacao | 7.45500 | -80.40000 | 2018 | 0 | 1 |
| LSA4_13 | Los Santos | El Cacao | 7.45000 | -80.40000 | 2018 | 0 | 1 |
| LSA4_14 | Los Santos | El Cacao | 7.46028 | -80.40000 | 2018 | 0 | 0 |
| LSA4_15 | Los Santos | El Cacao | 7.46083 | -80.40722 | 2018 | 0 | 1 |
| LSA5_1 | Los Santos | Pedasí | 7.53333 | -80.01861 | 2018 | 1 | 1 |
| LSA5_2 | Los Santos | Pedasí | 7.53361 | -80.02111 | 2018 | 0 | 1 |
| LSA5_3 | Los Santos | Pedasí | 7.53361 | -80.02194 | 2018 | 0 | 0 |
| LSA5_4 | Los Santos | Pedasí | 7.53333 | -80.02306 | 2018 | 1 | 1 |
| LSA5_5 | Los Santos | Pedasí | 7.54361 | -80.02194 | 2018 | 0 | 1 |
| LSA5_6 | Los Santos | Pedasí | 7.54028 | -80.02139 | 2018 | 1 | 1 |
| LSA5_7 | Los Santos | Pedasí | 7.53889 | -80.02444 | 2018 | 0 | 1 |
| LSA5_8 | Los Santos | Pedasí | 7.53528 | -80.02083 | 2018 | 1 | 1 |
| LSA5_9 | Los Santos | Pedasí | 7.53333 | -80.01667 | 2018 | 0 | 1 |
| LSA5_10 | Los Santos | Pedasí | 7.53361 | -80.01194 | 2018 | 1 | 1 |
| LSA5_11 | Los Santos | Pedasí | 7.53528 | -80.00972 | 2018 | 1 | 1 |
| LSA5_12 | Los Santos | Pedasí | 7.53556 | -80.00972 | 2018 | 1 | 1 |
| LSA5_13 | Los Santos | Pedasí | 7.54028 | -80.01361 | 2018 | 1 | 1 |
| LSA5_14 | Los Santos | Pedasí | 7.54028 | -80.01361 | 2018 | 0 | 1 |

| | | | | | | | |
|---|---|---|---|---|---|---|---|
| LSA5_15 | Los Santos | Pedasí | 7.54333 | -80.01556 | 2018 | 0 | 1 |
| LSA6_1 | Los Santos | Guararé | 7.82472 | -80.28806 | 2018 | 0 | 1 |
| LSA6_2 | Los Santos | Guararé | 7.82611 | -80.28056 | 2018 | 0 | 1 |
| LSA6_3 | Los Santos | Guararé | 7.81694 | -80.27722 | 2018 | 0 | 1 |
| LSA6_4 | Los Santos | Guararé | 7.82306 | -80.27694 | 2018 | 0 | 1 |
| LSA6_5 | Los Santos | Guararé | 7.82417 | -80.28028 | 2018 | 0 | 1 |
| LSA6_6 | Los Santos | Guararé | 7.82444 | -80.28694 | 2018 | 1 | 1 |
| LSA6_7 | Los Santos | Guararé | 7.82167 | -80.28083 | 2018 | 1 | 1 |
| LSA6_8 | Los Santos | Guararé | 7.82194 | -80.28528 | 2018 | 1 | 1 |
| LSA6_9 | Los Santos | Guararé | 7.82167 | -80.28528 | 2018 | 0 | 0 |
| LSA6_10 | Los Santos | Guararé | 7.82500 | -80.29194 | 2018 | 0 | 0 |
| LSA6_11 | Los Santos | Guararé | 7.81694 | -80.29167 | 2018 | 1 | 1 |
| LSA6_12 | Los Santos | Guararé | 7.81667 | -80.29056 | 2018 | 1 | 1 |
| LSA6_13 | Los Santos | Guararé | 7.81694 | -80.28833 | 2018 | 0 | 0 |
| LSA6_14 | Los Santos | Guararé | 7.82611 | -80.28944 | 2018 | 1 | 1 |
| LSA6_15 | Los Santos | Guararé | 7.83167 | -80.28889 | 2018 | 0 | 1 |
| LSA1_1 | Los Santos | La Villa | 7.94139 | -80.41694 | 2018 | 1 | 1 |
| LSA1_2 | Los Santos | La Villa | 7.94556 | -80.41667 | 2018 | 1 | 1 |
| LSA1_3 | Los Santos | La Villa | 7.94361 | -80.42222 | 2018 | 1 | 0 |
| LSA1_4 | Los Santos | La Villa | 7.94556 | -80.41778 | 2018 | 1 | 0 |
| LSA1_5 | Los Santos | La Villa | 7.94167 | -80.41944 | 2018 | 0 | 1 |
| LSA1_6 | Los Santos | La Villa | 7.94111 | -80.41833 | 2018 | 0 | 0 |
| LSA1_7 | Los Santos | La Villa | 7.94111 | -80.41861 | 2018 | 1 | 1 |
| LSA1_8 | Los Santos | La Villa | 7.94083 | -80.41833 | 2018 | 1 | 0 |
| LSA1_9 | Los Santos | La Villa | 7.93750 | -80.40944 | 2018 | 1 | 1 |
| LSA1_10 | Los Santos | La Villa | 7.93444 | -80.41056 | 2018 | 0 | 1 |
| LSA1_11 | Los Santos | La Villa | 7.93444 | -80.40028 | 2018 | 1 | 1 |
| LSA1_12 | Los Santos | La Villa | 7.94056 | -80.42056 | 2018 | 0 | 1 |
| LSA1_13 | Los Santos | La Villa | 7.94083 | -80.42194 | 2018 | 1 | 1 |
| LSA1_14 | Los Santos | La Villa | 7.94194 | -80.42667 | 2018 | 1 | 1 |
| LSA1_15 | Los Santos | La Villa | 7.94250 | -80.41694 | 2018 | 0 | 1 |
| LSA3_1 | Los Santos | Tonosí | 7.41167 | -80.45083 | 2018 | 0 | 1 |
| LSA3_2 | Los Santos | Tonosí | 7.41167 | -80.45083 | 2018 | 0 | 1 |
| LSA3_3 | Los Santos | Tonosí | 7.40861 | -80.44944 | 2018 | 0 | 0 |
| LSA3_4 | Los Santos | Tonosí | 7.40806 | -80.44611 | 2018 | 0 | 0 |
| LSA3_5 | Los Santos | Tonosí | 7.40750 | -80.44167 | 2018 | 0 | 1 |
| LSA3_6 | Los Santos | Tonosí | 7.40861 | -80.43333 | 2018 | 0 | 1 |
| LSA3_7 | Los Santos | Tonosí | 7.41167 | -80.43750 | 2018 | 0 | 1 |
| LSA3_8 | Los Santos | Tonosí | 7.41528 | -80.43667 | 2018 | 0 | 1 |
| LSA3_9 | Los Santos | Tonosí | 7.41500 | -80.43694 | 2018 | 0 | 1 |
| LSA3_10 | Los Santos | Tonosí | 7.41750 | -80.43806 | 2018 | 0 | 1 |
| LSA3_11 | Los Santos | Tonosí | 7.41444 | -80.43889 | 2018 | 0 | 1 |
| LSA3_12 | Los Santos | Tonosí | 7.41139 | -80.44028 | 2018 | 0 | 1 |

| | | | | | | | |
|---|---|---|---|---|---|---|---|
| LSA3_13 | Los Santos | Tonosí | 7.41139 | -80.44028 | 2018 | 0 | 0 |
| LSA3_14 | Los Santos | Tonosí | 7.42000 | -80.45583 | 2018 | 0 | 1 |
| LSA3_15 | Los Santos | Tonosí | 7.41583 | -80.44417 | 2018 | 0 | 1 |
| LSA4_1 | Los Santos | El Cacao | 7.44528 | -80.42000 | 2018 | 0 | 1 |
| LSA4_2 | Los Santos | El Cacao | 7.44833 | -80.41861 | 2018 | 0 | 1 |
| LSA4_3 | Los Santos | El Cacao | 7.45000 | -80.41806 | 2018 | 0 | 0 |
| LSA4_4 | Los Santos | El Cacao | 7.45361 | -80.41500 | 2018 | 0 | 1 |
| LSA4_5 | Los Santos | El Cacao | 7.45361 | -80.41500 | 2018 | 0 | 1 |
| LSA4_6 | Los Santos | El Cacao | 7.45556 | -80.40750 | 2018 | 0 | 0 |
| LSA4_7 | Los Santos | El Cacao | 7.45611 | -80.40528 | 2018 | 0 | 0 |
| LSA4_8 | Los Santos | El Cacao | 7.45611 | -80.40528 | 2018 | 0 | 1 |
| LSA4_9 | Los Santos | El Cacao | 7.45861 | -80.40778 | 2018 | 0 | 1 |
| LSA4_10 | Los Santos | El Cacao | 7.45000 | -80.40611 | 2018 | 0 | 1 |
| LSA4_11 | Los Santos | El Cacao | 7.45333 | -80.40417 | 2018 | 0 | 1 |
| LSA4_12 | Los Santos | El Cacao | 7.45500 | -80.40000 | 2018 | 0 | 1 |
| LSA4_13 | Los Santos | El Cacao | 7.45000 | -80.40000 | 2018 | 0 | 1 |
| LSA4_14 | Los Santos | El Cacao | 7.46028 | -80.40000 | 2018 | 0 | 1 |
| LSA4_15 | Los Santos | El Cacao | 7.46083 | -80.40722 | 2018 | 0 | 1 |
| LSA5_1 | Los Santos | Pedasí | 7.53333 | -80.01861 | 2018 | 1 | 1 |
| LSA5_2 | Los Santos | Pedasí | 7.53361 | -80.02111 | 2018 | 0 | 1 |
| LSA5_3 | Los Santos | Pedasí | 7.53361 | -80.02194 | 2018 | 0 | 1 |
| LSA5_4 | Los Santos | Pedasí | 7.53333 | -80.02306 | 2018 | 0 | 0 |
| LSA5_5 | Los Santos | Pedasí | 7.54361 | -80.02194 | 2018 | 0 | 1 |
| LSA5_6 | Los Santos | Pedasí | 7.54028 | -80.02139 | 2018 | 0 | 1 |
| LSA5_7 | Los Santos | Pedasí | 7.53889 | -80.02444 | 2018 | 1 | 1 |
| LSA5_8 | Los Santos | Pedasí | 7.53528 | -80.02083 | 2018 | 0 | 1 |
| LSA5_9 | Los Santos | Pedasí | 7.53333 | -80.01667 | 2018 | 0 | 1 |
| LSA5_10 | Los Santos | Pedasí | 7.53361 | -80.01194 | 2018 | 1 | 1 |
| LSA5_11 | Los Santos | Pedasí | 7.53528 | -80.00972 | 2018 | 0 | 1 |
| LSA5_12 | Los Santos | Pedasí | 7.53556 | -80.00972 | 2018 | 0 | 1 |
| LSA5_13 | Los Santos | Pedasí | 7.54028 | -80.01361 | 2018 | 0 | 1 |
| LSA5_14 | Los Santos | Pedasí | 7.54028 | -80.01361 | 2018 | 0 | 1 |
| LSA5_15 | Los Santos | Pedasí | 7.54333 | -80.01556 | 2018 | 0 | 1 |
| LSA6_1 | Los Santos | Guararé | 7.82472 | -80.28806 | 2018 | 1 | 1 |
| LSA6_2 | Los Santos | Guararé | 7.82611 | -80.28056 | 2018 | 0 | 1 |
| LSA6_3 | Los Santos | Guararé | 7.81694 | -80.27722 | 2018 | 0 | 1 |
| LSA6_4 | Los Santos | Guararé | 7.82306 | -80.27694 | 2018 | 0 | 1 |
| LSA6_5 | Los Santos | Guararé | 7.82417 | -80.28028 | 2018 | 0 | 1 |
| LSA6_6 | Los Santos | Guararé | 7.82444 | -80.28694 | 2018 | 0 | 1 |
| LSA6_7 | Los Santos | Guararé | 7.82167 | -80.28083 | 2018 | 0 | 1 |
| LSA6_8 | Los Santos | Guararé | 7.82194 | -80.28528 | 2018 | 1 | 0 |
| LSA6_9 | Los Santos | Guararé | 7.82167 | -80.28528 | 2018 | 1 | 1 |
| LSA6_10 | Los Santos | Guararé | 7.82500 | -80.29194 | 2018 | 1 | 1 |

| | | | | | | | |
|---|---|---|---|---|---|---|---|
| LSA6_11 | Los Santos | Guararé | 7.81694 | -80.29167 | 2018 | 0 | 0 |
| LSA6_12 | Los Santos | Guararé | 7.81667 | -80.29056 | 2018 | 0 | 1 |
| LSA6_13 | Los Santos | Guararé | 7.81694 | -80.28833 | 2018 | 1 | 1 |
| LSA6_14 | Los Santos | Guararé | 7.82611 | -80.28944 | 2018 | 0 | 1 |
| LSA6_15 | Los Santos | Guararé | 7.83167 | -80.28889 | 2018 | 0 | 0 |
| LSA1_1 | Los Santos | La Villa | 7.94139 | -80.41694 | 2018 | 0 | 0 |
| LSA1_2 | Los Santos | La Villa | 7.94556 | -80.41667 | 2018 | 1 | 1 |
| LSA1_3 | Los Santos | La Villa | 7.94361 | -80.42222 | 2018 | 0 | 0 |
| LSA1_4 | Los Santos | La Villa | 7.94556 | -80.41778 | 2018 | 0 | 0 |
| LSA1_5 | Los Santos | La Villa | 7.94167 | -80.41944 | 2018 | 0 | 0 |
| LSA1_6 | Los Santos | La Villa | 7.94111 | -80.41833 | 2018 | 0 | 1 |
| LSA1_7 | Los Santos | La Villa | 7.94111 | -80.41861 | 2018 | 0 | 0 |
| LSA1_8 | Los Santos | La Villa | 7.94083 | -80.41833 | 2018 | 0 | 0 |
| LSA1_9 | Los Santos | La Villa | 7.93750 | -80.40944 | 2018 | 0 | 0 |
| LSA1_10 | Los Santos | La Villa | 7.93444 | -80.41056 | 2018 | 0 | 1 |
| LSA1_11 | Los Santos | La Villa | 7.93444 | -80.40028 | 2018 | 0 | 0 |
| LSA1_12 | Los Santos | La Villa | 7.94056 | -80.42056 | 2018 | 1 | 1 |
| LSA1_13 | Los Santos | La Villa | 7.94083 | -80.42194 | 2018 | 1 | 1 |
| LSA1_14 | Los Santos | La Villa | 7.94194 | -80.42667 | 2018 | 1 | 1 |
| LSA1_15 | Los Santos | La Villa | 7.94250 | -80.41694 | 2018 | 1 | 1 |
| LSA3_1 | Los Santos | Tonosí | 7.41167 | -80.45083 | 2018 | 0 | 1 |
| LSA3_2 | Los Santos | Tonosí | 7.41167 | -80.45083 | 2018 | 0 | 1 |
| LSA3_3 | Los Santos | Tonosí | 7.40861 | -80.44944 | 2018 | 0 | 0 |
| LSA3_4 | Los Santos | Tonosí | 7.40806 | -80.44611 | 2018 | 0 | 1 |
| LSA3_5 | Los Santos | Tonosí | 7.40750 | -80.44167 | 2018 | 0 | 1 |
| LSA3_6 | Los Santos | Tonosí | 7.40861 | -80.43333 | 2018 | 0 | 0 |
| LSA3_7 | Los Santos | Tonosí | 7.41167 | -80.43750 | 2018 | 0 | 1 |
| LSA3_8 | Los Santos | Tonosí | 7.41528 | -80.43667 | 2018 | 0 | 0 |
| LSA3_9 | Los Santos | Tonosí | 7.41500 | -80.43694 | 2018 | 0 | 0 |
| LSA3_10 | Los Santos | Tonosí | 7.41750 | -80.43806 | 2018 | 0 | 0 |
| LSA3_11 | Los Santos | Tonosí | 7.41444 | -80.43889 | 2018 | 0 | 1 |
| LSA3_12 | Los Santos | Tonosí | 7.41139 | -80.44028 | 2018 | 0 | 1 |
| LSA3_13 | Los Santos | Tonosí | 7.41139 | -80.44028 | 2018 | 0 | 0 |
| LSA3_14 | Los Santos | Tonosí | 7.42000 | -80.45583 | 2018 | 0 | 0 |
| LSA3_15 | Los Santos | Tonosí | 7.41583 | -80.44417 | 2018 | 0 | 0 |
| LSA4_1 | Los Santos | El Cacao | 7.44528 | -80.42000 | 2018 | 0 | 0 |
| LSA4_2 | Los Santos | El Cacao | 7.44833 | -80.41861 | 2018 | 0 | 0 |
| LSA4_3 | Los Santos | El Cacao | 7.45000 | -80.41806 | 2018 | 0 | 0 |
| LSA4_4 | Los Santos | El Cacao | 7.45361 | -80.41500 | 2018 | 0 | 0 |
| LSA4_5 | Los Santos | El Cacao | 7.45361 | -80.41500 | 2018 | 0 | 0 |
| LSA4_6 | Los Santos | El Cacao | 7.45556 | -80.40750 | 2018 | 0 | 1 |
| LSA4_7 | Los Santos | El Cacao | 7.45611 | -80.40528 | 2018 | 0 | 0 |
| LSA4_8 | Los Santos | El Cacao | 7.45611 | -80.40528 | 2018 | 0 | 1 |

| | | | | | | | |
|---|---|---|---|---|---|---|---|
| LSA4_9 | Los Santos | El Cacao | 7.45861 | -80.40778 | 2018 | 0 | 0 |
| LSA4_10 | Los Santos | El Cacao | 7.45000 | -80.40611 | 2018 | 0 | 1 |
| LSA4_11 | Los Santos | El Cacao | 7.45333 | -80.40417 | 2018 | 0 | 0 |
| LSA4_12 | Los Santos | El Cacao | 7.45500 | -80.40000 | 2018 | 0 | 1 |
| LSA4_13 | Los Santos | El Cacao | 7.45000 | -80.40000 | 2018 | 0 | 1 |
| LSA4_14 | Los Santos | El Cacao | 7.46028 | -80.40000 | 2018 | 0 | 1 |
| LSA4_15 | Los Santos | El Cacao | 7.46083 | -80.40722 | 2018 | 0 | 0 |
| LSA5_1 | Los Santos | Pedasí | 7.53333 | -80.01861 | 2018 | 0 | 1 |
| LSA5_2 | Los Santos | Pedasí | 7.53361 | -80.02111 | 2018 | 0 | 1 |
| LSA5_3 | Los Santos | Pedasí | 7.53361 | -80.02194 | 2018 | 0 | 1 |
| LSA5_4 | Los Santos | Pedasí | 7.53333 | -80.02306 | 2018 | 0 | 1 |
| LSA5_5 | Los Santos | Pedasí | 7.54361 | -80.02194 | 2018 | 1 | 0 |
| LSA5_6 | Los Santos | Pedasí | 7.54028 | -80.02139 | 2018 | 0 | 1 |
| LSA5_7 | Los Santos | Pedasí | 7.53889 | -80.02444 | 2018 | 0 | 0 |
| LSA5_8 | Los Santos | Pedasí | 7.53528 | -80.02083 | 2018 | 0 | 0 |
| LSA5_9 | Los Santos | Pedasí | 7.53333 | -80.01667 | 2018 | 0 | 0 |
| LSA5_10 | Los Santos | Pedasí | 7.53361 | -80.01194 | 2018 | 0 | 1 |
| LSA5_11 | Los Santos | Pedasí | 7.53528 | -80.00972 | 2018 | 0 | 0 |
| LSA5_12 | Los Santos | Pedasí | 7.53556 | -80.00972 | 2018 | 0 | 0 |
| LSA5_13 | Los Santos | Pedasí | 7.54028 | -80.01361 | 2018 | 0 | 1 |
| LSA5_14 | Los Santos | Pedasí | 7.54028 | -80.01361 | 2018 | 0 | 0 |
| LSA5_15 | Los Santos | Pedasí | 7.54333 | -80.01556 | 2018 | 0 | 0 |
| LSA6_1 | Los Santos | Guararé | 7.84139 | -80.27861 | 2018 | 0 | 0 |
| LSA6_2 | Los Santos | Guararé | 7.84028 | -80.27639 | 2018 | 0 | 1 |
| LSA6_3 | Los Santos | Guararé | 7.83333 | -80.27500 | 2018 | 0 | 1 |
| LSA6_4 | Los Santos | Guararé | 7.83833 | -80.27667 | 2018 | 0 | 1 |
| LSA6_5 | Los Santos | Guararé | 7.82917 | -80.27667 | 2018 | 0 | 1 |
| LSA6_6 | Los Santos | Guararé | 7.82861 | -80.29139 | 2018 | 0 | 1 |
| LSA6_7 | Los Santos | Guararé | 7.83194 | -80.29222 | 2018 | 0 | 1 |
| LSA6_8 | Los Santos | Guararé | 7.83028 | -80.28778 | 2018 | 0 | 1 |
| LSA6_9 | Los Santos | Guararé | 7.82694 | -80.28361 | 2018 | 0 | 1 |
| LSA6_10 | Los Santos | Guararé | 7.82167 | -80.29361 | 2018 | 0 | 0 |
| LSA6_11 | Los Santos | Guararé | 7.81667 | -80.29444 | 2018 | 0 | 1 |
| LSA6_12 | Los Santos | Guararé | 7.81694 | -80.28972 | 2018 | 1 | 1 |
| LSA6_13 | Los Santos | Guararé | 7.82750 | -80.28944 | 2018 | 0 | 0 |
| LSA6_14 | Los Santos | Guararé | 7.81667 | -80.28778 | 2018 | 0 | 0 |
| LSA6_15 | Los Santos | Guararé | 7.81694 | -80.28333 | 2018 | 0 | 1 |
| LSA1_1 | Los Santos | La Villa | 7.94139 | -80.40528 | 2018 | 0 | 0 |
| LSA1_2 | Los Santos | La Villa | 7.93833 | -80.40639 | 2018 | 0 | 1 |
| LSA1_3 | Los Santos | La Villa | 7.94111 | -80.41083 | 2018 | 0 | 1 |
| LSA1_4 | Los Santos | La Villa | 7.93806 | -80.40028 | 2018 | 0 | 1 |
| LSA1_5 | Los Santos | La Villa | 7.93750 | -80.41028 | 2018 | 1 | 1 |
| LSA1_6 | Los Santos | La Villa | 7.94000 | -80.41306 | 2018 | 0 | 1 |

| | | | | | | | |
|---|---|---|---|---|---|---|---|
| LSA1_7 | Los Santos | La Villa | 7.95917 | -80.41139 | 2018 | 0 | 1 |
| LSA1_8 | Los Santos | La Villa | 7.93333 | -80.41361 | 2018 | 0 | 0 |
| LSA1_9 | Los Santos | La Villa | 7.93361 | -80.42444 | 2018 | 1 | 1 |
| LSA1_10 | Los Santos | La Villa | 7.93806 | -80.42056 | 2018 | 1 | 0 |
| LSA1_11 | Los Santos | La Villa | 7.93639 | -80.42500 | 2018 | 0 | 1 |
| LSA1_12 | Los Santos | La Villa | 7.93333 | -80.41694 | 2018 | 0 | 1 |
| LSA1_13 | Los Santos | La Villa | 7.93361 | -80.42222 | 2018 | 0 | 0 |
| LSA1_14 | Los Santos | La Villa | 7.93361 | -80.42417 | 2018 | 0 | 1 |
| LSA1_15 | Los Santos | La Villa | 7.94139 | -80.42333 | 2018 | 1 | 0 |
| LSA3_1 | Los Santos | Tonosí | 7.41167 | -80.45083 | 2018 | 0 | 1 |
| LSA3_2 | Los Santos | Tonosí | 7.41167 | -80.45083 | 2018 | 0 | 1 |
| LSA3_3 | Los Santos | Tonosí | 7.40861 | -80.44944 | 2018 | 0 | 1 |
| LSA3_4 | Los Santos | Tonosí | 7.40806 | -80.44611 | 2018 | 0 | 1 |
| LSA3_5 | Los Santos | Tonosí | 7.40750 | -80.44167 | 2018 | 0 | 0 |
| LSA3_6 | Los Santos | Tonosí | 7.40861 | -80.43333 | 2018 | 0 | 0 |
| LSA3_7 | Los Santos | Tonosí | 7.41167 | -80.43750 | 2018 | 0 | 1 |
| LSA3_8 | Los Santos | Tonosí | 7.41528 | -80.43667 | 2018 | 0 | 0 |
| LSA3_9 | Los Santos | Tonosí | 7.41500 | -80.43694 | 2018 | 0 | 0 |
| LSA3_10 | Los Santos | Tonosí | 7.41750 | -80.43806 | 2018 | 0 | 1 |
| LSA3_11 | Los Santos | Tonosí | 7.41444 | -80.43889 | 2018 | 0 | 1 |
| LSA3_12 | Los Santos | Tonosí | 7.41139 | -80.44028 | 2018 | 0 | 0 |
| LSA3_13 | Los Santos | Tonosí | 7.41139 | -80.44028 | 2018 | 0 | 0 |
| LSA3_14 | Los Santos | Tonosí | 7.42000 | -80.45583 | 2018 | 0 | 1 |
| LSA3_15 | Los Santos | Tonosí | 7.41583 | -80.44417 | 2018 | 0 | 0 |
| LSA4_1 | Los Santos | El Cacao | 7.44528 | -80.42000 | 2018 | 0 | 0 |
| LSA4_2 | Los Santos | El Cacao | 7.44833 | -80.41861 | 2018 | 0 | 0 |
| LSA4_3 | Los Santos | El Cacao | 7.45000 | -80.41806 | 2018 | 0 | 0 |
| LSA4_4 | Los Santos | El Cacao | 7.45361 | -80.41500 | 2018 | 0 | 1 |
| LSA4_5 | Los Santos | El Cacao | 7.45361 | -80.41500 | 2018 | 0 | 1 |
| LSA4_6 | Los Santos | El Cacao | 7.45556 | -80.40750 | 2018 | 0 | 0 |
| LSA4_7 | Los Santos | El Cacao | 7.45611 | -80.40528 | 2018 | 0 | 0 |
| LSA4_8 | Los Santos | El Cacao | 7.45611 | -80.40528 | 2018 | 0 | 1 |
| LSA4_9 | Los Santos | El Cacao | 7.45861 | -80.40778 | 2018 | 0 | 1 |
| LSA4_10 | Los Santos | El Cacao | 7.45000 | -80.40611 | 2018 | 0 | 0 |
| LSA4_11 | Los Santos | El Cacao | 7.45333 | -80.40417 | 2018 | 0 | 0 |
| LSA4_12 | Los Santos | El Cacao | 7.45500 | -80.40000 | 2018 | 0 | 0 |
| LSA4_13 | Los Santos | El Cacao | 7.45000 | -80.40000 | 2018 | 0 | 0 |
| LSA4_14 | Los Santos | El Cacao | 7.46028 | -80.40000 | 2018 | 0 | 0 |
| LSA4_15 | Los Santos | El Cacao | 7.46083 | -80.40722 | 2018 | 0 | 0 |
| LSA5_1 | Los Santos | Pedasí | 7.53333 | -80.01861 | 2018 | 0 | 0 |
| LSA5_2 | Los Santos | Pedasí | 7.53361 | -80.02111 | 2018 | 0 | 1 |
| LSA5_3 | Los Santos | Pedasí | 7.53361 | -80.02194 | 2018 | 0 | 1 |
| LSA5_4 | Los Santos | Pedasí | 7.53333 | -80.02306 | 2018 | 0 | 1 |

| | | | | | | | |
|---|---|---|---|---|---|---|---|
| LSA5_5 | Los Santos | Pedasí | 7.54361 | -80.02194 | 2018 | 1 | 1 |
| LSA5_6 | Los Santos | Pedasí | 7.54028 | -80.02139 | 2018 | 0 | 0 |
| LSA5_7 | Los Santos | Pedasí | 7.53889 | -80.02444 | 2018 | 0 | 0 |
| LSA5_8 | Los Santos | Pedasí | 7.53528 | -80.02083 | 2018 | 0 | 1 |
| LSA5_9 | Los Santos | Pedasí | 7.53333 | -80.01667 | 2018 | 0 | 1 |
| LSA5_10 | Los Santos | Pedasí | 7.53361 | -80.01194 | 2018 | 1 | 0 |
| LSA5_11 | Los Santos | Pedasí | 7.53528 | -80.00972 | 2018 | 0 | 0 |
| LSA5_12 | Los Santos | Pedasí | 7.53556 | -80.00972 | 2018 | 0 | 1 |
| LSA5_13 | Los Santos | Pedasí | 7.54028 | -80.01361 | 2018 | 0 | 1 |
| LSA5_14 | Los Santos | Pedasí | 7.54028 | -80.01361 | 2018 | 0 | 1 |
| LSA5_15 | Los Santos | Pedasí | 7.54333 | -80.01556 | 2018 | 0 | 1 |
| LSA6_1 | Los Santos | Guararé | 7.84139 | -80.27861 | 2018 | 0 | 0 |
| LSA6_2 | Los Santos | Guararé | 7.84028 | -80.27639 | 2018 | 0 | 0 |
| LSA6_3 | Los Santos | Guararé | 7.83333 | -80.27500 | 2018 | 0 | 1 |
| LSA6_4 | Los Santos | Guararé | 7.83833 | -80.27667 | 2018 | 0 | 1 |
| LSA6_5 | Los Santos | Guararé | 7.82917 | -80.27667 | 2018 | 0 | 1 |
| LSA6_6 | Los Santos | Guararé | 7.82861 | -80.29139 | 2018 | 0 | 0 |
| LSA6_7 | Los Santos | Guararé | 7.83194 | -80.29222 | 2018 | 0 | 0 |
| LSA6_8 | Los Santos | Guararé | 7.83028 | -80.28778 | 2018 | 0 | 0 |
| LSA6_9 | Los Santos | Guararé | 7.82694 | -80.28361 | 2018 | 0 | 1 |
| LSA6_10 | Los Santos | Guararé | 7.82167 | -80.29361 | 2018 | 1 | 1 |
| LSA6_11 | Los Santos | Guararé | 7.81667 | -80.29444 | 2018 | 0 | 1 |
| LSA6_12 | Los Santos | Guararé | 7.81694 | -80.28972 | 2018 | 0 | 1 |
| LSA6_13 | Los Santos | Guararé | 7.82750 | -80.28944 | 2018 | 0 | 1 |
| LSA6_14 | Los Santos | Guararé | 7.81667 | -80.28778 | 2018 | 0 | 0 |
| LSA6_15 | Los Santos | Guararé | 7.81694 | -80.28333 | 2018 | 0 | 1 |
| LSA1_1 | Los Santos | La Villa | 7.94139 | -80.40528 | 2018 | 0 | 0 |
| LSA1_2 | Los Santos | La Villa | 7.93833 | -80.40639 | 2018 | 0 | 0 |
| LSA1_3 | Los Santos | La Villa | 7.94111 | -80.41083 | 2018 | 0 | 0 |
| LSA1_4 | Los Santos | La Villa | 7.93806 | -80.40028 | 2018 | 0 | 0 |
| LSA1_5 | Los Santos | La Villa | 7.93750 | -80.41028 | 2018 | 0 | 1 |
| LSA1_6 | Los Santos | La Villa | 7.94000 | -80.41306 | 2018 | 0 | 1 |
| LSA1_7 | Los Santos | La Villa | 7.95917 | -80.41139 | 2018 | 0 | 1 |
| LSA1_8 | Los Santos | La Villa | 7.93333 | -80.41361 | 2018 | 1 | 1 |
| LSA1_9 | Los Santos | La Villa | 7.93361 | -80.42444 | 2018 | 0 | 1 |
| LSA1_10 | Los Santos | La Villa | 7.93806 | -80.42056 | 2018 | 1 | 0 |
| LSA1_11 | Los Santos | La Villa | 7.93639 | -80.42500 | 2018 | 1 | 1 |
| LSA1_12 | Los Santos | La Villa | 7.93333 | -80.41694 | 2018 | 1 | 1 |
| LSA1_13 | Los Santos | La Villa | 7.93361 | -80.42222 | 2018 | 0 | 0 |
| LSA1_14 | Los Santos | La Villa | 7.93361 | -80.42417 | 2018 | 0 | 0 |
| LSA1_15 | Los Santos | La Villa | 7.94139 | -80.42333 | 2018 | 1 | 0 |

**S4 Table.** Correlation coefficients for the variables NDVI vegetation index, rainfall, the difference between minimum and maximum temperature (Temp. diff), humidity, human population density (HPopD), maximum temperature (MaxT) and minimum temperature (MinT) averaged across all months (below diagonal) and from the rainy season months of May to November (above diagonal).

| | NDVI | Rainfall | Temp. diff | Humidity | HPopD | MaxT | MinT |
|---|---|---|---|---|---|---|---|
| NDVI | 1.00 | -0.02 | -0.44 | 0.45 | 0.08 | -0.39 | -0.47 |
| Rainfall | -0.02 | 1.00 | -0.36 | 0.03 | -0.01 | -0.39 | -0.28 |
| Temp. diff | -0.44 | -0.36 | 1.00 | -0.76 | 0.13 | **0.82** | **0.94** |
| Humidity | 0.45 | 0.03 | -0.76 | 1.00 | 0.26 | -0.73 | -0.80 |
| HPopD | 0.08 | -0.01 | 0.13 | 0.26 | 1.00 | 0.06 | -0.05 |
| MaxT | -0.39 | -0.39 | **0.82** | -0.73 | 0.06 | 1.00 | 0.72 |
| MinT | -0.47 | -0.28 | **0.94** | -0.80 | -0.05 | 0.72 | 1.00 |
